# Genome-wide LD scores enable robust heritability and genetic correlation estimation from summary statistics across genetic ancestries

**DOI:** 10.64898/2026.10.20.760508

**Authors:** Moonseong Jeong, Zhengtong Liu, Jiayi Ni, Ulzee An, Sungjoo Han, Aakarsh Anand, Prateek Anand, Mayuko Boffelli, Sriram Sankararaman

## Abstract

Our knowledge of human complex trait architecture across diverse populations remains limited. Estimating SNP heritability and genetic correlation from GWAS summary statistics – successful in revealing genetic architecture in homogeneous cohorts – is challenging in structured and admixed populations due, in part, to the difficulty of representing long-range LD. We present SUMMIT, a statistical framework that combines genome-wide LD scores, efficiently estimated using random projections, with GWAS summary statistics to estimate genome-wide and annotation-specific heritability and genetic correlation. In simulations that explore diverse aspects of genetic architecture across European (EUR), South Asian (SAS), and admixed African-European cohorts (AFR) in the UK Biobank (UKB), SUMMIT had increased accuracy and reduced bias compared to prior methods, with the largest gains in non-European ancestries reflecting improved modeling of long-range LD. Across 40 UKB traits, SUMMIT produced estimates of genome-wide common-SNP heritability 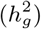 and heritability enrichments in genomic annotations that were broadly concordant across ancestries with the SUMMIT 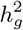 estimates more closely matching recent whole-genome sequencing based estimates. SUMMIT substantially improved genetic correlation analysis recovering more significant findings across ancestries (23 SAS and 3 AFR at FDR *<* 0.05 compared with none and two for cov-LDSC) with the non-European discoveries showing concordant effects in Europeans suggesting that, like heritability, global patterns of pleiotropy are largely shared across ancestries. By combining accurate genome-wide LD and efficient moment estimation, SUMMIT enables investigation into how heritability and genetic correlation vary across populations and genomic annotations, including in structured and admixed cohorts, at the genome-wide, annotation-specific, and cross-trait levels.

## Introduction

The rapid growth of biobanks has made it possible to study complex-trait genetics at a scale that was not feasible until recently. Beyond scale, there is increasing emphasis on building biobanks that represent genetic diversity across populations (Bycroft et al. 2018; All of Us Research Program Genomics Investigators 2024; Verma et al. 2024; Zhou et al. 2022; Abul-Husn and Kenny 2019). Despite this trend of biobank-scale datasets increasingly including heterogeneous, recently admixed, or structured populations, our understanding of genetic architecture beyond relatively homogeneous European ancestry populations remains limited, in part, due to the limitations of analytical methods for inferring parameters that govern architecture.

Heritability 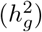 and genetic covariance/correlation (*r*_*g*_) are key parameters that describe complex-trait architecture, quantifying how much phenotypic variation is tagged by genotyped variants, and how genetic signal is distributed across genomic annotations, and shared across traits. Methods that use individual-level genotypes, including linear mixed models and scalable moment estimators, provide precise estimates of these parameters but require access to genotypes and phenotypes that can be difficult to obtain (Yang et al. 2011; Speed et al. 2012; Wu and Sankararaman 2018; Pazokitoroudi et al. 2020; Hou et al. 2019; Wu et al. 2022). This has made summary statistic methods a practical alternative for large-scale genetic analyses (Bulik-Sullivan et al. 2015b; Finucane et al. 2015; Speed and Balding 2019; Schwartzman et al. 2019; Ning et al. 2020; Lu et al. 2017; Shi et al. 2016, 2017; Zhang et al. 2021). Summary statistic methods, exemplified by LD score regression and its extensions, use as inputs GWAS statistics and a summary of the LD structure within the cohort *e*.*g*., the LD score captures the total LD between a SNP and all other SNPs in the genome (Bulik-Sullivan et al. 2015a,b; Finucane et al. 2015; Luo et al. 2021). While these methods tend to be less precise than individual-level methods, they circumvent the limitations of data access leading to their wide applicability. The reliability of these methods, however, depends critically on the accuracy of the LD scores. In practice, computational considerations lead to LD scores being estimated within a fixed window (usually a couple of Mb). This approximation has been shown to work well in relatively homogeneous populations (Bulik-Sullivan et al. 2015b). In structured or admixed populations, however, LD extends across substantial length scales *e*.*g*., admixture can generate long-range LD that can extend across chromosomes (Pfaff et al. 2001). Fixed-window LD scores are likely to miss the contribution of long-range LD which can lead to biased or unstable estimates; an approach to extend LDSC to admixed populations by regressing out genotype principal components (PCs) (Luo et al. 2021) does not fully account for long-range LD.

Here we present SUMMIT (Summary-statistics-based Unified Method for Multivariate Inference of Traits), a summary-statistics framework designed for estimating heritability and genetic covariance/correlation both genome-wide and within genomic annotations, using GWAS summary statistics and genome-wide LD information. SUMMIT is designed to provide accurate estimates across diverse genetic ancestries (as we show by applying it to GWAS in individuals of European ancestry) but is most effective in structured and admixed cohorts. In order to effectively capture long-range LD in structured and admixed populations, SUMMIT estimates LD scores across the genome without restricting to fixed-size windows. This is done in a scalable manner via random projections to estimate genome-wide LD scores – additionally adjusted for genotype PCs – giving unbiased estimate of total LD. These LD scores are then combined with GWAS summary statistics to reconstruct heritability and genetic covariance/correlation estimators (both genome-wide and stratified across genomic annotations). We show that this framework is *mathematically equivalent* to our previously proposed estimators that require individual-level data (Pazokitoroudi et al. 2020; Wu et al. 2022) leading to SUMMIT having high precision and reduced bias across genetic ancestries. Beyond genome-wide heritability and genetic covariance/correlation, SUMMIT also estimates these parameters stratified across genomic annotations, both overlapping and non-overlapping.

We performed extensive simulations to evaluate the bias and precision of SUMMIT across AFR (admixed African-European), SAS (South Asian), EUR, and EUR (300k) cohorts, where EUR is the 10k subset of EUR (300k) and EUR (300k) is the set of *≈* 300k unrelated white British individuals in the UK Biobank (UKB). We then apply SUMMIT to 40 UK Biobank traits across these ancestries to estimate genome-wide heritability, heritability enrichment in functional annotations, and genome-wide genetic correlations. Finally, we evaluate SUMMIT in a setting where in-sample LD is not available by analyzing GWAS summary statistics from external (non-UKB) studies with ancestry-matched LD scores computed from UKB. Across these analyses, SUMMIT improves the robustness, accuracy, and power of genetic architecture inference across ancestries, allowing us to examine how heritability and genetic correlation vary across populations and genomic annotations, including in structured and admixed cohorts, at the genome-wide, annotation-specific, and cross-trait levels.

## Results

### Overview of the SUMMIT framework

SUMMIT is a framework for estimating genetic architecture parameters, specifically heritability and genetic covariance/correlation both genome-wide and within genomic annotations, using GWAS summary statistics and genome-wide LD information. This framework has two steps. First, we estimate the genome-wide LD score for each SNP which measures the total squared correlation between that SNP and all the SNPs across the genome. Estimating LD scores genome-wide is computationally challenging, as it naively scales in *O* (*N*_LD_*M* ^2^) where *N*_LD_ is the number of individuals used for LD computation and *M* is the number of SNPs. SUMMIT efficiently estimates genome-wide LD scores by multiplying the genotype matrix by *B* random vectors and aggregating the resulting projections. This is a one-time computation for a given population and scales as *O*(*N*_LD_*MB*). Second, SUMMIT combines the genome-wide LD scores with GWAS summary statistics to estimate SNP heritability (SUM-RHE) and genetic covariance and genetic correlation (SUM-CORE). The same scheme can also be used to compute partitioned genome-wide LD scores with non-overlapping, overlapping, or continuous-valued SNP annotations, enabling estimation of annotation-specific heritability, genetic covariance, genetic correlation, and related annotation-level quantities. Required inputs and analysis options are summarized in Table 1.

**Table 1:** SUMMIT framework structure and required inputs. LD scores are computed once per population and annotation model. SUM-RHE and SUM-CORE then reuse these LD scores across traits. SUM-CORE has a default supplied-overlap mode, used in the primary analyses, and an ancillary summary-estimated-overlap mode used only in supplementary benchmarks. *τ* ^*\**^ is the standardized annotation effect, equivalent to the annotation-specific *τ* coefficient scaled by annotation variability and average per-SNP heritability, following Gazal et al. (2017). It measures the conditional relative change in per-SNP heritability associated with a one-standard-deviation increase in the annotation.

| Module | Estimand | Required inputs | Output |
| --- | --- | --- | --- |
| Genome-wide LD scores | Partitioned LD scores, $\tilde{\ell}_{jk}$ | LD genotype matrix $\mathbf{G}_{LD}$<br>LD covariates $\mathbf{C}_{LD}$<br>Annotation matrix $\mathbf{A}$ | LD-score file and blockwise LD aggregates |
| SUM-RHE | Genome-wide or partitioned SNP heritability, $h^2$ or $\{h_k^2\}$ | GWAS summary statistics $(\hat{\beta}_j, \widehat{SE}_j, n_j)$<br>Precomputed $\tilde{\ell}_{jk}$<br>Annotations (optional) | $\hat{h}^2, \hat{h}_k^2$, enrichment, $\tau^*$ |
| SUM-CORE, default | Genome-wide or annotation-specific genetic covariance/correlation, $\gamma_g, r_g, \{\gamma_k, r_{g,k}\}$ | Multiple GWAS summary statistics<br>Precomputed $\tilde{\ell}_{jk}$<br>Annotations (optional)<br>Supplied sample-overlap covariance $\hat{c}_{ov}$ on the HE scale (default) | $\hat{\gamma}_g, \hat{r}_g$, and optional annotation-specific $\{\hat{\gamma}_k, \hat{r}_{g,k}\}$ |

The key insight underlying the accuracy of parameter estimates in the SUMMIT framework is that these estimates are equivalent to randomized MoM estimators that we have previously developed and that have been shown to be highly accurate in populations of European ancestry (Pazokitoroudi et al. 2020; Wu et al. 2022) (see Methods for details). These previous methods require access to individual-level genotype data while SUMMIT works with GWAS summary statistics. Equally importantly, the use of genome-wide LD scores combined with the use of genotype PCs as covariates ensures that the SUMMIT framework remains accurate and robust when applied to structured and admixed populations.

We evaluated SUMMIT on UK Biobank cohorts spanning European (EUR), South Asian (SAS), and admixed African-European (AFR) ancestries. We first used simulations to assess the effectiveness of SUMMIT in estimating genome-wide and partitioned 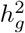 and genome-wide *r*_*g*_. We then applied SUMMIT to 40 complex traits that cover anthropometric, biomarker, behavioral, and disease phenotypes. These analyses were designed to evaluate SUMMIT across both large homogeneous cohorts and smaller, non-European or admixed cohorts where LD misspecification can be severe. Further, to evaluate its utility in a setting where in-sample LD information is not available, we applied SUMMIT to analyze non-UKB GWAS summary statistics using LD scores computed from UKB. Unless otherwise stated, cohort construction, trait definitions, GWAS procedures, variant QC, SNP-set differences across cohorts (Supplementary Fig. S1), and external-GWAS harmonization are described in Methods and Supplementary.

### Current approaches underestimate genome-wide LD in structured populations

We first estimated in-sample LD scores in each cohort to quantify how much genome-wide LD is captured by fixed-window and chromosome-wide summaries. We measured missed LD as the proportional difference between windowed or chromosome-wide LD scores and the SUMMIT genome-wide LD-score estimate (see Methods for details). Windowed LD scores tend to be substantially lower than genome-wide LD scores particularly in SAS and AFR cohorts, consistent with ancestry-driven long-range LD (Figure 1A, Supplementary Fig. S2 for the mean LD scores as a function of window size). Similarly, due to long-range LD, we find that chromosome-wide LD scores also tend to be lower than the genome-wide LD scores in these cohorts (Figure 1A, Supplementary Fig. S2). While regressing out PCs removes a large fraction of this long-range structure, windowed LD scores still underestimate LD in SAS and AFR relative to genome-wide estimates (Figure 1A, Supplementary Fig. S2). The underestimation is small in EUR (0.25%) but remains appreciable in SAS (8.7%) and AFR (13.4%), suggesting that genotype PCs alone do not fully remove long-range LD in structured cohorts, aligned with recent evidence that PC adjustment does not completely remove the effects of ancestry structure (Hu et al. 2025). Increasing the number of genotype PCs had minimal impact on recovering the remaining LD deficit (Supplementary Fig. S3), and genome-wide 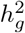 estimates for three representative traits were stable across these choices (Supplementary Fig. S4). These observations motivate the accurate estimation of genome-wide LD scores.

**Figure 1:**
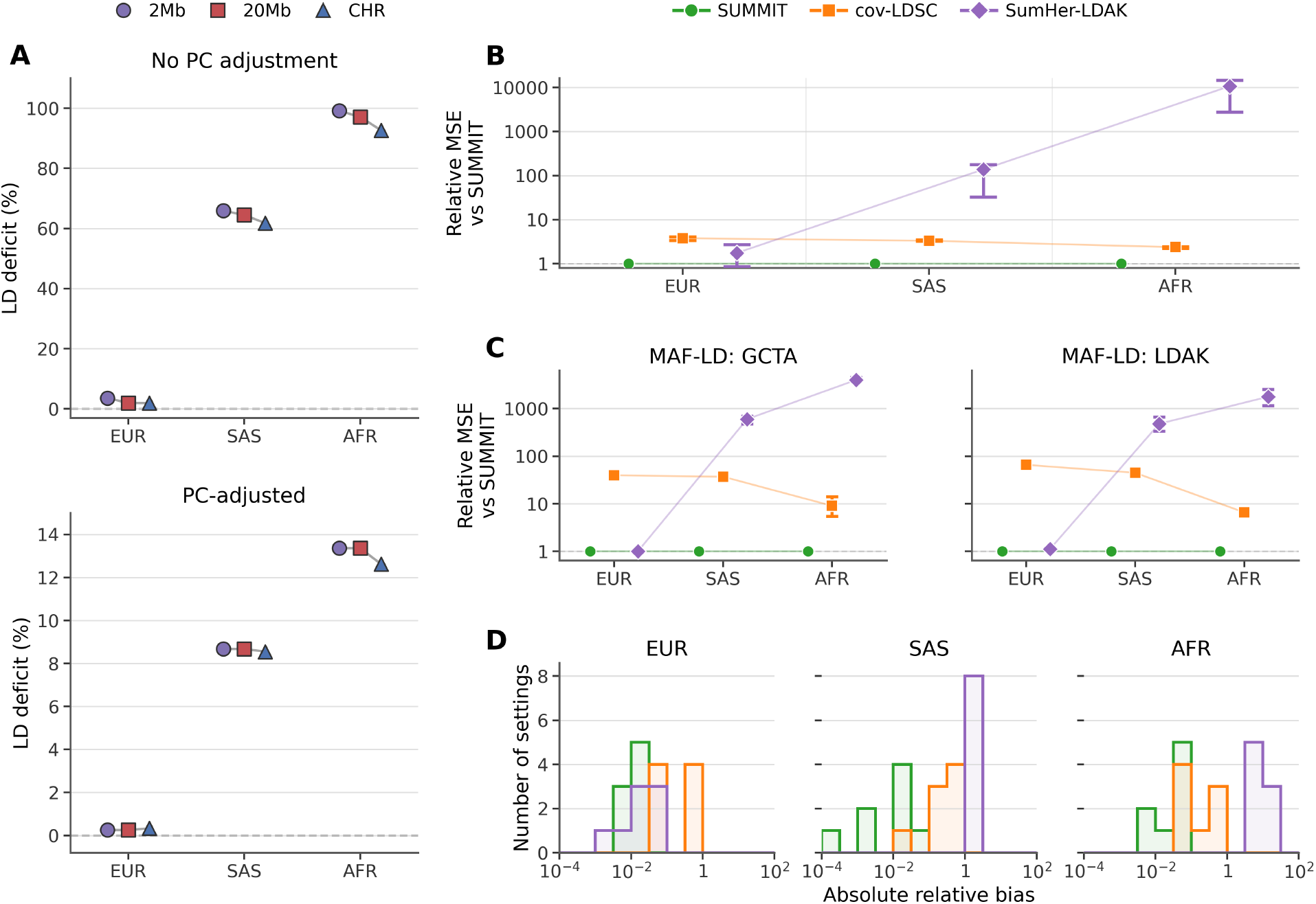
Average LD across UKB cohorts and genome-wide heritability simulation benchmarks for SUMMIT. **Panel A:** Each point shows the percentage of total LD not captured by a windowed or chromosome-wide estimate, defined as 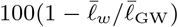, where 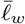 is the mean 2-Mb, 20-Mb, or chromosome-wide LD score and 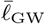 is the corresponding mean genome-wide LD score in the same UKB cohort. Genome-wide and chromosome-wide LD scores were each estimated with SUMMIT using *B* = 1,024 random vectors, and 2-Mb and 20-Mb LD scores were computed with LDSC. The top panel shows LD scores computed without genotype-PC adjustment, and the bottom panel shows LD scores computed after adjustment for the in-sample genotype PCs. **Panel B:** MSE of genome-wide 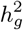 estimates relative to SUMMIT across the nine single-component settings defined by 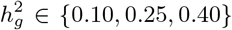 and *p*_causal_ ∈ {1.0, 0.10, 0.01}. A value of 1 denotes the same MSE as SUMMIT. Points show the median across settings, and bars show the interquartile range. **Panels C and D:** Results from MAF–LD simulations with 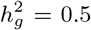. The eight settings comprised two effect-size architecures (GCTA and LDAK) with causal variants restricted to lower- and higher-MAF variants: MAF *<* 0.1 and MAF *≥* 0.1) and two causal proportions (*p*_causal_ = 1.0 and 0.01). Each method estimated heritability using the same 24-component MAF–LD model. **Panel C:** MSE of genome-wide 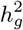 estimates relative to SUMMIT. A value of 1 denotes the same MSE as SUMMIT. Points show the median across the four settings within each simulation scheme and cohort, and bars show the interquartile range. **Panel D:** Within each setting, show the eight setting-level values for each method, separately by cohort; the horizontal axis is logarithmic. Full absolute relative bias was defined as 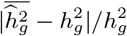, where the mean was calculated across 100 replicates. Histograms results are available in Supplementary Figs. S5–S13.

### Simulation benchmarks

The simulation benchmarks were designed to systematically explore the accuracy of SUMMIT both in homogeneous (EUR) and structured (SAS and AFR) cohorts. Based on our analyses in the previous section, we use the top 10 PCs in AFR and EUR and the top 20 in SAS. For our main results, we compare SUMMIT to cov-LDSC, the most directly comparable method, and SumHer-LDAK, an alternative method to model LD-dependent genetic architecture. For the LD summaries required by each method, we used the largest feasible window size (20 Mb for both LD scores and LDAK taggings; see “Benchmark methods” section of Methods for details). Exploration of sensitivity to window size and proportion of missing SNPs, comparison to additional methods, binary-trait simulations and simulations on EUR (300k), and additional accuracy metrics are shown in Supplementary Figures S5–S8.

### Genome-wide heritability

We first benchmarked genome-wide heritability estimation under nine genetic architectures per population (EUR, SAS, and AFR), varying 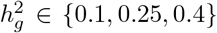 and causal proportion *p ∈ {*1.0, 0.1, 0.01*}* (details in Supplementary Note S1.9). In the main-text figure (Figure 1B–D), we compare SUMMIT to cov-LDSC and SumHer-LDAK. Full results including comparisons to LDSC, HDL, SumHer-GCTA, and RHE-mc are shown in the Supplementary Fig. S6 with details given in “Benchmark methods” section of Methods.

SUMMIT had the lowest mean squared error (MSE) across all methods in all three populations (Fig. 1B). In EUR, cov-LDSC had a 2.5–5.1-fold higher MSE than SUMMIT across the nine settings (median 3.8). This improved accuracy in EUR cannot be attributed primarily to missing long-range LD, because standard and genome-wide LD scores are nearly identical after PC residualization (Fig. 1A). Consistent with this interpretation, SUMMIT closely matched RHE-mc in EUR in the full benchmark (0.93–1.02-fold relative MSE; median 0.97; Supplementary Fig. S6), indicating that the SUMMIT estimator, relying on summary statistics, behaves equivalently to RHE-mc which uses individual-level data.

cov-LDSC, designed to be applied to admixed populations, had 2.3–4.0-fold higher MSE than SUMMIT in SAS (median 3.3) and 2.2–2.9-fold higher MSE in AFR (median 2.4). In these populations, LD score estimation is more challenging, and fixed-window LD scores underestimate genome-wide LD even after regressing out the PCs. We also find that RHE-mc (an individual-level method run without including PCs as covariates) was also less accurate than SUMMIT in these cohorts (1.3–48.1-fold higher MSE in SAS (median 8.9) and 1.5–33.8-fold higher MSE in AFR (median 9.6)). SUMMIT run without PC residualization obtained estimates that were close to RHE-mc but was less accurate than the default SUMMIT (Supplementary Fig. S9), suggesting that the improved accuracy of SUMMIT can be attributed to the use of PC residualization together with accurate genome-wide LD score estimation. The remaining summary-level methods that do not use PC residualization were substantially less stable in the structured cohorts (Supplementary Figs. S5–S6). Across LDSC, SumHer-GCTA, SumHer-LDAK and HDL, MSE was 10.9 to about 800-fold higher than SUMMIT in SAS and 26.3-fold to more than 10^4^-fold higher in AFR in the full benchmark; for HDL, many estimates were fixed at 1.0 because larger estimates were clipped.

The previous simulations highlight the effects of population structure on heritability estimation as they do not have stratified effects. We also simulated phenotypes with an environmental stratification component aligned with the first genotype PC (Supplementary Fig. S10; details in Supplementary Note S1.9). SUMMIT again had the lowest MSE and remained approximately unbiased, whereas cov-LDSC had 2.4–3.8-fold higher MSE. Methods run without covariates showed further inflation, especially in SAS and AFR.

We also examined the impact of missing SNPs on the heritability estimates by removing sets of 10%, 25%, or 50% of variants from the summary statistics with the LD scores estimated on the full SNP set. Across all nine single-component settings, the mean 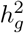 estimate decreased approximately in proportion to the fraction of SNPs retained (Supplementary Fig. S11; see Supplementary Note S1.9 for details).

Prior studies have shown the importance of accounting for the dependence of effect sizes on MAF and LD when estimating genome-wide 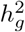 (Gazal et al. 2017; Speed and Balding 2019; Gazal et al. 2019). SUMMIT can model this dependence by estimating 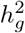 where SNPs are assigned to annotations based on their MAF and local LD. In simulations with MAF- and LD-dependent architectures (Yang et al. (2011); Speed et al. (2012); Speed and Balding (2019); see Supplementary Note S1.9 for simulation details and “Variant annotations” in Methods for details on MAF and LD annotations), SUMMIT had the lowest median MSE (Fig. 1C) and relative bias (Fig. 1D) with no significant bias across architectures, whereas cov-LDSC and SumHer-LDAK were biased (Supplementary Table S1). Full error summaries are shown in Supplementary Figs. S12 and S13.

### Heritability enrichment

Next, we evaluated SUMMIT for annotation-specific heritability and enrichment. We simulated traits in which protein-coding regions, defined as the Baseline-LD `Coding_UCSC` annotation and its 500-bp flanks, were enriched for heritability and compared them with matched null simulations where there is no enrichment in these regions (Supplementary Note S1.9). We evaluated the ability of SUMMIT to distinguish these two cases. Discrimination was evaluated using the coding-enrichment statistic 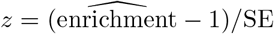 to distinguish enriched simulations from null simulations with the same total heritability and causal proportion (with balanced class prevalence), where we additionally varied enrichment magnitudes and effect-size architectures. SUMMIT was most accurate in distinguishing enriched from null simulations with highest area under the ROC (AUROC) and PR (AUPR) curves (Fig. 2A; Supplementary Fig. S14) while also having controlled false positive rates (FPR) in all three ancestries (Fig. 2B; Supplementary Fig. S15).

**Figure 2:**
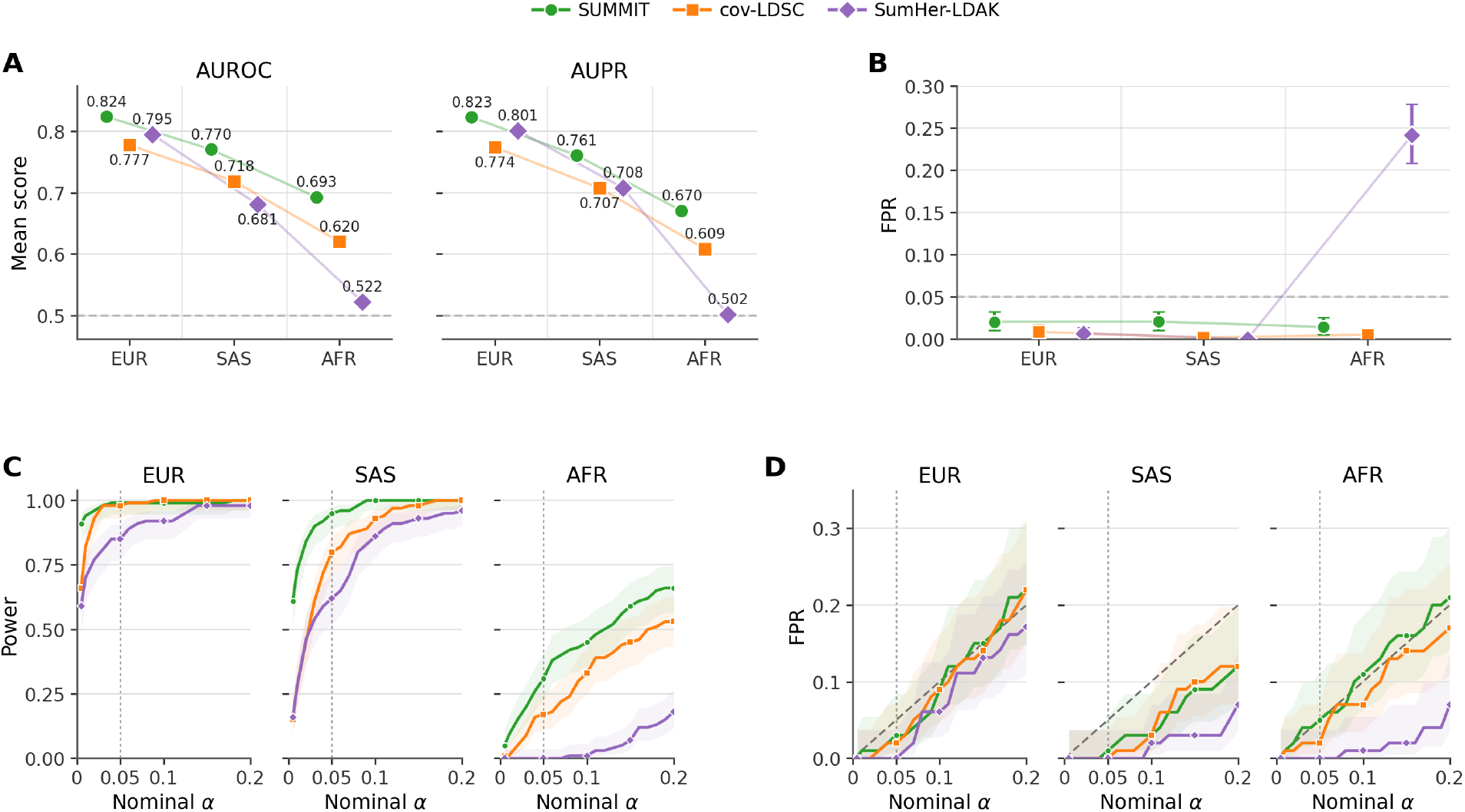
Simulation benchmarks for heritability enrichment with disjoint (A,B) and overlapping (C,D) annotations. **Panel A:** Area under ROC (AUROC) and PR (AUPR) curves on the task of distinguishing protein coding annotations that are enriched for heritability from those that are not enriched (see Supplementary Note S1.9 for details). **Panel B:** False-positive rate (FPR at *α* = 0.05) for the one-sided test of enrichment in coding regions. **Panel C:** Power and **Panel D:** FPR for detecting enrichment of the focal promoter-age annotation in a model with overlapping annotations as a function of nominal *α*. Error bars in panel B and shaded regions in Panels C and D show pointwise 95% Wilson confidence intervals. The dashed line in Panel A marks random classification, the dashed line in Panel B marks the nominal FPR, and the diagonal dashed lines in Panel D mark equality between the observed FPR and nominal *α*. Vertical dotted lines mark *α* = 0.05.

We then evaluated the accuracy of SUMMIT to detect enrichment in a model with overlapping annotations. We considered sixteen components: a focal ancient-promoter annotation, 14 regulatory and gene-model annotations, and a base annotation (Supplementary Table S2). All simulations had genome-wide 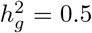. Under the null, all heritability was assigned to the base component; under the alternative, heritability of 0.05 was assigned to the focal annotation with the rest assigned to the base component. Across ancestries, SUMMIT had the highest power to detect enrichment in the focal annotation (Fig. 2C) while having calibrated FPR at *α* = 0.05 (Fig. 2D) with substantial gains in the non-European ancestries: SUMMIT shows an increase in power of 19% in SAS and 82% in AFR over cov-LDSC (although the power of all methods tends to be reduced in AFR compared to EUR and SAS). These results support the use of SUMMIT to estimate heritability enrichment both with non-overlapping and overlapping annotations across ancestries.

### Genetic correlation

We next evaluated estimation of genome-wide genetic-correlation. We simulated architectures with true null genome-wide *r*_*g*_ (*r*_*g*_ = 0: `Nullrg`), ones where the genetic correlations was constant across MAF-LD bins (*r*_*g*_ is identical across bins: `Constrg`), and where genetic correlations varied across bins (*r*_*g*_ is higher in higher LD bins: `Gradrg`) architectures (Supplementary Note S1.9). In null *r*_*g*_ architecture, SUMMIT remained calibrated in all cohorts (Fig. 3A) with more precise estimates (median reported cov-LDSC SEs were 7.2, 5.5 and 2.5 times those of SUMMIT in EUR, SAS and AFR, respectively). Across these architectures, SUMMIT accurately estimated both genome-wide and bin-level *r*_*g*_ whereas cov-LDSC showed larger dispersion, particularly in low-MAF and low-LD bins (Supplementary Fig. S16). We observed that when the true *r*_*g*_ varied across LD quartiles (`Gradrg` architecture), methods that estimate genome-wide *r*_*g*_ using a single-component could be biased, most prominently in EUR (300k) (Fig. 3B and Supplementary Fig. S17). This observation motivates our use of MAF–LD-partitioned model for genome-wide *r*_*g*_ estimation while exploring sensitivity to this choice in real data analyses.

**Figure 3:**
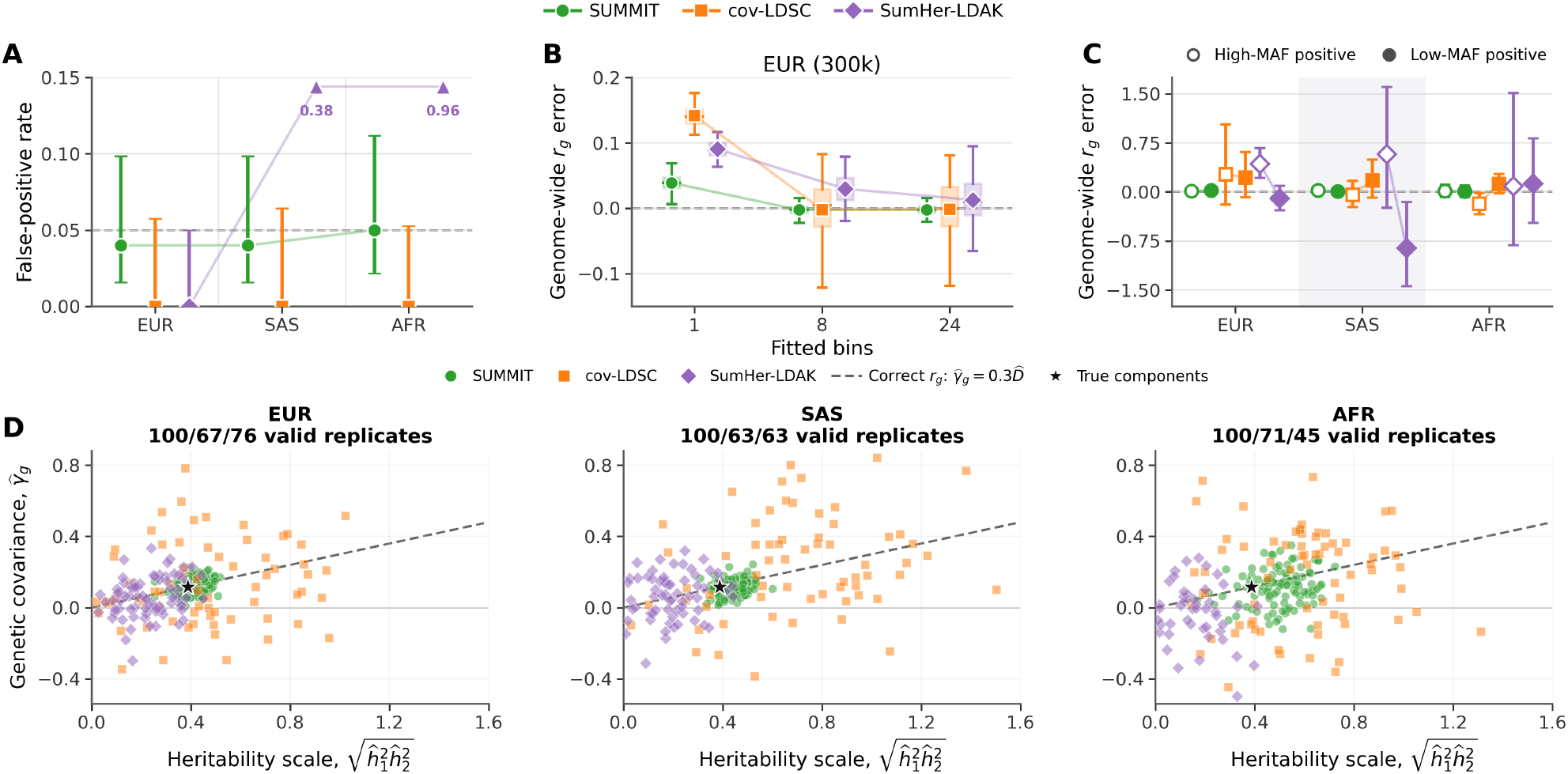
Simulation benchmarks for genome-wide genetic covariance and genetic correlation. **Panel A:** Empirical rejection rate for the two-sided test of genome-wide *r*_*g*_ = 0 (*α* = 0.05). Error bars show 95% confidence intervals and the dashed line marks the nominal level. **Panel B:** Error in genome-wide *r*_*g*_ estimates in EUR (300k) under a genetic architecture where *r*_*g*_ varies with LD bins (termed Gradrg). For each method, we assess the impact of fitting a single component (1 bin) and multiple MAF-LD stratified components (8 or 24 bins). Boxes show the interquartile range, center lines show medians, symbols show means, and whiskers show the 2.5th and 97.5th percentiles across 100 replicates. **Panel C:** Genome-wide *r*_*g*_ error under signed genetic correlation architectures with identical genome-wide *r*_*g*_, but differed in whether the high-MAF or the low-MAF bins had positive *r*_*g*_ while the remaining MAF bins had negative *r*_*g*_. Open (filled) symbols denote the architecture with positive genetic correlation in high (low) MAF and negative genetic correlation in low (high) MAF respectively. Points show means and error bars show 95% replicate-bootstrap confidence intervals. **Panel D:** Decomposition of the genome-wide *r*_*g*_ estimate into estimated genetic covariance, 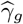 and heritability, 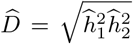, in the `Gradrg` architecture. Each point is one valid replicate where 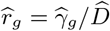. The dashed line is fixed at the true relationship, 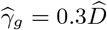 (not fitted to the points) and therefore contains estimates with the correct genome-wide *r*_*g*_ while the black star marks true values of genetic covariance and product of heritabilities. Replicates were retained when genetic covariance and both heritability estimates were finite and the estimated heritability product was positive. All such replicates are shown without filtering on magnitude. Valid replicate counts for SUMMIT, cov-LDSC, and SumHer-LDAK were 100*/*67*/*76 in EUR, 100*/*63*/*63 in SAS, and 100*/*71*/*45 in AFR.

We also simulated two architectures in which bin-specific genetic covariance and *r*_*g*_ changed sign between low- and high-MAF bins, while total genetic covariance and genome-wide *r*_*g*_ = 0.30 were held fixed (MAF cutoff of 0.1; Supplementary Note S1.9). One architecture assigned positive covariance to low-MAF bins and negative covariance to high-MAF bins; the other reversed these signs. These MAF–LD dependent genetic correlation architectures were motivated by prior evidence that effect-size vary with MAF and LD under negative selection (Gazal et al. 2017; Zeng et al. 2018; Schoech et al. 2019) and by our observation that, analogously, some trait pairs show MAF-dependent *r*_*g*_ in EUR (300k) (Supplementary Fig. S18). Supplied-overlap and summary-estimated-overlap modes produced broadly consistent point estimates, with greater variability in the latter (Supplementary Fig. S19). Compared to cov-LDSC and SumHer-LDAK, SUMMIT genome-wide *r*_*g*_ estimates had lower errors (centered around zero) under these architectures in all three ancestries (Fig. 3C).

An accurate *r*_*g*_ estimate can result from compensating errors in genetic covariance and heritability (since *r*_*g*_ = *γ*_*g*_*/D* where 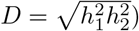, but this cannot be robust when the errors do not “cancel out”. We therefore examined the two components of *r*_*g*_ separately. While SUMMIT estimates remained concentrated near the true covariance and heritability scale, SumHer-LDAK in EUR and SAS, and cov-LDSC in SAS, often shifted away from the true components but approximately along the dashed line (Figure 3D), so that errors in genetic covariance and heritability partially cancelled. This cancellation failed when the two components shifted by incompatible amounts, most clearly for SumHer-LDAK in AFR.

### Runtime measurements

The main computational cost in SUMMIT is genome-wide LD-score estimation, which is performed once for each reference panel, SNP set and annotation model, and then reused across traits. We used *B* = 1,024 random vectors for the single-component simulation benchmarks in all four cohorts, and higher for multi-component settings (*≥* 10,000). Mean and SNP-level genome-wide LD-score estimates converged as *B* increased, and downstream genome-wide 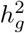 estimates were stable over the evaluated range (Supplementary Figs. S20–S22). These estimates were stable relative to high-*B* reference calculations (Supplementary Fig. S22). This step took much less time with similar memory usage when compared to windowed LD scores despite estimating genome-wide LD (Supplementary Table S3,S4), as well as linearly scaling to sample size *N*, number of variants *M*, and number of bins *K* (Supplementary Fig. S23). After LD scores and GWAS summary statistics were available, SUMMIT heritability estimation took seconds per trait and was similar to or faster than cov-LDSC and SumHer-LDAK in the single-component benchmarks.

### Application to UK Biobank summary statistics from 40 traits

#### SUMMIT accurately estimates SNP heritability across populations

Applying SUMMIT to 40 UK Biobank traits, we found that total common-SNP 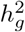 estimates were broadly stable across cohorts for the same trait (Fig. 4A; all 40 traits in the Supplementary Fig. S24; see “Benchmark methods” section in Methods for additional details). To quantify the variation in these estimates across cohorts for method *m*, we computed

**Figure 4:**
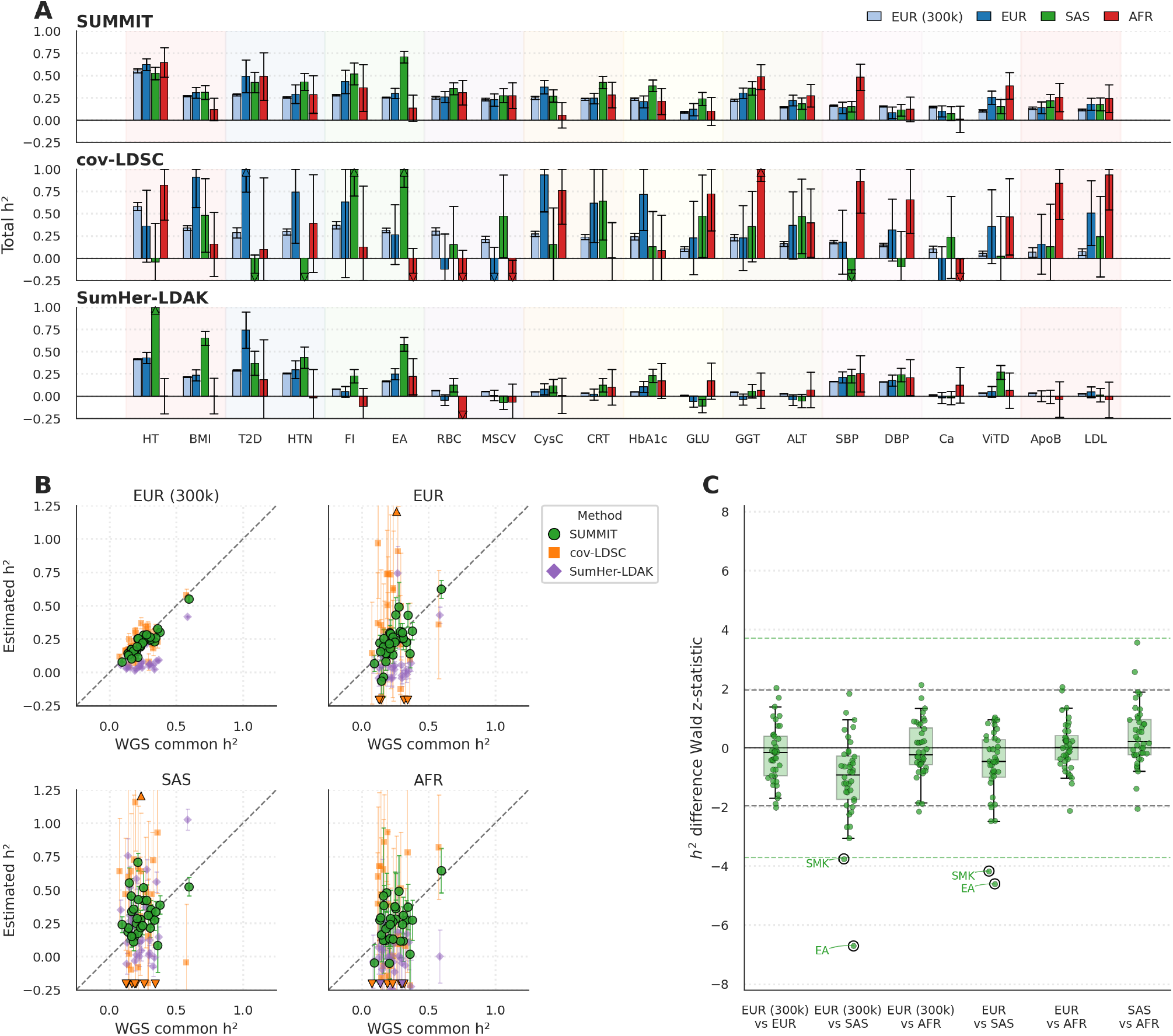
Genome-wide 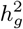 estimates of real traits in UKB. Genome-wide 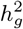 was estimated using a MAF-LD partitioned model with 24 MAF–LD bins (6 MAF, 4 LD). Out of 40 polygenic complex traits (30 of which overlap with Wainschtein et al. (2026)) used in our analysis, here we plot 2 traits from each of 10 broad categories. We only show 2 other representative methods for panels A and B. Results for all other methods, annotations, and 40 traits are available in Supplementary Figures S25–S27. **Panel A:** Genome-wide 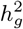 estimates from three methods, ordered by broad trait classes. **Panel B:** Plot of genome-wide 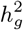 estimates vs UKB European WGS common variant estimates from Wainschtein et al. (2026). Bold triangles indicate points that are out of axes bounds. **Panel C:** SUMMIT test statistics for pairwise differences in common-SNP heritability across populations, computed for all 40 traits. For each trait and population pair, we tested for equality of heritability using a Wald *z* statistic. Dashed lines at *±*1.96 indicate nominal two-sided *p <* 0.05 and dashed lines at *±*3.71 indicate Bonferroni significance (240 tests for 40 traits *×* 6 population pairs; *p <* 0.05*/*240). SUMMIT identified EA and SMK as the only traits with Bonferroni-significant differences in 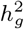, with significant differences between SAS and both European cohorts. The full list of Bonferroni-significant hits is available in Supplementary Table S5.

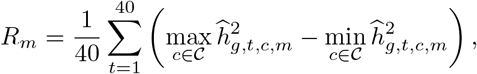

where 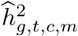 is the genome-wide estimate for trait *t*, cohort *c*, and method *m*, and *C* denotes the four cohorts. This statistic averages, over traits, the range of the four cohort-specific point estimates; smaller values indicate less variation in estimated 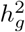 across cohorts. We observe that the cross-cohort range was substantially lower for SUMMIT (0.209) compared to cov-LDSC (0.965) and SumHer-LDAK (0.263). Among the 96 ancestry-trait pairs for which SUMMIT estimated 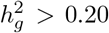, cov-LDSC returned 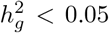 in 21 cases and SumHer-LDAK in 43 cases. We confirmed that the same qualitative pattern held when we used the coarser 8-bin MAF–LD annotation model instead of 24 bins (Supplementary Figs. S25 and S26).

We next used the European WGS common-variant (MAF *≥* 1%) estimates from Wainschtein et al. (2026) as a WGS-based benchmark for genome-wide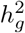. For each method and ancestry, we compared estimated genome-wide 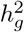 to the matched WGS estimate by calculating Pearson *r* and MSE relative to the WGS estimates (across 30 matched traits). In EUR (300k), SUMMIT more closely matched the WGS estimates (Pearson *r* = 0.93, MSE 0.2%) than cov-LDSC (*r* = 0.68, MSE = 0.7%) or SumHer-LDAK (*r* = 0.40, MSE = 2.7%) (Fig. 4B). We see qualitatively similar trends in the smaller non-European cohorts, with SUMMIT obtaining lower MSEs (3.1% and 2.2% in SAS and AFR compared with 56% and 19% for cov-LDSC and 6.7% and 9.1% for SumHer-LDAK).

We also tested whether the SUMMIT genome-wide 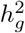 estimates for a trait differed across populations (Fig. 4C; full results in Supplementary Fig. S27 and Supplementary Table S5). The only traits with any significant difference in 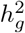 were educational attainment (EA) and smoking status (SMK). SAS had higher estimated 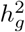 than EUR (300k) for EA 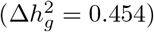 and for SMK 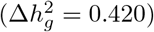, with similar trends when SAS estimates are compared to EUR. Our results mirror prior cross-ancestry results for EA (Momin et al. 2023). We caution both traits have been known to be confounded by population structure (Howe et al. 2022; Nivard et al. 2024; Pasman et al. 2022) so the observed differences in the 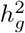 estimates are unlikely to be explained by differences in genetic architecture alone. Overall, our results are consistent with evidence that common-variant effects are broadly conserved across human populations (Yengo et al. 2022; Hou et al. 2023; Verma et al. 2024; Hu et al. 2025; Karczewski et al. 2025).

#### Heritability analysis reveals shared enrichment in constrained and regulatory annotations across populations

We next applied SUMMIT to genomic annotations (specifically the baseline-LD v2.2 annotations; see “Variant annotations” in Methods) to assess how its annotation-specific heritability estimates varied across ancestries. For each of EUR, SAS, and AFR cohorts, we computed the Pearson correlation *r* of the annotation-specific 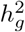 with respect to the corresponding estimates in EUR (300k) and meta-analyzed *r* across traits (correlations were Fisher *z*-transformed, combined across traits with inverse-variance weights, and transformed back to the correlation scale). SUMMIT gives the most concordant annotation-specific 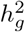 estimates with the EUR (300k) reference, with concordance steadily decreasing from EUR to SAS to AFR: the correlations for SUMMIT were *r* = 0.936, 0.931 and 0.571 for EUR, SAS and AFR, respectively, compared with 0.936, 0.822 and 0.435 for cov-LDSC (Fig. 5A; detailed method comparison and enrichment-profile summaries provided in Supplementary Fig. S28).

**Figure 5:**
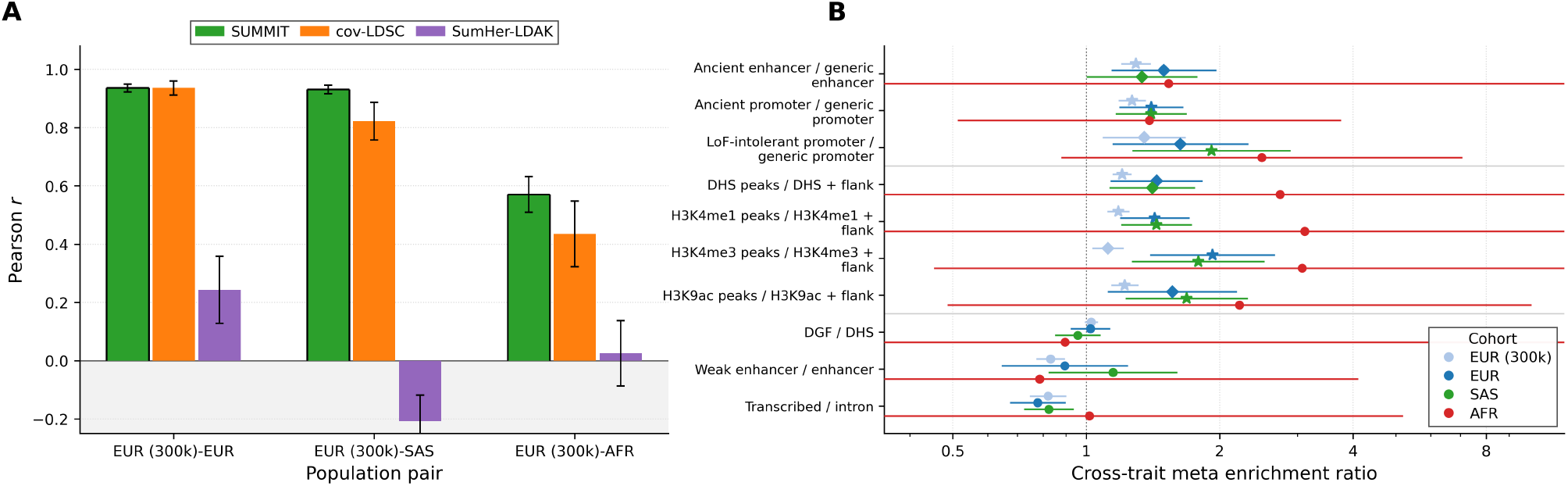
Cross-population concordance of partitioned heritability estimates and enrichment ratios. **Panel A:** Concordance of annotation-specific 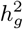 estimates for SUMMIT, cov-LDSC, and SumHer-LDAK. For each trait, we computed Pearson correlation *r* between the vectors of annotation-specific 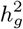 estimates in each of the cohorts (EUR, SAS, and AFR) and those in EUR (300k). Correlations were meta-analyzed across traits with error bars denoting standard error. **Panel B:** Ratio of enrichments between pairs of focal/background annotations across ancestries. Horizontal bars show 95% confidence intervals on a logarithmic ratio axis; intervals extending beyond the plotting range are truncated. Circles did not pass BH FDR *<* 0.05, diamonds passed BH FDR *<* 0.05 but not Bonferroni correction, and stars passed Bonferroni correction (*p <* 0.05*/*40 = 0.00125). The correction included 40 tests (10 annotation pairs *×* four cohorts). The first three pairs follow Hujoel et al. (2019); the next four compare core DHS or histone-mark peaks with their broader flanked annotations (Finucane et al. 2015; Gazal et al. 2017); the final three serve as negative controls.

Annotation-level heritability coefficients tend to be noisy in the smaller cohorts. We instead focused on whether there is an enrichment of per-SNP heritability in a focal annotation compared to a matched background annotation (Fig. 5B). For this analysis, we compared ancient enhancer with enhancer, ancient promoter with promoter, and LoF-intolerant promoter with promoter, following the comparisons of Hujoel et al. (2019) that showed increased enrichments in annotations that older sequence age and were more conserved. We also compared DHS, H3K4me1, H3K4me3, and H3K9ac peaks with their corresponding broader flanked annotations, motivated by prior evidence for heritability enrichment in core regulatory elements (Finucane et al. 2015; Gazal et al. 2017). As negative controls, we additionally tested DGF versus DHS, weak enhancer versus enhancer, and transcribed chromatin versus intron, for which we did not expect enrichment ratios greater than one a priori. This is equivalent to testing whether the per-SNP heritability is higher in the focal annotation (the numerator) compared to the broader annotation (the denominator).

For each pair of annotations compared above, we obtained estimates of the enrichment ratio and SEs by meta-analyzing across all traits. To do this, we estimated the log enrichment ratio and its (block-jackknife) standard error for each pair of annotations in a trait followed by meta-analyses of the log ratios. To estimate the overall standard error while accounting for correlations among traits, we recomputed the meta-analyzed estimate in each jackknife block. We tested each enrichment ratio using a one-sided Wald test of log(ratio) *>* 0 (with Benjamini–Hochberg FDR and Bonferroni correction applied across 40 tests: 10 annotation pairs *×* 4 cohorts).

We observed significant enrichment for ancient enhancer versus enhancer, ancient and LoF-intolerant promoter versus promoter, and DHS, H3K4me1, H3K4me3, and H3K9ac peaks versus their flanked annotations in EUR (300k), EUR, and SAS (BH FDR *<* 0.05); five, three, and five ratios, respectively, remained significant after Bonferroni correction (*p <* 0.05*/*40 = 0.00125). None of the comparisons were significant in AFR and none of the negative controls was significantly greater than one in any cohort. Finally, we also confirmed that the annotation-specific 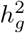 estimates in EUR (300k) from SUMMIT were concordant with prior estimates (Finucane et al. 2015; Gazal et al. 2017; Hujoel et al. 2019) and with estimates from imputed SNPs (MAF *>* 0.001) in the same individuals (Supplementary Figs. S29, S30, S31, and S32).

#### SUMMIT improves power for genetic correlation analysis across ancestries

We next estimated genome-wide genetic correlations using the MAF–LD model. This is motivated by our simulations that show variation in *r*_*g*_ across MAF–LD strata can bias *r*_*g*_ estimates based on a single-component model. Additionally, we found evidence of trait pairs that showed variation in *r*_*g*_ due to MAF–LD though such gradients were not pervasive (Supplementary Fig. S18).

SUMMIT produced substantially more complete genome-wide *r*_*g*_ maps than cov-LDSC across the traits (Fig. 6; all 40 traits and alternative annotations in Supplementary Figs S33–S36). More importantly, SUMMIT identified 621 EUR (300k), 23 SAS, and 3 AFR hits, compared with 195, 0, and 2 hits for cov-LDSC (BH FDR *<* 0.05; Fig. 7A–C), with most cov-LDSC hits also found to be significant in SUMMIT. All 23 SAS and three AFR SUMMIT hits were also significant in EUR (300k). The SAS estimates were strongly concordant with the corresponding EUR (300k) estimates (*r* = 0.95; Fig. 7E); the three AFR pairs are shown without a correlation coefficient (Fig. 7F). Direct tests comparing non-EUR estimates with EUR (300k) identified three SAS pairs with evidence of different *r*_*g*_ estimates (EA–FI, CRT–EA, and CysC–EA; BH FDR *<* 0.05) and no differences in AFR. Because all three SAS differences involved EA, a trait with inflated SAS heritability that we attribute to non-genetic effects rather than a real ancestry-specific signal, we interpret these differences cautiously. The same qualitative pattern held at the Bonferroni threshold (Supplementary Fig. S37). SUMMIT estimates were also stable across annotation choices (Supplementary Figs. S38, S39, S40, S41, S42).

**Figure 6:**
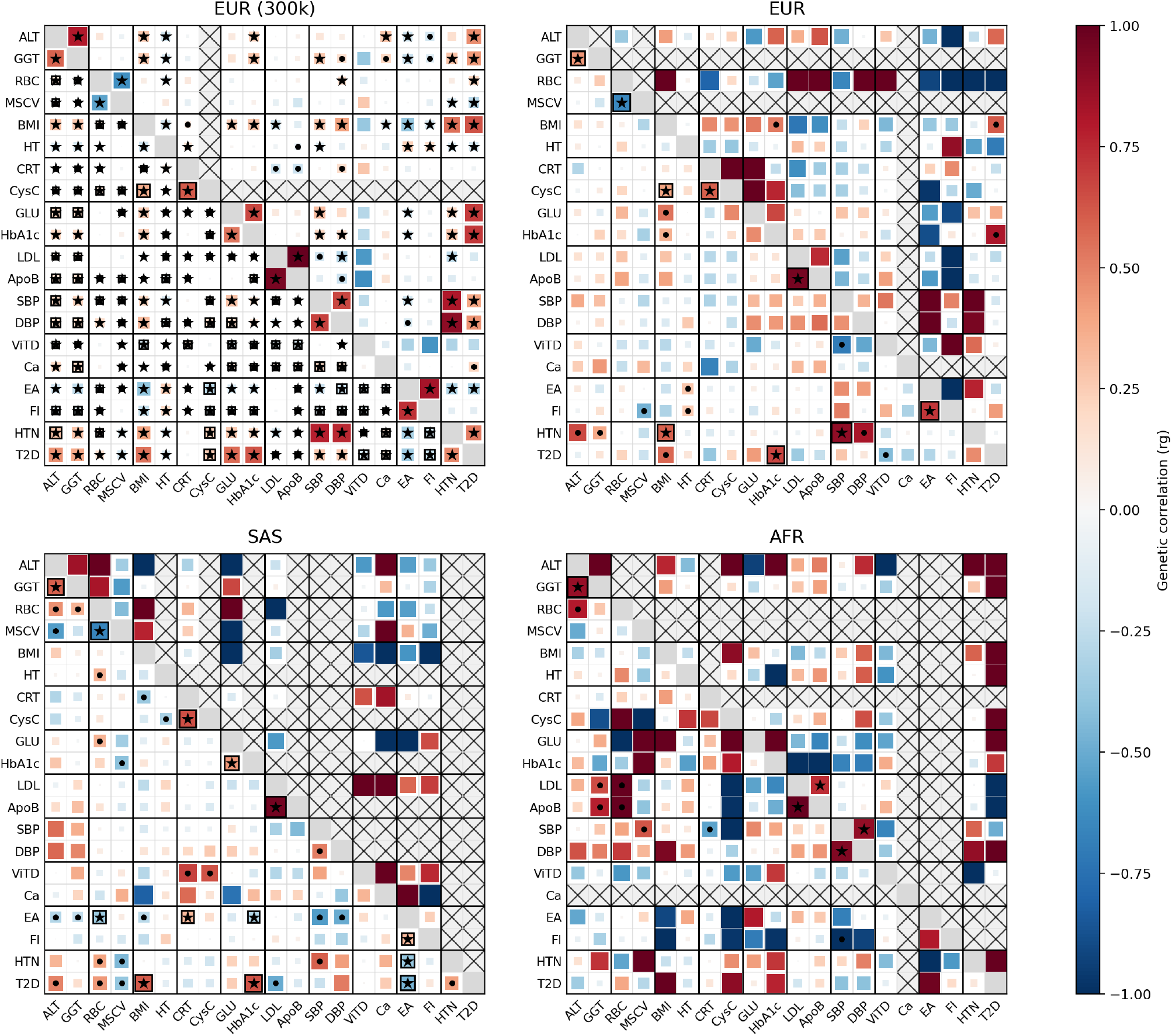
Heatmap of genome-wide *r*_*g*_ estimates. In each subplot, **lower triangle:** SUMMIT estimates of genome-wide *r*_*g*_ for each pair of traits. **upper triangle:** cov-LDSC estimates of genome-wide *r*_*g*_. Stars indicate significant pairs (BH FDR *<* 0.05), dots indicate nominally significant pairs, and black boxes indicate FDR-significant pairs for one method that are not significant in the other. Grey boxes with crosses indicate that a valid *r*_*g*_ estimate was not available because of negative heritability estimates. Results using all 40 traits are also available in Supplementary Figures S33–S36.

**Figure 7:**
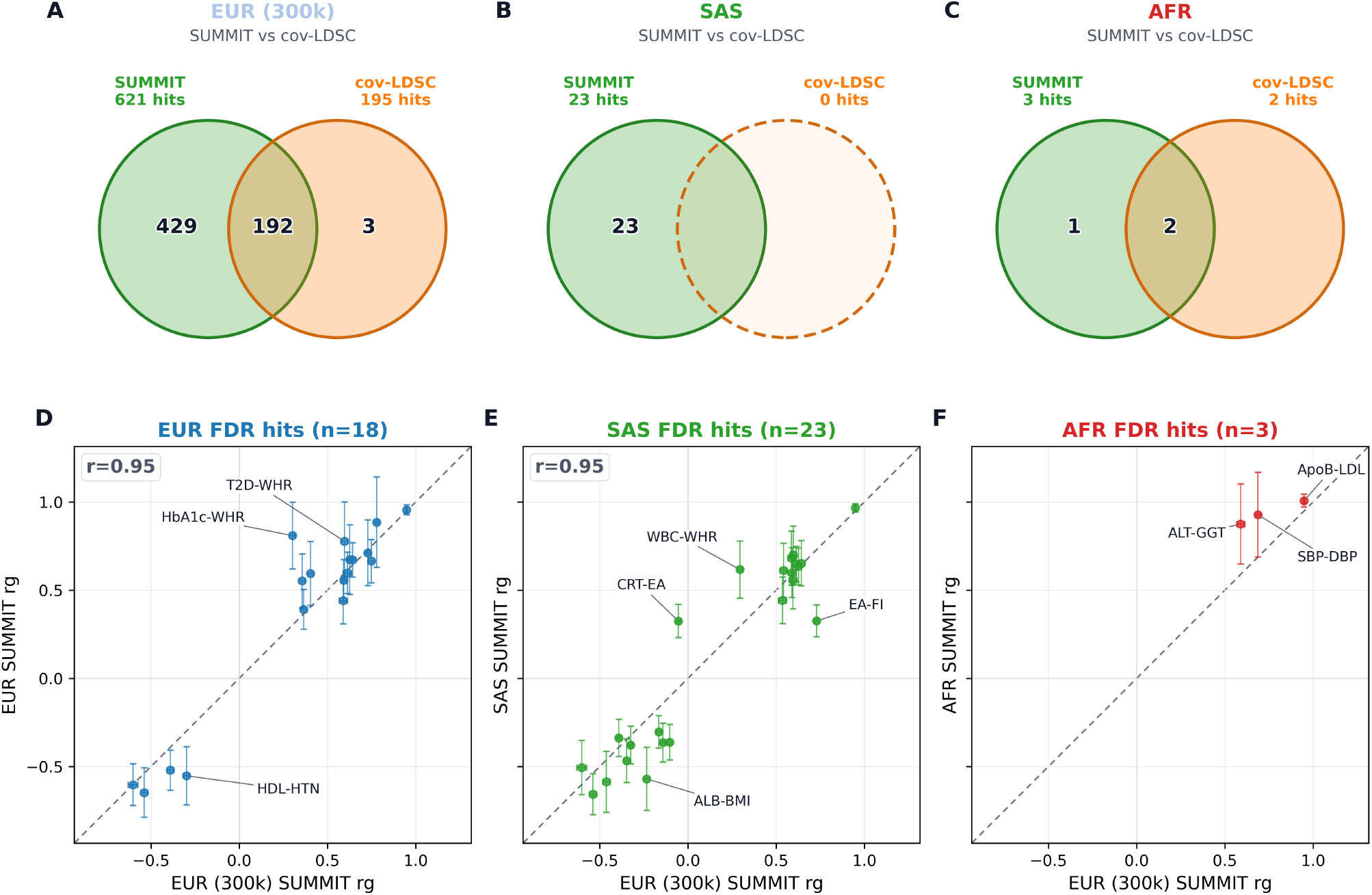
Cross-population concordance and method overlap of genome-wide *r*_*g*_ hits. **Panels A–C:** Overlap of SUMMIT and cov-LDSC FDR-significant hits within EUR (300k), SAS, and AFR. SUMMIT identifies more hits across populations with majority of the cov-LDSC hits also recovered by SUMMIT. **Panels D–F:** SUMMIT *r*_*g*_ estimates for FDR-significant hits in EUR, SAS, and AFR plotted against the corresponding EUR (300k) estimates. Error bars show one standard error, the dashed line marks equality. Pearson correlations are shown in Panels D and E (no correlation coefficient is reported in Panel F given only three AFR pairs were available).

#### SUMMIT provides robust heritability enrichment estimates with an external reference

While our analyses have primarily focused on the setting where in-sample genome-wide LD scores are available, we next explored the possibility of applying SUMMIT in a setting where the LD reference is obtained from a cohort that is distinct from that of the GWAS summary statistics. We harmonized external summary statistics from FinnGen, GLGC, MVP, Genes & Health, and T2DGGI to ancestry-matched UKB genotype references (Supplementary Table S6). We identified external summary statistics for 31 traits that could be matched to the 40 traits analyzed in UKB (14 in EUR, 7 in SAS, and 10 in AFR) and analyzed them using ancestry-matched UKB LD scores.

Prior work has shown that reference LD can bias heritability estimates while enrichments are more robust (Finucane et al. 2015). Consistent with these observations, estimates of 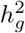 from SUMMIT showed substantial differences when comparing the estimates obtained from UKB to those in the external GWAS (as did cov-LDSC) reflecting the combined effects of reference mismatch, phenotype heterogeneity, and differences in GWAS scale on 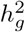 estimates (Supplementary Fig. S43). We therefore focused on heritability enrichment in 13 annotations representing selection, conservation, regulatory sequence age, core regulatory annotations and fine-mapped molecular QTLs motivated by prior work (Gazal et al. 2017; Hujoel et al. 2019; Hormozdiari et al. 2018). These annotations showed broadly concordant enrichment patterns between UKB and the external GWAS (Fig. 8A). In each ancestry, we attempted to test if a given annotation was enriched for 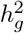 by testing if the median of the enrichment values for that annotation across external GWAS is significantly greater than 1 (one-sided test with SE estimated via SNP-level jackknife). We corrected jointly over 78 annotation-by-ancestry tests (13 annotations *×* three ancestries *×* two data sources: UKB and external GWAS). Among the external GWAS, 12 annotation-by-ancestry tests passed BH FDR *<* 0.05 (eight in EUR and four in SAS), and four passed Bonferroni correction (*p <* 0.05*/*78): ancient promoter age and BLUEPRINT methylation QTL in EUR, and CpG content and coding regions in SAS. No annotation was significantly enriched in AFR although all annotations had median enrichments greater than 1 in AFR. The corresponding UKB estimates showed the same directional pattern but with significant signals concentrated in the EUR (300k) cohort due to the smaller sample sizes of SAS (*N* = 7,912) and AFR (*N* = 3,416) in UKB compared to the external GWAS (SAS: *N ≈* 15,000–40,000 and AFR: *N ≈* 23,000–129,000).

**Figure 8:**
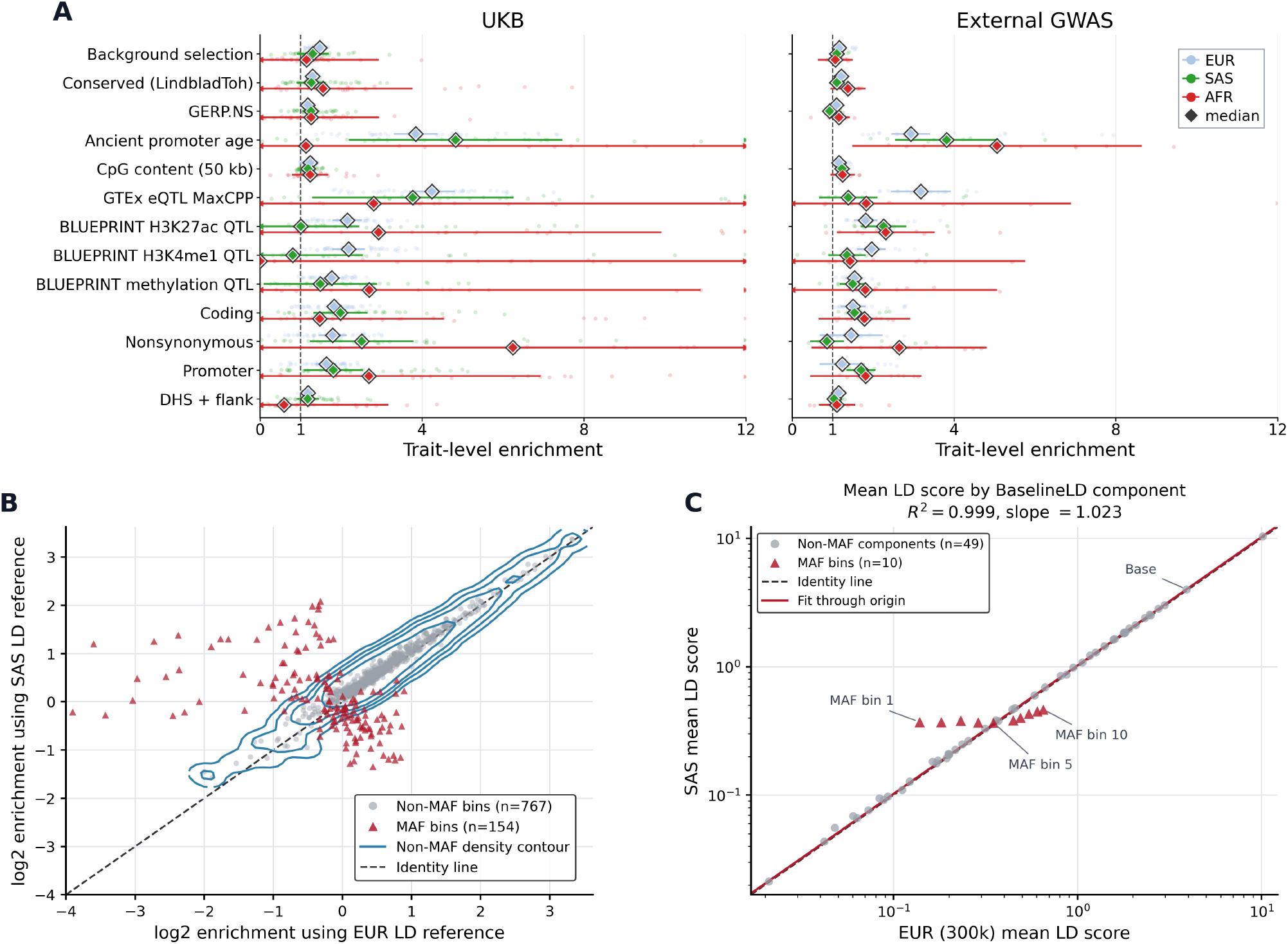
Functional enrichment in external GWAS and sensitivity to LD-reference mismatch. We curated external GWAS summary statistics that broadly matched phenotypes and cohorts in our UKB analysis and analyzed them using ancestry-matched UKB LD references. We estimated heritability enrichment and then evaluated sensitivity to a deliberately mismatched LD reference. **Panel A:** SUMMIT heritability enrichment estimates for selected genomic annotations in UKB and external GWAS. Faint points show trait-level enrichment estimates, colored by ancestry; diamonds show the median across estimates and horizontal segments show the jackknife SE. The enrichment axis is truncated at 12, with boundary arrows marking estimates extending outside the plotted window. **Panels B**,**C:** To evaluate sensitivity of enrichment estimates to the LD reference, SUMMIT was run on 16 external EUR GWAS using annotations from the the Baseline-LD v2.2 annotation model, analyzing each trait with either the EUR (300k) or deliberately mismatched SAS LD/annotation reference while holding the summary statistics fixed. **Panel B:** Log2 enrichment estimates using the SAS LD reference versus the EUR (300k) LD reference across trait-annotation estimates. Gray points denote annotations except MAF (*n* = 767), red triangles denote MAF-bin annotations (*n* = 154), blue contours show the smoothed density of non-MAF annotations, and the dashed line marks equality. The lower plotting bound is truncated at *−*4 for display. **Panel C:** Mean LD scores for each annotation computed on the SAS and EUR (300k) references. Gray circles denote the annotations except MAF and red triangles denote MAF bins; the dashed line marks equality and the solid red line is the fit through the origin across the non-MAF annotations. We additionally compare SNP-level LD-scores in Supplementary Fig. S44B.

To isolate the effect of the LD reference, we analyzed 16 external EUR GWAS using either the precomputed EUR (300k) or SAS Baseline-LD resource. The summary statistics and SNP set were held fixed, while each reference was used with its supplied genome-wide LD scores and annotations. Enrichment estimates for annotations other than MAF remained consistent across references (*r* = 0.989; Fig. 8B), whereas total and annotation-specific 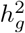 estimates shifted downward with the SAS reference (Supplementary Fig. S44A). Although SNP-level LD scores differed between references (Supplementary Fig. S44B), the mean LD scores for the annotations (except the MAF annotations) were consistent (Fig. 8C). Thus, enrichment estimates can remain informative when reference mismatch affects the fitted annotations by approximately the same factor while annotations that are reference-specific e.g., MAF, tend to be more sensitive.

## Discussion

We developed SUMMIT, a novel framework for estimating heritability and genetic correlation across diverse populations. Through an unbiased stochastic estimator for genome-wide LD scores, SUMMIT addresses a key bottleneck in summary-level analysis – the presence of long-range LD – of structured and admixed populations while remaining computationally efficient. In simulations with real UK Biobank genotypes, SUMMIT improved precision even in European cohorts. In the non-European cohorts (SAS and AFR), SUMMIT also reduced bias from long-range LD missed by prior approaches.

Applied to 40 real complex traits, SUMMIT produced common-SNP heritability estimates that are broadly comparable across cohorts for many anthropometric and biomarker traits and are robust to the specific annotation models used, with estimates close to recent WGS-based estimates in the UKB European ancestry individuals (Wainschtein et al. 2026). We find that large cross-ancestry differences in common-SNP heritability are uncommon, consistent with prior cross-ancestry analyses (Yengo et al. 2022; Hou et al. 2023; Verma et al. 2024; Hu et al. 2025; Karczewski et al. 2025): only two traits (educational attainment and smoking status) were significantly different between SAS and EUR. SUMMIT also improved genetic-correlation estimation: recovering most cov-LDSC discoveries while also identifying a substantial number of additional trait pairs in all three ancestries. The trait-pairs identified by SUMMIT to have significant genetic correlation estimates in non-European ancestries were also significant in EUR with strongly correlated estimates, lending support to the global pleiotropy being shared across ancestries. SUMMIT produced more stable *h*^2^ enrichment estimates across cohorts showing enrichment in constrained and evolutionarily old regulatory annotations and in core regulatory peaks. These patterns agree with prior partitioned-heritability work linking complex-trait heritability to functional annotations, evolutionary constraint and regulatory sequence age (Finucane et al. 2015; Gazal et al. 2017; Hujoel et al. 2019; Zeng et al. 2018).

Our application of SUMMIT to external-GWAS reveal the limits of reference-based summary-statistics analysis. We demonstrate that SUMMIT behaves similarly to LDSC in this regard, as reference mismatch caused biased absolute heritability estimates, while enrichment retained substantial signal. These results support cautious use of SUMMIT for heritability enrichment under moderate reference mismatch, but accurate absolute 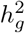 estimates still require closely matched, and preferably in-sample, LD scores.

Several limitations remain. First, our models of admixed populations assume shared causal effects across the ancestral segments. While supported for common variants in African-European individuals (Hou et al. 2023) including the AFR cohort from UKB analyzed here, this assumption may not hold for specific traits, admixed populations, genomic regions or classes of variants. This limitation motivates building local-ancestry-aware LD scores (more below). Second, SUM-CORE requires the cross-trait moment to be on the same HE scale. This is straightforward when sample overlap and phenotype covariance are known, or when GWAS are non-overlapping, but it may not be possible for public GWAS releases that do not report sufficient information. Third, annotation-rich models remain statistically and biologically difficult in smaller cohorts, especially for genetic correlations and conditional annotation coefficients. Fourth, our real-data analyses focus on UK Biobank, with much smaller non-European cohorts limiting the power of our analyses.

A natural next step is to build reusable genome-wide LD-score resources for diverse biobanks and consortia, so that summary-statistics analyses can be performed without repeatedly accessing individual-level phenotypes. This is especially relevant as resources such as All of Us, MVP and GBMI expand the scale and diversity of human genetic studies (All of Us Research Program Genomics Investigators 2024; Verma et al. 2024; Zhou et al. 2022). Given the computational and statistical efficiency of SUMMIT, a phenome-wide scan in large biobanks with limited compute is also possible. Methodologically, extending SUMMIT to local-ancestry-aware LD scores would help separate ancestry-specific tagging from shared genetic effects in admixed cohorts, where recent work suggests that common-variant causal effects are often highly correlated across local ancestry backgrounds once LD and tagging are modeled carefully (Hou et al. 2023; Hu et al. 2025). WGS-scale inference is another important direction, especially for rare variants and rare non-coding annotations where frequency effects, LD sparsity and annotation imbalance are more severe. In its current form, SUMMIT provides a practical summary-statistics tool for estimating where common-SNP heritability and cross-trait sharing are concentrated, while delineating the recoverable signal under LD-reference mismatch.

## Methods

### Notation and setup

Let *M* denote the number of SNPs and let *j ∈ {*1, …, *M*} index SNPs. Let 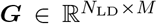 denote the in-sample or reference genotype matrix used to estimate LD quantities, and let 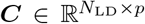 denote the corresponding covariates. After covariate residualization and re-standardization, we write the genotype matrix as ***X***. Let ***A*** *∈* ℝ^*M ×K*^ denote the annotation matrix, with *k, ℓ ∈ {*1, …, *K}* indexing annotations. For a disjoint binary partition, *S*_*k*_ = *{j* : *A*_*jk*_ = 1*}* and *M*_*k*_ = |*S*_*k*_|. For continuous or overlapping annotations, we use *A*_*jk*_ as the annotation weight and define 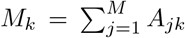 the version of the method applied to continuous-valued annotations is given in Supplementary Note S1.7.

For trait *a, N*_*a*_ denotes the GWAS sample size used and *n*_*aj*_ denotes the SNP-specific sample count. In single-trait analyses we write *N ≡ N*_1_. In bivariate analyses, *a ∈ {*1, 2*}, N*_1_ and *N*_2_ denote the trait-specific sample sizes and *Ñ* denotes the number of overlapping individuals. The genome-wide LD quantities are computed once per population and reused across traits and sample sizes. For all our benchmarks, we use in-sample genotype to estimate the LD scores unless specified otherwise.

### Estimation of genome-wide LD scores

We estimate genome-wide partitioned LD scores once per population and reuse them across all downstream analyses. We construct a standardized genotype matrix 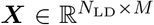 by centering and scaling each SNP. If covariates are present, we residualize genotypes using the same covariates used for the corresponding GWAS/LD analysis, and then re-standardize each SNP column to have mean 0 and sample variance 1 in the LD cohort. This matched covariate adjustment follows the principle of covariate-adjusted LD score regression (Luo et al. 2021) and is particularly important in structured or admixed cohorts, where long-range LD can otherwise induce substantial confounding (Price et al. 2008).

Define the sample LD correlation between SNPs *j* and *j*′ by

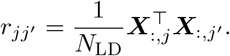

For a binary annotation *k*, the partitioned LD score is

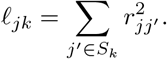

For weighted or overlapping annotations, we use the corresponding weighted form,

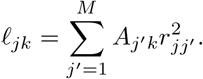

Computing all *ℓ*_*jk*_ directly would require forming genome-wide LD matrices which scales as *O*(*N*_*LD*_*M* ^2^). Instead, we estimate these LD scores with randomized projections. We draw independent spherical random vectors ***z***^(*v*)^ *∈* ℝ^*M*^, *v* = 1, …, *B*, uniformly from the sphere of radius 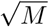 so that E[***z***^(*v*)^***z***^(*v*)*⊤*^] = ***I***_*M*_.

For each bin *k*, form

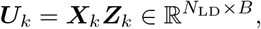

where ***X***_*k*_ denotes the columns of ***X*** indexed by *S*_*k*_ and ***Z***_*k*_ denotes the rows of the random projection matrix indexed by *S*_*k*_. Then

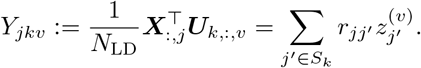

By isotropy of the spherical vectors,

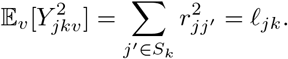

This gives an estimator of the genome-wide LD score:

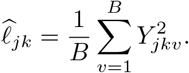

Finally, we apply a finite-sample correction to obtain the genome-wide LD-score estimates used in all down-stream analyses,

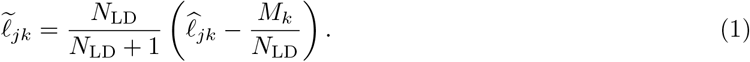

The derivation of the finite-sample correction, the treatment of diagonal SNP pairs, and the chunked implementation are described in Supplementary Note S1.1. In practice, we stream SNP blocks and never materialize dense *M × M* LD matrices.

### HE-scale summary-statistics

All primary analyses use summary statistics (which we term the “HE-scale summary statistics”) that are obtained by transforming standard GWAS summary statistics to recover the method-of-moments (MoM) estimates. For trait *a*, we transform the marginal OLS coefficient and standard error for SNP *j* to an HE-scale statistic 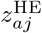 as described in Supplementary Note S1.2. We then define

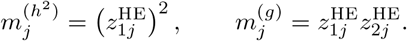

### Heritability estimation from summary statistics that recovers individual-level estimates

Our approach estimates SNP heritability either genome-wide or in specific annotations (when provided) using only GWAS summary statistics and precomputed genome-wide LD scores. Given GWAS summary statistics and genome-wide LD scores, we reconstruct the normal equations that are used by individual-level methods such as RHE-mc (Pazokitoroudi et al. 2020) thereby obtaining an estimator with the accuracy of an individual-level method while only requiring access to summary statistics. This is a direct extension of Jeong et al. (2024) to support annotation-rich models and the use of LD scores.

Let ***y*** *∈* ℝ^*N*^ be a standardized phenotype vector and ***X*** *∈* ℝ^*N ×M*^ the corresponding standardized genotype matrix. We consider the *K*-component variance component model

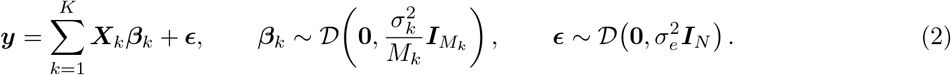

Define 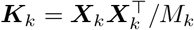. Then

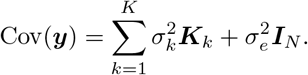

Matching the model covariance to ***yy***^*⊤*^ gives the MoM normal equations

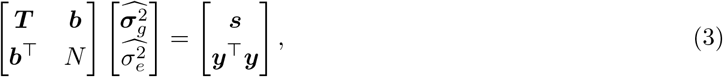

where, for *k, ℓ ∈ {*1, …, *K}*,

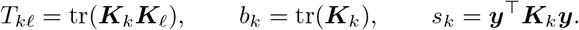

With standardized genotypes, *b*_*k*_ = *N*.

The trace terms are reconstructed from the genome-wide LD scores (Supplementary Note S1.3) as

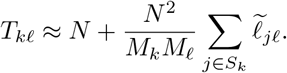

The right-hand side can be computed from GWAS summary statistics with an appropriate finite-sample adjustment (which we term the “HE-scale moments”):

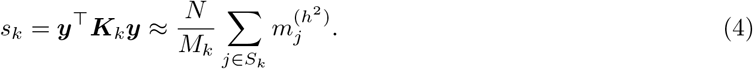

Equations (3)–(4) define the MoM normal equations for the genetic and residual variance components using GWAS summary statistics and precomputed genome-wide LD scores, and extends the estimator proposed by GWASH (Schwartzman et al. 2019) to genome-wide, annotation-specific LD scores. From summary statistics alone, the variance components are identified only up to the overall phenotype scale, since ***y***^*⊤*^***y*** is needed to determine the corresponding residual variance 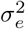. For heritability estimation, however, this absolute scale is not required. SUM-RHE therefore works on the normalized scale ***y***^*⊤*^***y*** = *N*, under which the genetic variance components are the partitioned heritabilities, and solves the same normal equations to estimate 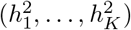.

### Genetic covariance and correlation estimation from summary statistics

SUM-CORE extends our previous individual-level genetic correlation method, SCORE (Wu et al. 2022), to summary statistics by reconstructing the moment equations that previously needed individual-level data with genome-wide LD scores and GWAS summary statistics. Let *a ∈ {*1, 2*}* index traits. Let 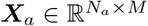 contain the standardized SNPs for trait *a*, and let ***y***_*a*_ be the corresponding standardized phenotype vector. We parameterize the cross-trait genetic covariance through the second moment

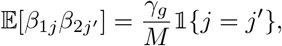

where *γ*_*g*_ is the genome-wide genetic covariance. Similar to the univariate case, we only work with moments and no distributional assumption on the SNP effects is required.

Let *Ñ* denote the number of overlapping individuals and define the cross-GRM

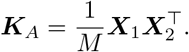

SCORE solves the two-equation MoM system

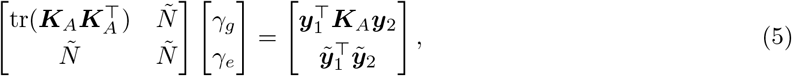

where 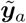 denotes the phenotype vector restricted to the overlapping individuals and *γ*_*e*_ captures the overlap-induced non-genetic contribution.

Let

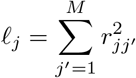

denote the genome-wide LD score for SNP *j*. After eliminating *γ*_*e*_, the summary-statistic reconstruction gives the aggregate moment

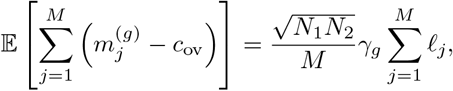

where *c*_ov_ denotes the population overlapping phenotype covariance on the HE scale. Its supplied sample estimate is

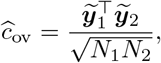

computed from the projected and normalized phenotype values for individuals present in both GWAS. In the default supplied-overlap mode, *ĉ*_ov_ is provided as an input. For non-overlapping GWAS, *c*_ov_ = *ĉ*_ov_ = 0.

Given *ĉ*_ov_, the reconstructed SCORE estimator is

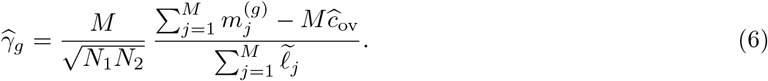

We then report genetic correlation by normalizing with SUM-RHE heritability estimates,

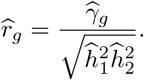

SUM-CORE also supports an ancillary summary-only mode in which *ĉ*_ov_ is estimated internally by robust bivariate LD-score regression applied to the HE-scale moments. This mode is useful when no external estimate of the overlapping phenotype covariance is available, but it is not the primary estimator used in this paper; details and benchmarks are provided in Supplementary Note S1.5 and Supplementary Fig. S19.

### Partitioned genetic covariance and correlation

For annotation-specific analyses, SUM-CORE uses the same reconstruction with annotation-specific LD-score aggregates, yielding a *K*-component linear system for component-specific genetic covariances *γ*_*k*_; the aggregate moment system, weighted/overlapping annotation extension, and ridge-stabilized solution are given in Supplementary Note S1.6. We report partitioned genetic correlations as

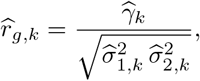

using the corresponding SUM-RHE component heritability estimates.

### Standard errors

We compute standard errors by a leave-one-block-out SNP jackknife unless otherwise stated. For each jackknife replicate, we remove one approximately equal-sized SNP block and recompute the estimator from the remaining blocks. For SUM-CORE in supplied-overlap mode, the primary jackknife conditions on the supplied *ĉ*_ov_; in summary-only mode, *ĉ*_ov_ is re-estimated within each replicate. Standard errors for derived quantities, including enrichment statistics and partitioned genetic correlations, are obtained by recomputing the derived quantity in each jackknife replicate and applying the standard jackknife variance formula (see Supplementary Note S1.8).

### UK Biobank cohorts and traits

We curate three UK Biobank cohorts spanning homogeneous and admixed settings. **AFR:** starting from the 4,327 African-European admixed individuals with *≥* 5% AFR and *≥* 5% EUR ancestry used in Hou et al. (2023), we further restrict to 3,416 unrelated samples by removing 3rd-degree relatives (PLINK 2.0 KING; Manichaikul et al. (2010)). After QC on HM3 SNPs (MAF *≥* 0.01, HWE *≥* 10^*−*8^, excluding the MHC), we retain 1,002,034 variants (array + well-imputed). **SAS:** we select South Asian individuals by clustering samples with pan-UKB genotype PCs and self-identified ethnicity (Indian, Pakistani, Bangladeshi), removing outliers. Applying similar variant QC (common, non-MHC autosomal) and restricting to unrelated samples yields 7,912 individuals and 448,403 array SNPs. **EUR / EUR (300k):** we use the 291,273 unrelated white British individuals on *M* = 454, 207 common array SNPs (MAF *>* 0.01, non-MHC), matching Jeong et al. (2024). For simulation benchmarks we use a *N* = 10, 060 subsample and denote it as EUR; the full cohort is denoted EUR (300k). For Baseline-LD annotation results we additionally perform a sensitivity analysis using the EUR (300k) MAF *>* 10^*−*3^ imputed panel with 7,774,235 SNPs, of which 37.9% had MAF *<* 0.01.

For real-trait analyses, we analyze 40 UK Biobank traits spanning continuous anthropometric traits, blood biomarkers, behavioral traits and binary disease traits (Supplementary Data). Because the available common-SNP set differs across cohorts after population-aware QC, we analyze the largest high-quality non-MHC common-SNP set available in each cohort rather than forcing all populations onto a smaller intersection. We report SNP-set overlap and MAF distributions in Supplementary Fig. S1. Cross-cohort comparisons below should therefore be interpreted as stability of cohort-specific common-SNP heritability and genetic-correlation estimates under a shared QC and annotation pipeline.

### Variant annotations

For MAF–LD partitioning, we calculated in-sample MAF and 2-Mb local LD scores once within each cohort. We partitioned SNPs into either two MAF bins (MAF *<* 0.1, MAF *≥* 0.1) or six MAF bins: [0, 0.025), [0.025, 0.05), [0.05, 0.075), [0.075, 0.10), [0.10, 0.125), and [0.125, 0.50] in EUR and SAS; and [0, 0.05), [0.05, 0.10), [0.10, 0.15), [0.15, 0.20), [0.20, 0.25), and [0.25, 0.50] in AFR. We crossed these bins with four cohort-specific local-LD-score quartiles to form 8- or 24-component MAF–LD models. The resulting cohort-specific annotation assignments were held fixed across methods and were used in both simulation benchmarks and real-data analyses. SUMMIT used genome-wide annotation-specific LD scores for fitting, cov-LDSC used 20-Mb annotation-specific LD scores, and SumHer used 20-Mb taggings.

For functional annotation, we started from the Baseline-LD model v2.2 annotation panel from the LDSC/Price-lab resource (Finucane et al. 2015; Gazal et al. 2017). The released local Baseline-LD v2.2 panel contains 97 annotation columns. We aligned these annotations to the UKBB analysis SNP sets and, when both a focal annotation X and its `X.flanking.500` companion were present, replaced them with a single `X_w_flanking` indicator. Together with retained unpaired annotations and population-specific MAF decile bins, the collapsed analysis panel contained 59 annotation columns. For example, `DHS_Trynka` and `DHS_Trynka.flanking.500` were combined as `DHS_Trynka_w_flanking`. Unless otherwise specified, enrichment results are reported for these collapsed annotation groups rather than the raw released columns.

### Benchmark methods

We ran all benchmark methods on the same post-QC, non-MHC SNP set used for each cohort-specific analysis. Simulated phenotypes were generated from standardized genotypes under single-component, MAF–LD-partitioned, coding-enrichment, overlapping-annotation, or signed genetic-covariance designs. GWAS summary statistics were computed with PLINK 2.0 (v2.0.0-a.6LM, 14 October 2024). Conventional summary-statistic methods used the unadjusted simulated GWAS unless noted below; cov-LDSC and SUMMIT used PLINK2 ‘–glm hide-covar’ summary statistics adjusted for in-sample genotype PCs and matched covariate-adjusted LD scores. We used 10 PCs for AFR, EUR and EUR (300k), and 20 PCs for SAS in the heritability simulations. In real-data analyses, additional covariates of age, sex, age *×* sex, age^2^, age^2^*×* sex were used. Unlike in simulated phenotypes, all methods used the covariate-adjusted GWAS summary statistics for estimation.

LDSC was run with LDSC v1.0.1 using ‘–h2’, ‘–ref-ld’, ‘–w-ld’ and ‘–not-M-5-50’. Simulation runs also used ‘–chisq-max 99999’. LDSC estimates used in the main simulation comparison used 20-Mb LD-score windows, with other window sizes retained for sensitivity analyses. cov-LDSC used the same regression code with covariate-adjusted LD scores computed after projecting out the same covariates used in the GWAS.

SumHer was run with LDAK version 6. For each simulated trait, we first generated LDAK summary files with ‘–linear’ and then estimated heritability with ‘–sum-hers’ using in-sample tagging and matrix files. We report two SumHer settings: a GCTA-style tagging model (‘–power −1’, ‘–ignore-weights YES’), denoted SumHer-GCTA, and the LDAK tagging model (‘–power −0.25’), denoted SumHer-LDAK. The main simulation comparison used 20-Mb taggings, with smaller tagging windows used for sensitivity checks. For real-trait partitioned analyses, LDAK GWAS summary files were generated with the same phenotype and covariate files used for the PLINK2 analyses before running SumHer with the in-sample tagging files.

HDL was run with HDL v1.4.1 in R. We used cohort-specific HDL LD references built from the matched UK Biobank genotype data, set ‘Nref’ to the corresponding reference-panel sample size, and used ‘jack-knife.df=TRUE’.

RHE-mc was run through GENIE v1.2.1 using the additive genetic model (‘-m G’), 100 random vectors and 100 jackknife blocks.

## Software availability

https://github.com/sriramlab/SUMMIT

## Competing interest statement

The authors declare no competing interests.

## Acknowledgments

This research was conducted using the UK Biobank Resource under application 33127. We thank the participants of UK Biobank for making this work possible.

## Notes

https://github.com/sriramlab/SUMMIT

### Competing Interest Statement

The authors have declared no competing interest.

### Funding

- National Institutes of Health: GM125055, HG006399
- National Science Foundation: CAREER-1943497

## References

Abul-Husn NS and Kenny EE. 2019. Personalized medicine and the power of electronic health records. Cell 177: 58–69.

All of Us Research Program Genomics Investigators. 2024. Genomic data in the All of Us Research Program. Nature 627: 340–346.

Bulik-Sullivan B, Finucane HK, Anttila V, Gusev A, Day FR, Loh PR, ReproGen Consortium, Psychiatric Genomics Consortium, Genetic Consortium for Anorexia Nervosa of the Wellcome Trust Case Control Consortium 3, Duncan L, et al. 2015a. An atlas of genetic correlations across human diseases and traits. Nature Genetics 47: 1236–1241.

Bulik-Sullivan BK, Loh PR, Finucane HK, Ripke S, Yang J, Schizophrenia Working Group of the Psychiatric Genomics Consortium, Patterson N, Daly MJ, Price AL, and Neale BM. 2015b. LD score regression distinguishes confounding from polygenicity in genome-wide association studies. Nature Genetics 47: 291–295.

Bycroft C, Freeman C, Petkova D, Band G, Elliott LT, Sharp K, Motyer A, Vukcevic D, Delaneau O, O’Connell J, et al. 2018. The UK Biobank resource with deep phenotyping and genomic data. Nature 562: 203–209.

Finer S, Martin HC, Khan A, Hunt KA, MacLaughlin B, Ahmed Z, Ashcroft R, Durham C, MacArthur DG, McCarthy MI, et al. 2020. Cohort profile: East London Genes & Health (ELGH), a community-based population genomics and health study in British Bangladeshi and British Pakistani people. International Journal of Epidemiology 49: 20–21i.

Finucane HK, Bulik-Sullivan B, Gusev A, Trynka G, Reshef Y, Loh PR, Anttila V, Xu H, Zang C, Farh K, et al. 2015. Partitioning heritability by functional annotation using genome-wide association summary statistics. Nature Genetics 47: 1228–1235.

Gazal S, Finucane HK, Furlotte NA, Loh PR, Palamara PF, Liu X, Schoech A, Bulik-Sullivan B, Neale BM, Gusev A, et al. 2017. Linkage disequilibrium-dependent architecture of human complex traits shows action of negative selection. Nature Genetics 49: 1421–1427.

Gazal S, Marquez-Luna C, Finucane HK, and Price AL. 2019. Reconciling S-LDSC and LDAK functional enrichment estimates. Nature Genetics 51: 1202–1204.

Gaziano JM, Concato J, Brophy M, Fiore L, Pyarajan S, Breeling J, Whitbourne S, Deen J, Shannon C, Humphries D, et al. 2016. Million Veteran Program: A mega-biobank to study genetic influences on health and disease. Journal of Clinical Epidemiology 70: 214–223.

Graham SE, Clarke SL, Wu KHH, et al. 2021. The power of genetic diversity in genome-wide association studies of lipids. Nature 600: 675–679.

Hormozdiari F, Gazal S, van de Geijn B, Finucane HK, Ju CJT, Loh PR, Schoech A, Reshef Y, Liu X, O’Connor L, et al. 2018. Leveraging molecular quantitative trait loci to understand the genetic architecture of diseases and complex traits. Nature Genetics 50: 1041–1047.

Hou K, Burch KS, Majumdar A, Shi H, Mancuso N, Wu Y, Sankararaman S, and Pasaniuc B. 2019. Accurate estimation of SNP-heritability from biobank-scale data irrespective of genetic architecture. Nature Genetics 51: 1244–1251.

Hou K, Ding Y, Xu Z, Wu Y, Bhattacharya A, Mester R, Belbin GM, Buyske S, Conti DV, Darst BF, et al. 2023. Causal effects on complex traits are similar for common variants across segments of different continental ancestries within admixed individuals. Nature Genetics 55: 549–558.

Howe LJ, Nivard MG, Morris TT, Hansen AF, Rasheed H, Cho Y, Chittoor G, Ahlskog R, Lind PA, Palviainen T, et al. 2022. Within-sibship genome-wide association analyses decrease bias in estimates of direct genetic effects. Nature Genetics 54: 581–592.

Hu S, Ferreira LAF, Shi S, Hellenthal G, Marchini J, Lawson DJ, and Myers SR. 2025. Fine-scale population structure and widespread conservation of genetic effect sizes between human groups across traits. Nature Genetics 57: 379–389.

Hujoel MLA, Gazal S, Hormozdiari F, van de Geijn B, and Price AL. 2019. Disease heritability enrichment of regulatory elements is concentrated in elements with ancient sequence age and conserved function across species. The American Journal of Human Genetics 104: 611–624.

Jacobs BM, Stow D, Hodgson S, et al. 2024. Genetic architecture of routinely acquired blood tests in a British South Asian cohort. Nature Communications 15: 8929.

Jeong M, Pazokitoroudi A, Liu Z, and Sankararaman S. 2024. Scalable summary-statistics-based heritability estimation method with individual genotype level accuracy. Genome Research 34: 1286–1293.

Karczewski KJ, Gupta R, Kanai M, et al. 2025. Pan-UK Biobank genome-wide association analyses enhance discovery and resolution of ancestry-enriched effects. Nature Genetics 57: 2408–2417.

Kurki MI, Karjalainen J, Palta P, et al. 2023. FinnGen provides genetic insights from a well-phenotyped isolated population. Nature 613: 508–518.

Lu Q, Li B, Ou D, Erlendsdottir M, Powles RL, Jiang T, Hu Y, Chang D, Jin C, Dai W, et al. 2017. A powerful approach to estimating annotation-stratified genetic covariance via GWAS summary statistics. The American Journal of Human Genetics 101: 939–964.

Luo Y, Li X, Wang X, Gazal S, Mercader JM, 23 and Me Research Team, SIGMA Type 2 Diabetes Consortium, Neale BM, Florez JC, Auton A, et al. 2021. Estimating heritability and its enrichment in tissue-specific gene sets in admixed populations. Human Molecular Genetics 30: 1521–1534.

Manichaikul A, Mychaleckyj JC, Rich SS, Daly K, Sale M, and Chen WM. 2010. Robust relationship inference in genome-wide association studies. Bioinformatics 26: 2867–2873.

Momin MM, Shin J, Lee S, Truong B, Benyamin B, and Lee SH. 2023. A method for an unbiased estimate of cross-ancestry genetic correlation using individual-level data. Nature Communications 14: 722.

Muller ME. 1959. A note on a method for generating points uniformly on N-dimensional spheres. Communications of the ACM 2: 19–20.

Ning Z, Pawitan Y, and Shen X. 2020. High-definition likelihood inference of genetic correlations across human complex traits. Nature Genetics 52: 859–864.

Nivard MG, Belsky DW, Harden KP, Baier T, Andreassen OA, Ystrøm E, van Bergen E, and Lyngstad TH. 2024. More than nature and nurture, indirect genetic effects on children’s academic achievement are consequences of dynastic social processes. Nature Human Behaviour 8: 771–778.

Pasman JA, Demange PA, Guloksuz S, Willemsen AHM, Abdellaoui A, ten Have M, Hottenga JJ, Boomsma DI, de Geus E, Bartels M, et al. 2022. Genetic risk for smoking: disentangling interplay between genes and socioeconomic status. Behavior Genetics 52: 92–107.

Pazokitoroudi A, Wu Y, Burch KS, Hou K, Zhou A, Pasaniuc B, and Sankararaman S. 2020. Efficient variance components analysis across millions of genomes. Nature Communications 11: 4020.

Pfaff CL, Parra EJ, Bonilla C, Hiester K, McKeigue PM, Kamboh MI, Hutchinson RG, Ferrell RE, Boerwinkle E, and Shriver MD. 2001. Population structure in admixed populations: Effect of admixture dynamics on the pattern of linkage disequilibrium. The American Journal of Human Genetics 68: 198–207.

Price AL, Weale ME, Patterson N, Myers SR, Need AC, Shianna KV, Ge D, Rotter JI, Torres E, Taylor KD, et al. 2008. Long-range LD can confound genome scans in admixed populations. The American Journal of Human Genetics 83: 132–135.

Schoech AP, Jordan DM, Loh PR, Gazal S, O’Connor LJ, Balick DJ, Palamara PF, Finucane HK, Sunyaev SR, and Price AL. 2019. Quantification of frequency-dependent genetic architectures in 25 UK Biobank traits reveals action of negative selection. Nature Communications 10: 790.

Schwartzman A, Schork AJ, Zablocki R, and Thompson WK. 2019. A simple, consistent estimator of SNP heritability from genome-wide association studies. The Annals of Applied Statistics 13: 2509–2538.

Shi H, Kichaev G, and Pasaniuc B. 2016. Contrasting the genetic architecture of 30 complex traits from summary association data. The American Journal of Human Genetics 99: 139–153.

Shi H, Mancuso N, Spendlove S, and Pasaniuc B. 2017. Local genetic correlation gives insights into the shared genetic architecture of complex traits. The American Journal of Human Genetics 101: 737–751.

Speed D and Balding DJ. 2019. SumHer better estimates the SNP heritability of complex traits from summary statistics. Nature Genetics 51: 277–284.

Speed D, Hemani G, Johnson MR, and Balding DJ. 2012. Improved heritability estimation from genome-wide SNPs. The American Journal of Human Genetics 91: 1011–1021.

Suzuki K, Hatzikotoulas K, Southam L, et al. 2024. Genetic drivers of heterogeneity in type 2 diabetes pathophysiology. Nature 627: 347–357.

Verma A, Huffman JE, Rodriguez A, et al. 2024. Diversity and scale: Genetic architecture of 2068 traits in the VA Million Veteran Program. Science 385: eadj1182.

Wainschtein P et al. 2026. Estimation and mapping of the missing heritability of human phenotypes. Nature 649: 1219–1227.

Wu Y, Burch KS, Ganna A, Pajukanta P, Pasaniuc B, and Sankararaman S. 2022. Fast estimation of genetic correlation for biobank-scale data. The American Journal of Human Genetics 109: 24–32.

Wu Y and Sankararaman S. 2018. A scalable estimator of SNP heritability for biobank-scale data. Bioinformatics 34: i187–i194.

Yang J, Lee SH, Goddard ME, and Visscher PM. 2011. GCTA: A tool for genome-wide complex trait analysis. The American Journal of Human Genetics 88: 76–82.

Yengo L, Vedantam S, Marouli E, et al. 2022. A saturated map of common genetic variants associated with human height. Nature 610: 704–712.

Zeng J, de Vlaming R, Wu Y, Robinson MR, Lloyd-Jones LR, Yengo L, Yap CX, Xue A, Sidorenko J, McRae AF, et al. 2018. Signatures of negative selection in the genetic architecture of human complex traits. Nature Genetics 50: 746–753.

Zhang Y, Lu Q, Ye Y, Huang K, Liu W, Wu Y, Zhong X, Li B, Yu Z, Travers BG, et al. 2021. SUPERGNOVA: Local genetic correlation analysis reveals heterogeneous etiologic sharing of complex traits. Genome Biology 22: 262.

Zhou W et al. 2022. Global Biobank meta-analysis initiative: Powering genetic discovery across human disease. Cell Genomics 2: 100192.

