## Supplementary Information for "Genome-wide LD scores enable robust heritability and genetic correlation estimation from summary statistics across genetic ancestries"

**S1 Supplementary Note** **2**

### S1 Supplementary Note

#### S1.1 Genome-wide LD scores

##### S1.1.1 Random-projection estimator

Let $\boldsymbol{X} \in \mathbb{R}^{N_{\mathrm{LD}} \times M}$ be the standardized (optionally covariate-residualized) genotype matrix in the LD reference cohort. Define the sample LD correlation

$$r_{jj'} = \frac{1}{N_{\mathrm{LD}}} \boldsymbol{X}_{:,j}^\top \boldsymbol{X}_{:,j'}. \tag{7}$$

For a disjoint bin $k$ with SNP set $S_k$ and size $M_k = |S_k|$, we define the in-sample LD score as

$$\ell_{jk} = \sum_{j' \in S_k} r_{jj'}^2. \tag{8}$$

Which is the convention used throughout this paper. In particular, when $j \in S_k$, the sum includes the SNP self-correlation $r_{jj}^2 = 1$.

Draw independent probe vectors $\boldsymbol{z}^{(v)} \in \mathbb{R}^M$ uniformly from the sphere of radius $\sqrt{M}$, $v = 1, \ldots, B$. Equivalently, sample $\boldsymbol{g}^{(v)} \sim \mathcal{N}(\boldsymbol{0}, \boldsymbol{I}_M)$ and set

$$\boldsymbol{z}^{(v)} = \sqrt{M} \frac{\boldsymbol{g}^{(v)}}{\|\boldsymbol{g}^{(v)}\|_2}, \tag{9}$$

which yields $\boldsymbol{z}^{(v)}$ uniform on the sphere (Muller 1959) and satisfies

$$\mathbb{E}\left[\boldsymbol{z}^{(v)} \boldsymbol{z}^{(v)\top}\right] = \boldsymbol{I}_M. \tag{10}$$

For each bin $k$, define

$$Y_{jkv} := \sum_{j' \in S_k} r_{jj'} z_{j'}^{(v)}. \tag{11}$$

Conditioning on $\boldsymbol{X}$ and using (10),

$$\begin{aligned} \mathbb{E}\left[Y_{jkv}^2 \mid \boldsymbol{X}\right] &= \mathbb{E}\left[\left(\sum_{j' \in S_k} r_{jj'} z_{j'}^{(v)}\right)^2 \middle| \boldsymbol{X}\right] \\ &= \sum_{j' \in S_k} r_{jj'}^2 = \ell_{jk}. \end{aligned} \tag{12}$$

Therefore,

$$\widehat{\ell}_{jk} := \frac{1}{B} \sum_{v=1}^{B} Y_{jkv}^2 \qquad \text{satisfies} \qquad \mathbb{E}\left[\widehat{\ell}_{jk} \mid \boldsymbol{X}\right] = \ell_{jk}. \tag{13}$$

For nonnegative weighted or overlapping annotations, we use

$$\ell_{jk} = \sum_{j'=1}^{M} A_{j'k} r_{jj'}^2, \qquad M_k = \sum_{j'=1}^{M} A_{j'k}. \tag{14}$$

The randomized estimator is obtained by weighting each probe contribution by $\sqrt{A_{j'k}}$:

$$Y_{jkv} = \sum_{j'=1}^{M} r_{jj'} \sqrt{A_{j'k}}\, z_{j'}^{(v)}. \tag{15}$$

By isotropy,

$$\mathbb{E}_v[Y_{jkv}^2] = \sum_{j'=1}^{M} A_{j'k} r_{jj'}^2 = \ell_{jk}. \tag{16}$$

The contribution from the SNP self-correlation is $A_{jk}$. For a disjoint binary annotation, $A_{j'k} = \mathbb{1}\{j' \in S_k\}$, and this contribution reduces to $\mathbb{1}\{j \in S_k\}$.

##### S1.1.2 Finite-sample correction

Let $(X_j, X_{j'})$ denote a generic standardized genotype pair from the LD reference population, with

$$\rho_{jj'} = \mathrm{Cov}(X_j, X_{j'}).$$

Then the *population* LD score is:

$$\ell_{jk}^\star = \sum_{j'=1}^{M} A_{j'k} \rho_{jj'}^2. \tag{17}$$

For a disjoint binary annotation, $A_{j'k} = \mathbb{1}\{j' \in S_k\}$ and $M_k = \sum_{j'=1}^{M} A_{j'k} = |S_k|$. For $j \neq j'$, write

$$r_{jj'} = \frac{1}{N_{\mathrm{LD}}} \sum_{i=1}^{N_{\mathrm{LD}}} u_i, \qquad u_i = X_{ij} X_{ij'}.$$

Assume $(X_j, X_{j'})$ is jointly Gaussian with mean 0, variance 1, and covariance $\rho_{jj'}$. Then $\mathbb{E}[u_i] = \rho_{jj'}$ and $\mathbb{E}[u_i^2] = 1 + 2\rho_{jj'}^2$ by Isserlis' theorem. Hence

$$\mathbb{E}[r_{jj'}^2] = \frac{1 + \rho_{jj'}^2}{N_{\mathrm{LD}}} + \rho_{jj'}^2 = \frac{N_{\mathrm{LD}} + 1}{N_{\mathrm{LD}}} \rho_{jj'}^2 + \frac{1}{N_{\mathrm{LD}}}. \tag{18}$$

Taking expectation over the random probes and the LD reference cohort, and separating the SNP self-correlation, for which $r_{jj}^2 = \rho_{jj}^2 = 1$, gives

$$\mathbb{E}[\widehat{\ell}_{jk}] = \frac{N_{\mathrm{LD}} + 1}{N_{\mathrm{LD}}} \ell_{jk}^\star + \frac{M_k}{N_{\mathrm{LD}}} - \frac{2A_{jk}}{N_{\mathrm{LD}}}. \tag{19}$$

Under this Gaussian calculation, an exactly unbiased correction would be

$$\frac{N_{\mathrm{LD}}}{N_{\mathrm{LD}} + 1} \left( \widehat{\ell}_{jk} - \frac{M_k}{N_{\mathrm{LD}}} \right) + \frac{2A_{jk}}{N_{\mathrm{LD}} + 1}.$$

Dropping the $O(N_{\mathrm{LD}}^{-1})$ term leads to the genome-wide LD score estimate:

$$\widetilde{\ell}_{jk} = \frac{N_{\mathrm{LD}}}{N_{\mathrm{LD}} + 1} \left( \widehat{\ell}_{jk} - \frac{M_k}{N_{\mathrm{LD}}} \right), \tag{20}$$

For a disjoint binary annotation, the omitted term is nonzero only when $j \in S_k$ and is at most $2/(N_{\mathrm{LD}} + 1)$.

###### S1.1.3 Chunking and computational complexity

We compute $\boldsymbol{X}_k \boldsymbol{Z}_k$ and $\boldsymbol{X}^\top (\boldsymbol{X}_k \boldsymbol{Z}_k)$ in SNP chunks, streaming blocks of columns of $\boldsymbol{X}$ and accumulating intermediate products without forming dense $M \times M$ matrices. Runtime scales as

$$\mathcal{O}\left( N_{\mathrm{LD}} \, B \sum_k M_k \right),$$

up to constants reflecting annotation masking and I/O, and peak memory is controlled by the SNP chunk size and $B$. Numerical settings used in the experiments as well as corresponding runtime and memory usage are summarized in Supplementary Tables S3,S4.

#### S1.2 Exact HE-scale moments from summary statistics

All primary analyses use HE-scale summary-statistics moments. These are the finite-sample moments corresponding to covariate-projected and re-standardized SNPs and phenotypes in the HE/SCORE moment equations. We work with the covariate-adjusted marginal model

$$\boldsymbol{y}_a = \boldsymbol{C}_a \boldsymbol{\alpha}_a + \boldsymbol{x}_{aj} \beta_{aj} + \boldsymbol{\varepsilon}_a, \qquad a \in \{1, 2\},$$

where $\boldsymbol{C}_a$ includes the intercept column and all GWAS covariates. Define

$$\boldsymbol{P}_a = \boldsymbol{I} - \boldsymbol{C}_a (\boldsymbol{C}_a^\top \boldsymbol{C}_a)^{-1} \boldsymbol{C}_a^\top.$$

Let $q_a = \operatorname{rank}(\boldsymbol{C}_a)$, including the intercept column, and let $N_a$ denote the effective GWAS sample size used to normalize the HE-scale moment for trait $a$. For SNP $j$, define

$$d_{aj} = \boldsymbol{x}_{aj}^\top \boldsymbol{P}_a \boldsymbol{x}_{aj}, \qquad R_a = \boldsymbol{y}_a^\top \boldsymbol{P}_a \boldsymbol{y}_a.$$

Normalize projected SNPs and phenotypes by

$$\tilde{\boldsymbol{x}}_{aj} = \sqrt{\frac{N_a}{d_{aj}}} \boldsymbol{P}_a \boldsymbol{x}_{aj}, \qquad \tilde{\boldsymbol{y}}_a = \sqrt{\frac{N_a}{R_a}} \boldsymbol{P}_a \boldsymbol{y}_a,$$

so that $\tilde{\boldsymbol{x}}_{aj}^\top \tilde{\boldsymbol{x}}_{aj} = N_a$ and $\tilde{\boldsymbol{y}}_a^\top \tilde{\boldsymbol{y}}_a = N_a$.

##### S1.2.1 Derivation from OLS summary statistics

By the Frisch–Waugh–Lovell theorem,

$$\widehat{\beta}_{aj} = \frac{\boldsymbol{x}_{aj}^\top \boldsymbol{P}_a \boldsymbol{y}_a}{\boldsymbol{x}_{aj}^\top \boldsymbol{P}_a \boldsymbol{x}_{aj}} = \frac{u_{aj}}{d_{aj}}, \qquad u_{aj} := \boldsymbol{x}_{aj}^\top \boldsymbol{P}_a \boldsymbol{y}_a.$$

Since the SNP coefficient is estimated after projecting out $q_a$ covariates, the residual degrees of freedom are $\nu_{aj} = n_{aj} - q_a - 1$, where $n_{aj}$ is the regression sample count for SNP $j$. The residual sum of squares for the marginal regression is $R_a - \widehat{\beta}_{aj}^2 d_{aj}$, giving

$$\widehat{\mathrm{SE}}_{aj}^2 = \frac{R_a - \widehat{\beta}_{aj}^2 d_{aj}}{\nu_{aj} d_{aj}}.$$

Solving for $d_{aj}$ gives

$$d_{aj} = \frac{R_a}{\widehat{\beta}_{aj}^2 + \nu_{aj} \widehat{\mathrm{SE}}_{aj}^2}.$$

Define the HE-scale statistic

$$z_{aj}^{\mathrm{HE}} := \frac{\tilde{\boldsymbol{x}}_{aj}^\top \tilde{\boldsymbol{y}}_a}{\sqrt{N_a}}.$$

Then

$$z_{aj}^{\mathrm{HE}} = \frac{1}{\sqrt{N_a}} \left( \sqrt{\frac{N_a}{d_{aj}}} \boldsymbol{P}_a \boldsymbol{x}_{aj} \right)^\top \left( \sqrt{\frac{N_a}{R_a}} \boldsymbol{P}_a \boldsymbol{y}_a \right) \tag{21}$$

$$= \sqrt{N_a}\, \widehat{\beta}_{aj} \sqrt{\frac{d_{aj}}{R_a}} = \frac{\sqrt{N_a} \widehat{\beta}_{aj}}{\sqrt{\widehat{\beta}_{aj}^2 + \nu_{aj} \widehat{\mathrm{SE}}_{aj}^2}}. \tag{22}$$

##### S1.2.2 Univariate SUM-RHE moment

Define

$$m_j^{(h^2)} := \left(z_j^{\mathrm{HE}}\right)^2.$$

For a component $k$,

$$\tilde{\boldsymbol{y}}^\top \tilde{\boldsymbol{K}}_k \tilde{\boldsymbol{y}} = \frac{1}{M_k} \sum_{j \in S_k} \left(\tilde{\boldsymbol{x}}_j^\top \tilde{\boldsymbol{y}}\right)^2 = \frac{N}{M_k} \sum_{j \in S_k} m_j^{(h^2)}. \tag{23}$$

Thus $m_j^{(h^2)}$ is the exact HE-scale replacement for $z_j^2$ in SUM-RHE. Under a multi-component model,

$$\mathbb{E}\left[m_j^{(h^2)}\right] = 1 + \frac{N}{M} \sum_{c=1}^{K} \tau_c \widetilde{\ell}_{jc},$$

with the precise coefficient parameterization depending on whether the model is written in terms of per-component variance components $\sigma_c^2$ or per-SNP effects $\tau_c$.

##### S1.2.3 Bivariate SUM-CORE moment

For two traits, define

$$m_j^{(g)} := z_{1j}^{\mathrm{HE}} z_{2j}^{\mathrm{HE}} = \frac{\sqrt{N_1 N_2}\, \widehat{\beta}_{1j} \widehat{\beta}_{2j}}{\sqrt{\left(\widehat{\beta}_{1j}^2 + \nu_{1j} \widehat{\mathrm{SE}}_{1j}^2\right)\left(\widehat{\beta}_{2j}^2 + \nu_{2j} \widehat{\mathrm{SE}}_{2j}^2\right)}}.$$

Then

$$\tilde{\boldsymbol{y}}_1^\top \tilde{\boldsymbol{K}}_{12} \tilde{\boldsymbol{y}}_2 = \frac{1}{M} \sum_{j=1}^{M} \left(\tilde{\boldsymbol{x}}_{1j}^\top \tilde{\boldsymbol{y}}_1\right)\left(\tilde{\boldsymbol{x}}_{2j}^\top \tilde{\boldsymbol{y}}_2\right) = \frac{\sqrt{N_1 N_2}}{M} \sum_{j=1}^{M} m_j^{(g)}. \tag{24}$$

Thus $m_j^{(g)}$ is the exact HE-scale replacement for the Wald product $z_{1j} z_{2j}$ in SUM-CORE.

##### S1.2.4 Relation to ordinary Wald $z$-scores

Let $z_j^{\mathrm{W}} = \widehat{\beta}_j / \widehat{\mathrm{SE}}_j$ denote the usual Wald statistic. Then

$$z_j^{\mathrm{HE}} = \sqrt{\frac{N}{\nu_j}} \frac{z_j^{\mathrm{W}}}{\sqrt{1 + (z_j^{\mathrm{W}})^2 / \nu_j}},$$

and

$$m_j^{(g)} = \sqrt{\frac{N_1 N_2}{\nu_{1j} \nu_{2j}}} \frac{z_{1j}^{\mathrm{W}} z_{2j}^{\mathrm{W}}}{\sqrt{\left(1 + (z_{1j}^{\mathrm{W}})^2 / \nu_{1j}\right)\left(1 + (z_{2j}^{\mathrm{W}})^2 / \nu_{2j}\right)}}.$$

Under the usual small-effect regime $(z_j^{\mathrm{W}})^2 = o(\nu_j)$ and $N_a / \nu_{aj} \to 1$, the HE-scale moments and Wald moments are asymptotically equivalent. In finite samples, however, the HE-scale moments are the exact quantities that correspond to the projected kernels used in SUM-RHE and SUM-CORE. Consequently, intercepts estimated from ordinary Wald products, including standard bivariate LDSC intercepts (Bulik-Sullivan et al. 2015a), are not automatically on the HE scale used by SUM-CORE.

#### S1.3 Partitioned heritability: trace and quadratic-form identities

Let $\boldsymbol{X} \in \mathbb{R}^{N\times M}$ be the standardized GWAS genotype matrix for a single trait and $\boldsymbol{X}_k$ the SNP columns in bin $S_k$. Define $\boldsymbol{K}_k = \boldsymbol{X}_k\boldsymbol{X}_k^\top/M_k$. Then

$$\begin{aligned} T_{k\ell} \equiv \operatorname{tr}(\boldsymbol{K}_k\boldsymbol{K}_\ell) &= \frac{1}{M_kM_\ell}\operatorname{tr}\big(\boldsymbol{X}_k\boldsymbol{X}_k^\top\boldsymbol{X}_\ell\boldsymbol{X}_\ell^\top\big) = \frac{1}{M_kM_\ell}\|\boldsymbol{X}_k^\top\boldsymbol{X}_\ell\|_F^2 \\ &= \frac{N^2}{M_kM_\ell}\sum_{j\in S_k}\sum_{j'\in S_\ell} r_{jj'}^2 \\ &= \frac{N^2}{M_kM_\ell}\sum_{j\in S_k}\ell_{j\ell} \end{aligned} \tag{25}$$

We then approximate $T_{k\ell}$ from the genome-wide LD score as:

$$T_{k\ell} \approx N + \frac{N^2}{M_kM_\ell}\sum_{j\in S_k}\widetilde{\ell}_{j\ell}, \tag{26}$$

The quadratic-form right-hand side is reconstructed from HE-scale moments:

$$\boldsymbol{y}^\top\boldsymbol{K}_k\boldsymbol{y} \approx \frac{N}{M_k}\sum_{j\in S_k} m_j^{(h^2)}. \tag{27}$$

#### S1.4 Genetic correlation: SCORE reconstruction with overlapping phenotype covariance

##### S1.4.1 Cross-trace identity with sample overlap

Let $\boldsymbol{X}_1 \in \mathbb{R}^{N_1 \times M}$ and $\boldsymbol{X}_2 \in \mathbb{R}^{N_2 \times M}$ contain the same standardized SNPs for two traits, and let $\tilde{N}$ be the number of overlapping individuals. Define $\boldsymbol{K}_A = \boldsymbol{X}_1 \boldsymbol{X}_2^\top / M$. Let $\rho_{jk} = \mathrm{Cov}(x_{nj}, x_{nk})$ and $\ell_j = \sum_{k=1}^{M} \rho_{jk}^2$.

Consider

$$T := \mathrm{tr}\big(\boldsymbol{X}_1 \boldsymbol{X}_2^\top \boldsymbol{X}_2 \boldsymbol{X}_1^\top\big) = \sum_{i_1=1}^{N_1} \sum_{i_2=1}^{N_2} \left( \sum_{j=1}^{M} x_{1j}^{(i_1)} x_{2j}^{(i_2)} \right)^2 . \tag{28}$$

For non-overlapping pairs, independence across individuals gives

$$\mathbb{E}\Big[(x_1^{(i_1)} \cdot x_2^{(i_2)})^2\Big] = \sum_{j=1}^{M} \sum_{k=1}^{M} \rho_{jk}^2 = \sum_{j=1}^{M} \ell_j .$$

For overlapping pairs, $x_1^{(i)} = x_2^{(i)} = x^{(i)}$ and a Gaussian moment calculation gives

$$\mathbb{E}\Big[(x^{(i)} \cdot x^{(i)})^2\Big] = M^2 + 2 \sum_{j=1}^{M} \ell_j .$$

Combining the $(N_1 N_2 - \tilde{N})$ non-overlapping pairs with the $\tilde{N}$ overlapping pairs yields

$$\mathbb{E}[T] = (N_1 N_2 + \tilde{N}) \sum_{j=1}^{M} \ell_j + \tilde{N} M^2 .$$

Therefore

$$\mathbb{E}\big[\mathrm{tr}(\boldsymbol{K}_A \boldsymbol{K}_A^\top)\big] = \frac{N_1 N_2 + \tilde{N}}{M^2} \sum_{j=1}^{M} \ell_j + \tilde{N} \tag{29}$$

##### S1.4.2 Eliminating the overlap term

Starting from the SCORE MoM system (5), the second equation gives

$$\tilde{N}(\gamma_g + \gamma_e) = \tilde{\boldsymbol{y}}_1^\top \tilde{\boldsymbol{y}}_2 ,$$

and hence

$$\tilde{N} \gamma_e = \tilde{\boldsymbol{y}}_1^\top \tilde{\boldsymbol{y}}_2 - \tilde{N} \gamma_g .$$

Substituting this into the first equation yields

$$\begin{aligned} \boldsymbol{y}_1^\top \boldsymbol{K}_A \boldsymbol{y}_2 &= \mathrm{tr}(\boldsymbol{K}_A \boldsymbol{K}_A^\top) \gamma_g + \tilde{N} \gamma_e \\ &= \Big\{ \mathrm{tr}(\boldsymbol{K}_A \boldsymbol{K}_A^\top) - \tilde{N} \Big\} \gamma_g + \tilde{\boldsymbol{y}}_1^\top \tilde{\boldsymbol{y}}_2 . \end{aligned} \tag{30}$$

Using Eq. (29) and the HE-scale quadratic-form identity (24), let $c_{\mathrm{ov}}$ denote the population overlapping phenotype covariance on the HE scale. Its supplied sample estimate is

$$\widehat{c}_{\mathrm{ov}} := \frac{\tilde{\boldsymbol{y}}_1^\top \tilde{\boldsymbol{y}}_2}{\sqrt{N_1 N_2}} , \tag{31}$$

where $\tilde{\boldsymbol{y}}_a$ denotes the entries of the projected and normalized phenotype $\tilde{\boldsymbol{y}}_a$ restricted to the individuals overlapping between the two GWAS samples. Then the aggregated SUM-CORE moment is

$$\mathbb{E}\left[\sum_{j=1}^{M} m_j^{(g)}\right] = \frac{N_1 N_2 + \tilde{N}}{M\sqrt{N_1 N_2}}\gamma_g \sum_{j=1}^{M} \ell_j + M c_{\text{ov}} \approx \frac{\sqrt{N_1 N_2}}{M}\gamma_g \sum_{j=1}^{M} \ell_j + M c_{\text{ov}} \tag{32}$$

Equivalently,

$$\mathbb{E}[m_j^{(g)}] = b\ell_j + c_{\text{ov}}, \qquad b = \frac{\sqrt{N_1 N_2}}{M}\gamma_g. \tag{33}$$

Summing (33) over SNPs gives

$$\widehat{\gamma}_g = \frac{M}{\sqrt{N_1 N_2}} \frac{\sum_{j=1}^{M} m_j^{(g)} - M\widehat{c}_{\text{ov}}}{\sum_{j=1}^{M} \widetilde{\ell}_j}. \tag{34}$$

##### S1.4.3 Default supplied-overlap mode

The primary SUM-CORE analyses use supplied-overlap mode. In this mode, $\widehat{c}_{\text{ov}}$ is an input to the method and is treated as fixed in the main jackknife unless otherwise stated. The supplied quantity must correspond to the same covariate adjustment, phenotype normalization, SNP filtering, and HE-scale moment definition as $m_j^{(g)}$. When the two GWAS are known a priori to be non-overlapping, such as summary statistics from two independent biobanks, we set $c_{\text{ov}} = \widehat{c}_{\text{ov}} = 0$.

#### S1.5 SUM-CORE summary-only overlap-covariance estimation

When no external HE-scale estimate of the overlapping phenotype covariance is available, SUM-CORE supports an ancillary summary-only mode. Let

$$x_j = \widetilde{\ell}_j, \qquad z_j^{\mathrm{HE}} = m_j^{(g)}.$$

We fit the HE-scale regression moment

$$z_j^{\mathrm{HE}} = bx_j + c_{\mathrm{ov}} + \varepsilon_j$$

by weighted least squares with weights $w_j = 1/x_j$:

$$\min_{b, c_{\mathrm{ov}}} \sum_{j=1}^{M} \frac{1}{x_j}(z_j^{\mathrm{HE}} - bx_j - c_{\mathrm{ov}})^2.$$

Define

$$L_1 = \sum_{j=1}^{M} x_j, \qquad T_1 = \sum_{j=1}^{M} z_j^{\mathrm{HE}}, \qquad S_0 = \sum_{j=1}^{M} \frac{1}{x_j}, \qquad T_0 = \sum_{j=1}^{M} \frac{z_j^{\mathrm{HE}}}{x_j}.$$

The weighted normal equations reduce to

$$L_1 b + M c_{\mathrm{ov}} = T_1, \qquad Mb + S_0 c_{\mathrm{ov}} = T_0,$$

so

$$\widehat{c}_{\mathrm{ov,WLS}} = \frac{L_1 T_0 - M T_1}{L_1 S_0 - M^2}, \qquad \hat{b}_{\mathrm{WLS}} = \frac{T_1 - M\widehat{c}_{\mathrm{ov,WLS}}}{L_1}. \tag{35}$$

The first weighted normal equation is exactly the aggregated SCORE/SUM-CORE moment (32). Thus the WLS regression is used only to estimate the overlapping phenotype covariance; once $\widehat{c}_{\mathrm{ov}}$ is obtained, $\widehat{\gamma}_g$ is computed by Eq. (34).

In practice we use a leave-one-block-out version of (35), optionally excluding SNPs with extremely large marginal association statistics when fitting $\widehat{c}_{\mathrm{ov}}$. For small sample sizes, SNP-level genome-wide LD scores can be noisy for this fit. In that case, we use deterministic covariate-adjusted LD scores from a fixed window (20 Mb in the current benchmarks) to estimate $\widehat{c}_{\mathrm{ov}}$, while the final genetic covariance estimate is still obtained from the reconstructed SCORE equations using stochastic genome-wide LD scores. Because this mode is less stable than supplied-overlap mode, it is used only in supplementary benchmarks (Supplementary Fig. S19).

**Relation to bivariate LDSC.** Equation (33) has the same form as bivariate LDSC (Bulik-Sullivan et al. 2015a), but the direction of derivation is different. Here the regression arises by eliminating the overlap term from the SCORE moment equations and then reconstructing the resulting scalar identity from HE-scale summary statistics. Standard bivariate LDSC is written in terms of ordinary Wald $z$-score products, estimates an intercept on that scale, and uses variance-based regression weights with an LD redundancy correction to improve efficiency. Since SUM-CORE uses $m_j^{(g)}$ on the HE scale, the LDSC intercept is not directly interchangeable with $c_{\mathrm{ov}}$ in Eq. (33).

#### S1.6 Partitioned genetic correlation

With the same SNP partition $\{S_k\}_{k=1}^K$, we parameterize annotation-specific cross-trait genetic covariance by

$$\mathrm{Cov}(\boldsymbol{\beta}_{1,k}, \boldsymbol{\beta}_{2,k}) = \frac{\gamma_k}{M_k}\boldsymbol{I}_{M_k}, \qquad \mathrm{Cov}(\boldsymbol{\beta}_{1,k}, \boldsymbol{\beta}_{2,\ell}) = \mathbf{0} \quad (k \neq \ell).$$

Define

$$t_k = \sum_{j\in S_k} m_j^{(g)}, \qquad L_{k\ell} = \sum_{j\in S_k} \ell^\star_{j\ell}, \qquad g_\ell = \frac{\gamma_\ell}{M_\ell}, \qquad \boldsymbol{M} = (M_1, \dots, M_K)^\top.$$

The aggregate partitioned SUM-CORE moment is

$$\mathbb{E}[t_k] = \sqrt{N_1 N_2}\sum_{\ell=1}^K L_{k\ell}g_\ell + M_k c_{\mathrm{ov}}, \qquad k = 1, \dots, K.$$

Equivalently,

$$\mathbb{E}\left[\frac{\boldsymbol{t} - \boldsymbol{M}c_{\mathrm{ov}}}{\sqrt{N_1N_2}}\right] = \boldsymbol{L}\boldsymbol{g}.$$

In practice, we construct $\widehat{\boldsymbol{L}}$ by replacing $\ell^\star_{j\ell}$ with the precomputed genome-wide LD-score estimates $\widetilde{\ell}_{j\ell}$ and summing over $j \in S_k$. Given $\widehat{c}_{\mathrm{ov}}$, we solve

$$\frac{\boldsymbol{t} - \boldsymbol{M}\widehat{c}_{\mathrm{ov}}}{\sqrt{N_1N_2}} = \widehat{\boldsymbol{L}}\widehat{\boldsymbol{g}}$$

and recover $\widehat{\gamma}_k = M_k\widehat{g}_k$. When $\widehat{\boldsymbol{L}}$ is ill-conditioned, as can occur for overlapping or highly collinear annotations, we solve the system with a small ridge regularization proportional to $\mathrm{tr}(\widehat{\boldsymbol{L}})$.

The population quantity $c_{\mathrm{ov}}$, and its supplied or summary-estimated sample quantity $\widehat{c}_{\mathrm{ov}}$, are the same for every component. We therefore obtain $\widehat{c}_{\mathrm{ov}}$ once per trait pair and reuse it for all partitioned analyses.

#### S1.7 Continuous and overlapping annotations

Let $\boldsymbol{A} \in \mathbb{R}^{M\times K}$ be a real-valued annotation matrix with $A_{jk} \geq 0$ denoting the weight of SNP $j$ in component $k$, and define

$$M_k = \sum_{j=1}^M A_{jk}.$$

For partitioned SUM-CORE, replace bin-wise sums by weighted sums:

$$t_k = \sum_{j=1}^M A_{jk}m_j^{(g)}, \qquad L_{k\ell} = \sum_{j=1}^M A_{jk}\ell^\star_{j\ell}.$$

With $g_\ell = \gamma_\ell/M_\ell$, the aggregate system

$$\mathbb{E}\left[\frac{\boldsymbol{t} - \boldsymbol{M}c_{\mathrm{ov}}}{\sqrt{N_1N_2}}\right] = \boldsymbol{L}\boldsymbol{g}.$$

continues to hold. The partitioned heritability normal equations use the same weighted replacement for hard membership sums.

##### S1.8 Jackknife standard errors

We use a delete-one-block SNP jackknife with $B_{\mathrm{JK}}$ approximately equal-sized genomic blocks. For an estimator $\widehat{\theta}$, let $\widehat{\theta}^{(-b)}$ denote the estimate obtained after deleting block $b$, and let

$$\bar{\theta}_{\mathrm{JK}} = \frac{1}{B_{\mathrm{JK}}} \sum_{b=1}^{B_{\mathrm{JK}}} \widehat{\theta}^{(-b)}.$$

The jackknife variance estimate is

$$\widehat{\mathrm{Var}}_{\mathrm{JK}}(\widehat{\theta}) = \frac{B_{\mathrm{JK}} - 1}{B_{\mathrm{JK}}} \sum_{b=1}^{B_{\mathrm{JK}}} \left( \widehat{\theta}^{(-b)} - \bar{\theta}_{\mathrm{JK}} \right)^2 .$$

For SUM-RHE, each replicate recomputes the HE-scale moment sums and the LD-score aggregate matrix with the deleted SNP block removed. For SUM-CORE in supplied-overlap mode, the main jackknife conditions on the supplied $\widehat{c}_{\mathrm{ov}}$. In summary-only mode, $\widehat{c}_{\mathrm{ov}}$ is re-estimated in each replicate. Derived quantities such as enrichment, $\tau^*$, genome-wide $h^2$, and partitioned $r_g$ are recomputed within each replicate before applying the jackknife formula.

#### S1.9 Simulation details

All simulations used the post-QC, non-MHC genotype data for the corresponding UK Biobank cohort. Phenotypes were generated from standardized genotypes, and each simulation setting was evaluated over 100 replicates unless noted otherwise. For summary-statistic methods, GWAS summary statistics were generated from the simulated phenotypes and analyzed with the same SNP set, LD-score/reference, and annotation pipeline described in the benchmark methods. For individual-level RHE-mc, the simulated phenotypes were analyzed directly with the matched genotype and annotation files. We compared methods to the known simulation truth using bias, mean squared error (MSE), reported standard errors, and calibration of nominal tests where relevant.

**Heritability simulations** For genome-wide heritability simulations, all SNPs were assigned to a single annotation component. We simulated traits with $h_g^2 \in \{0.10, 0.25, 0.40\}$ and causal proportions $p_{\text{causal}} \in \{1, 0.1, 0.01\}$. Causal SNPs were sampled within the genome-wide component, and causal effects were drawn from a Gaussian distribution with equal variance across selected SNPs; the MAF and LD exponents were set to zero. Environmental noise was added so that the phenotypic variance was one.

For the stratified-effects simulations, we added a fixed ancestry component to the standardized $h_g^2 = 0.25$, $p_{\text{causal}} = 1$ phenotypes, $y_{\text{strat}} = \text{scale}(y) + 0.2\,\text{scale}(\text{PC}_1)$, following simulations in Luo et al. (2021). Covariate-adjusted methods used GWAS summary statistics and LD scores adjusted for 10 PCs in EUR/AFR and 20 PCs in SAS, while no-PC comparisons used unadjusted GWAS inputs and LD scores/taggings. Accuracy was summarized from $\hat{h}_g^2 - h_g^2$ and MSE across replicates, with relative MSE computed against the designated baseline method in the corresponding figure.

**Random-SNP omission sensitivity.** We reused the genome-wide heritability simulation phenotypes and GWAS summary statistics for EUR, SAS and AFR under all nine combinations of $h_g^2 \in \{0.10, 0.25, 0.40\}$ and $p_{\text{causal}} \in \{0.01, 0.10, 1.00\}$, with 100 replicates per setting. Within each cohort, we generated one random ordering of SNPs using seed 20260811 and used nested masks to omit 0%, 10%, 25%, or 50% of variants. The same cohort-specific masks were used across simulation settings and replicates. Each mask was applied to both the GWAS summary statistics and LD-score rows by variant intersection; LD-score values for retained variants were not recomputed. Thus, this experiment evaluates reduction of the modeled SNP set rather than surrogate-variant assignment. SUMMIT was fitted using the single-component model with 10 SNP-jackknife blocks. We report the ratio of the mean estimate after omission to the corresponding mean estimate from the complete SNP set. Confidence intervals were obtained from 10,000 paired bootstrap resamples of the simulation replicates.

**Heritability enrichment simulations** For the two-component enrichment benchmark, SNPs were partitioned into `Coding.w.flanking`, defined as the union of the Baseline-LD `Coding_UCSC` and `Coding_UCSC.flanking.500` annotations, and its complement. We varied genome-wide heritability $h_g^2 \in \{0.10, 0.25, 0.40\}$ with $p_{\text{causal}} \in \{1, 0.01\}$, two genetic architectures, and two enrichment scenarios ("weak" or "strong" enrichment of the coding region). The GCTA architecture used equal effect-size variance within each component, whereas the LDAK architecture weighted SNPs by $\text{MAF}^{0.75}\text{LD}^{-1}$. The "weak" enrichment setting multiplied the architecture-specific per-SNP effect-variance weights by 1.5 in the coding component and 0.75 in its complement; the "strong" setting used multipliers of 3.0 and 0.3. Component-level heritability targets were normalized so that they summed to the specified genome-wide heritability after applying the architecture-specific SNP weights. Varying $h_g^2 \in \{0.10, 0.25, 0.40\}$, $p_{\text{causal}} \in \{1, 0.01\}$, two effect-generation schemes, and two enrichment magnitudes gave 24 enriched settings per cohort, with 100 replicates per setting. For discrimination, $z = (\widehat{\text{enrichment}} - 1)/\text{SE}$ was calculated in each replicate. AUROC and AUPR were calculated separately for each enriched setting using null simulations matched for $h_g^2$ and $p_{\text{causal}}$, and the 24 setting-specific values were averaged with equal weight. Because the numbers of finite enriched and null statistics occasionally differed, AUPR was calculated with total weight 0.5 assigned to each class within a setting. For testing calibration of enrichment estimates, we applied the methods to data generated under a single heritability component (so that there is no enrichment) but analyzed using the coding/rest annotation. We evaluated the false-positive rate of the enrichment statistic using the method-reported estimates and standard errors.

**Overlapping-annotation enrichment simulations.** We evaluated enrichment inference using the 16-component overlapping Baseline-LD model listed in Supplementary Table S2. Phenotypes were simulated under an infinitesimal GCTA architecture with genome-wide $h_g^2 = 0.5$. In the null setting, all simulated heritability was assigned to the base component. In the focal alternative, $h^2_{\text{base}} = 0.45$ and $h^2_{\text{focal}} = 0.05$, where the focal component was the ancient-promoter annotation; the remaining annotations were included in the fitted model but were not assigned separate simulation components. The resulting true focal enrichments were 7.406, 7.075, and 8.874 in EUR, SAS, and AFR, respectively, reflecting cohort-specific annotation frequencies. We generated 100 null and 100 focal-alternative replicates per cohort. Power and false-positive rates were calculated from the one-sided enrichment statistic $z = (\widehat{\text{enrichment}} - 1)/\text{SE}$ among finite estimates, with pointwise Wilson confidence intervals.

**MAF–LD simulations** For MAF–LD heritability benchmarks, phenotypes were generated under GCTA or LDAK effect-size models, with genome-wide heritability fixed at $h_g^2 = 0.5$. We selected causal variants from either the low-MAF group, defined as MAF in $[0, 0.1)$, or the higher-MAF group, defined as MAF in $[0.1, 0.5]$. These two groups were used only to define the set of variants that could be causal and were distinct from the six MAF bins used for inference.

Within the selected MAF group, all variants were causal when $p_{\text{causal}} = 1$, whereas variants were selected independently with probability 0.01 when $p_{\text{causal}} = 0.01$. In the GCTA simulations, causal effects had equal expected variance. In the LDAK simulations, causal variants were selected in the same manner, but their effect-size variances were proportional to

$$\frac{\{\text{MAF}_j(1 - \text{MAF}_j)\}^{0.75}}{\ell_j^{(2\,\text{Mb})}},$$

where $\ell_j^{(2\,\text{Mb})}$ is the cohort-specific LD score calculated using a 2-Mb window. The effect-size variances were normalized within each replicate to sum to $h_g^2 = 0.5$. Varying the two effect-size models, two MAF groups, and two causal proportions gave eight settings per cohort, with 100 replicates per setting.

For inference, all methods used the same cohort-specific 24-component MAF–LD annotation, formed by combining six MAF bins with four quartiles of the 2-Mb local LD score. SUMMIT and cov-LDSC used PC-adjusted GWAS summary statistics and matched PC-adjusted LD scores, whereas SumHer-LDAK used unadjusted summary statistics to match its unadjusted SNP taggings. SUMMIT used genome-wide LD scores, cov-LDSC used 20-Mb LD scores, and SumHer-LDAK used 20-Mb taggings. We compared methods using their genome-wide $\widehat{h}_g^2$ estimates.

**MAF–LD partitioned genetic correlation simulations** For the MAF–LD partitioned genetic-correlation simulations reported here, we used a GCTA 8-bin architecture. The annotation combined two MAF bins, $[0, 0.1)$ and $[0.1, 0.5]$, with four LD-score bins. Both traits had constant heritability across bins: $h^2_{1,b} = 0.03125$ and $h^2_{2,b} = 0.075$ for each of the eight bins, giving genome-wide heritabilities of 0.25 and 0.60. All SNPs were causal within each of the eight bins ($p_{\text{causal}} = 1$). For SNP $j$ in bin $b$, the pair of trait effects was drawn jointly with bin-specific variances and covariance divided by the number of SNPs in that bin. We used three bin-specific correlation patterns: `Nullrg`, with $\rho_b = 0$ in every bin; `Constrg`, with $\rho_b = 0.3$ in every bin; and `Gradrg`, with $\rho_b = (0.18, 0.26, 0.34, 0.42, 0.18, 0.26, 0.34, 0.42)$. The first four entries correspond to the four LD-score quartiles in the low-MAF stratum and the final four to the same quartiles in the high-MAF stratum; thus, the LD gradient was repeated in both MAF strata. Environmental correlation was set to zero. The null setting was used to assess calibration, and the constant and gradient settings were used to evaluate recovery of genome-wide and bin-specific genetic-correlation patterns.

For the signed genetic correlation architecture in Figure 3C, both traits retained the same heritability profile as above, with $h^2_{1,b} = 0.03125$ and $h^2_{2,b} = 0.075$ in each bin. We considered two architectures: one in which genetic covariances (and correlations) are positive in the low-MAF and negative in the high MAF bins and the second with these patterns reversed. In the low-MAF-positive architecture, the bin-specific genetic correlations were $\rho_b = (0.8, 0.8, 0.8, 0.8, -0.2, -0.2, -0.2, -0.2)$; in the high-MAF-positive architecture, this profile was reversed. Both architectures therefore had genome-wide $h_1^2 = 0.25$, genome-wide $h_2^2 = 0.60$, genome-wide genetic covariance $\gamma_g = 0.11619$, and genome-wide $r_g = 0.30$. We generated 100 replicates per architecture and cohort, with environmental correlation set to zero.

Figure 3D used the eight-bin `Gradrg` architecture. For each method and replicate, we calculated $\widehat{D} = \sqrt{\widehat{h}_1^2 \widehat{h}_2^2}$ and plotted $\widehat{\gamma}_g$ against $\widehat{D}$. Replicates were included when $\widehat{\gamma}_g$, $\widehat{h}_1^2$, and $\widehat{h}_2^2$ were finite and the estimated heritability product was positive. No additional filtering by estimate magnitude was applied.

### S2 Supplementary Tables

#### S2.1 Bias in genome-wide heritability simulation benchmarks

Table S1: **Relative bias in the MAF–LD heritability simulations.** Within each simulation setting, replicate-level relative error was defined as $r_i = (\widehat{h}^2_{g,i} - h^2_g)/h^2_g$, and signed relative bias was calculated as $\bar{r}$ across 100 replicates. The table reports the median and range of $|\bar{r}|$ across eight architectures for each method and cohort: two effect-size models, two causal-MAF supports, and two causal proportions. Bias was tested separately in each architecture using a two-sided one-sample $t$-test of the 100 replicate-level relative errors (with significance defined as $p < 0.05/(8 \times 3)$, corresponding to Bonferroni correction over the eight settings and three cohorts). The positive- and negative-bias columns partition the significant settings according to the sign of $\bar{r}$.

| Cohort | Method | Median absolute relative bias (%) | Range (%) | Significant settings (of 8) | Positive bias | Negative bias |
|---|---|---|---|---|---|---|
| EUR | SUMMIT | 1.56 | [0.47, 2.49] | 0/8 | 0 | 0 |
| EUR | cov-LDSC | 29.23 | [3.42, 62.29] | 4/8 | 0 | 4 |
| EUR | SumHer-LDAK | 2.31 | [0.26, 4.18] | 3/8 | 3 | 0 |
| SAS | SUMMIT | 1.47 | [0.01, 3.20] | 0/8 | 0 | 0 |
| SAS | cov-LDSC | 26.41 | [1.73, 62.64] | 3/8 | 0 | 3 |
| SAS | SumHer-LDAK | 208.28 | [145.03, 294.29] | 8/8 | 8 | 0 |
| AFR | SUMMIT | 4.69 | [0.36, 9.28] | 0/8 | 0 | 0 |
| AFR | cov-LDSC | 9.06 | [4.07, 85.52] | 3/8 | 2 | 1 |
| AFR | SumHer-LDAK | 927.08 | [626.22, 1762.55] | 8/8 | 8 | 0 |

#### S2.2 Overlapping annotations enrichment simulations

Table S2: **Annotations used in the overlapping-annotation enrichment simulation.** The fitted model contained the base annotation, the focal ancient-promoter annotation, and 14 additional regulatory and gene-model annotations.

| Component | Annotation |
|---|---|
| 1 | base |
| 2 | Ancient_Sequence_Age_Human_Promoter_w_flanking |
| 3 | Human_Promoter_Villar_w_flanking |
| 4 | TSS_Hoffman_w_flanking |
| 5 | Human_Promoter_Villar_ExAC_w_flanking |
| 6 | Promoter_UCSC_w_flanking |
| 7 | PromoterFlanking_Hoffman_w_flanking |
| 8 | BivFlnk_w_flanking |
| 9 | UTR_3_UCSC_w_flanking |
| 10 | H3K4me3_peaks_Trynka |
| 11 | H3K9ac_peaks_Trynka |
| 12 | H3K9ac_Trynka_w_flanking |
| 13 | H3K4me3_Trynka_w_flanking |
| 14 | Coding_UCSC_w_flanking |
| 15 | H3K27ac_PGC2_w_flanking |
| 16 | Enhancer_Hoffman_w_flanking |

#### S2.3 Runtime and memory measurement

Table S3: **Runtime of reference-statistic construction and downstream heritability estimation.** Wall-clock times are in seconds. Reference-statistic construction is performed once per cohort and SNP set. For SUMMIT, the projection dimensions were $B = 1{,}024$ for all the cohorts. The asterisk denotes that cov-LDSC EUR (300k) was run on the EUR-50k subset ($N = 50{,}112$), rather than the full EUR (300k) LD sample due to memory and runtime constraints.

| | | | Reference statistic | | | Estimation | | |
|---|---|---|---|---|---|---|---|---|
| Population | $N$ | $M$ | SUMMIT | cov-LDSC | SumHer LDAK | SUMMIT | cov-LDSC | SumHer LDAK |
| EUR (300k) | 291,273 | 454,207 | 967 | 18,192* | 6,232 | 3.7 | 4.4 | 3.9 |
| EUR | 10,060 | 454,207 | 70 | 8,068 | 813 | 3.7 | 4.3 | 3.5 |
| SAS | 7,912 | 448,403 | 55 | 6,338 | 657 | 4.2 | 3.5 | 3.6 |
| AFR | 3,416 | 1,002,034 | 89 | 11,982 | 1,148 | 7.3 | 7.8 | 8.6 |

Table S4: **Memory usage during reference-statistic construction.** Values are peak sampled resident set size in GB for single-component reference-statistic construction at default settings. SUMMIT values were measured using the projection dimensions in Table S3. For cov-LDSC, the EUR (300k) asterisk denotes the run on the EUR-50k subset ($N = 50{,}112$).

| Population | $N$ | $M$ | SUMMIT | cov-LDSC | SumHer LDAK |
|---|---|---|---|---|---|
| EUR (300k) | 291,273 | 454,207 | 12.0 | 13.2* | 18.4 |
| EUR | 10,060 | 454,207 | 1.8 | 2.3 | 1.6 |
| SAS | 7,912 | 448,403 | 1.5 | 1.8 | 1.5 |
| AFR | 3,416 | 1,002,034 | 1.3 | 1.9 | 3.8 |

#### S2.4 Total heritability comparisons

Table S5: **Population-differential heritability hits.** Bonferroni-significant traits and adjusted $p$ values, ordered from most to least significant within each population pair, method and annotation model. Adjustment uses 240 tests per method and model (40 traits $\times$ 6 population pairs).

| Population pair | Method | Bins | Bonferroni-significant traits (adjusted $p$ value) |
|---|---|---|---|
| AFR – SAS | SumHer-GCTA | 8 | HT(0.00708) |
| AFR – SAS | SumHer-GCTA | 24 | HT($3.01 \times 10^{-4}$) |
| AFR – SAS | SumHer-LDAK | 24 | HT($4.36 \times 10^{-4}$) |
| EUR – SAS | SUMMIT | 8 | EA($7.58 \times 10^{-4}$), SMK(0.00548) |
| EUR – SAS | SUMMIT | 24 | EA($9.72 \times 10^{-4}$), SMK(0.00706) |
| EUR – SAS | SumHer-GCTA | 8 | HT($5.14 \times 10^{-7}$), BMD($1.69 \times 10^{-5}$), SMK($5.77 \times 10^{-5}$), BMI(0.00288) |
| EUR – SAS | SumHer-GCTA | 24 | HT($2.94 \times 10^{-7}$), BMD($8.47 \times 10^{-6}$), SMK($8.26 \times 10^{-5}$), BMI(0.00282) |
| EUR – SAS | SumHer-LDAK | 8 | BMD($3.63 \times 10^{-4}$), SMK(0.00331), HT(0.0173) |
| EUR – SAS | SumHer-LDAK | 24 | HT($7.76 \times 10^{-7}$), BMD($7.73 \times 10^{-6}$), SMK($1.53 \times 10^{-4}$), BMI(0.0067) |
| EUR (300k) – SAS | SUMMIT | 8 | EA($2.89 \times 10^{-9}$), SMK(0.0289) |
| EUR (300k) – SAS | SUMMIT | 24 | EA($4.67 \times 10^{-9}$), SMK(0.0403) |
| EUR (300k) – SAS | SumHer-GCTA | 8 | HT($6.39 \times 10^{-13}$), BMD($4.89 \times 10^{-7}$), BMI($1.26 \times 10^{-6}$), SMK($3.61 \times 10^{-6}$), EA($1.92 \times 10^{-5}$), SLP(0.0405) |
| EUR (300k) – SAS | SumHer-GCTA | 24 | HT($3.73 \times 10^{-13}$), BMD($2.13 \times 10^{-7}$), BMI($1.27 \times 10^{-6}$), SMK($4.98 \times 10^{-6}$), EA($9.98 \times 10^{-6}$), SLP(0.0334) |
| EUR (300k) – SAS | SumHer-LDAK | 8 | BMD($2.06 \times 10^{-5}$), HT($3.22 \times 10^{-5}$), SMK(0.00208), BMI(0.0187), EA(0.0261) |
| EUR (300k) – SAS | SumHer-LDAK | 24 | HT($3.84 \times 10^{-12}$), BMD($1.90 \times 10^{-7}$), BMI($4.76 \times 10^{-6}$), SMK($1.84 \times 10^{-5}$), EA($2.10 \times 10^{-5}$), SLP(0.03) |

#### S2.5 External GWAS resources used for validation

We evaluated external GWAS summary statistics spanning European, South Asian, and admixed African-ancestry analyses. All external files used in SUMMIT contained effect estimates and standard errors and were converted to a common format with columns `SNP`, `A1`, `A2`, `N`, `BETA`, `SE`, and `Z`. For each local UKB reference cohort, external variants were restricted to SNPs overlapping the corresponding local genotype panel with allele-compatible effect alleles.

Table S6: **External GWAS summary statistics used for external validation.** For each external source, we report the external analysis set, traits included, sample-size range in the harmonized SUMMIT inputs, downloaded variant counts, and allele-compatible overlap with the matched UKB genotype reference.

| Source | Matched UKB reference | External analysis set | Traits used | Sample size in harmonized files | Variants | Model/QC notes |
|---|---|---|---|---|---|---|
| FinnGen R12 (Kurki et al. 2023) | UKB EUR (300k) | Finnish biobank participants linked to nationwide health registries. | Hyperlipoproteinemia, hypertension, ischemic heart disease, T2D. | 479,432–500,348 | 21.33M downloaded per trait; 450,271 overlap variants. | Disease definitions based on Finnish health registry records. |
| GLGC 2021 no-UKB EUR (Graham et al. 2021) | UKB EUR (300k) | European ancestry-specific lipid meta-analysis with UKB cohorts removed. | HDL-C, LDL-C, total cholesterol, log triglycerides, non-HDL-C. | Median $N$: 562,102–922,058 | 29.89–37.01M downloaded; 454,202–454,207 overlap variants. | Preferred GLGC EUR release for analyses where UKB sample overlap matters. |
| GLGC 2021 SAS, all cohorts (Graham et al. 2021) | UKB SAS | South Asian ancestry-specific lipid meta-analysis, all available cohorts. | HDL-C, LDL-C, total cholesterol, log triglycerides, non-HDL-C. | Median $N$: 38,314–40,962 | 16.50–16.89M downloaded; 440,152–441,429 overlap variants. | Potential UKB sample overlap; retained for diagnostic analyses or analyses where overlap is acceptable. |
| GLGC 2021 no-UKB SAS (Graham et al. 2021) | UKB SAS | South Asian ancestry-specific lipid meta-analysis with UKB cohorts removed. | HDL-C, LDL-C, total cholesterol, log triglycerides, non-HDL-C. | Median $N$: 32,096–34,135 | 15.13–15.40M downloaded; 438,944–439,628 overlap variants. | Preferred GLGC SAS release for genetic-correlation analyses against UKB SAS. |
| GLGC 2021 no-UKB AFR (Graham et al. 2021) | UKB AFR | African ancestry-specific lipid meta-analysis with UKB cohorts removed. | HDL-C, LDL-C, total cholesterol, log triglycerides, non-HDL-C. | Median $N$: 23,248–92,554 | 26.68–31.92M downloaded; 1,000,648–1,000,699 overlap variants. | Broadly ancestry-matched, but external AFR cohorts may differ from UKB AFR in admixture, ascertainment and environment. |
| MVP R4 GIA (Gaziano et al. 2016; Verma et al. 2024) | UKB EUR (300k), UKB AFR | U.S. Veterans assigned to genetically inferred ancestry groups. | HbA1c, ALT, AST, glucose, hypertension phecode, ischemic heart disease phecode. | EUR: 338,848–428,138; AFR: 98,735–118,233 | EUR: 19.71–19.74M downloaded, 451,549 overlap; AFR: 36.96–36.97M downloaded, 997,865 overlap. | MVP Release 4 GWAS atlas; GIA denotes genetically inferred ancestry grouping. |
| Genes & Health v010 quantitative GWAS (Finer et al. 2020; Jacobs et al. 2024) | UKB SAS | British Bangladeshi and British Pakistani adults linked to NHS electronic health records. | Median ALT, AST, GGT and HbA1c. | 15,759–40,044 | 30.21–37.59M downloaded; 433,361–433,362 overlap variants. | Quantitative blood-test GWAS from Genes & Health; phenotypes were derived from routinely acquired EHR blood tests. |
| T2DGGI 2024 (Suzuki et al. 2024) | UKB EUR (300k), UKB SAS, UKB AFR | Ancestry-specific T2D summary statistics from a global multi-ancestry meta-analysis. | T2D. | EUR $N_{\mathrm{eff}}$: 1,812,013; SAS median $N_{\mathrm{eff}}$: 50,599; AFR median $N_{\mathrm{eff}}$: 152,337 | EUR: 19.30M downloaded, 449,207 overlap; SAS: 14.48M downloaded, 442,146 overlap; AFR: 19.26M downloaded, 999,840 overlap. | Binary trait analyzed using effective sample size in SUMMIT; SAS T2DGGI was excluded from genetic-correlation figures because of likely UKB overlap. |

*Notes.* Sample size is the $N$ or $N_{\mathrm{eff}}$ column in the harmonized SUMMIT-formatted files. For meta-analysis files with SNP-varying $N$, ranges summarize the median $N$ across retained overlap variants for each trait. Downloaded variant counts exclude file headers. Overlap variants are allele-compatible variants retained after intersecting each external GWAS with the matched UKB genotype reference.

### S3 Supplementary Figures

#### S3.1 Population-specific SNP sets, allele frequencies and PC sensitivity

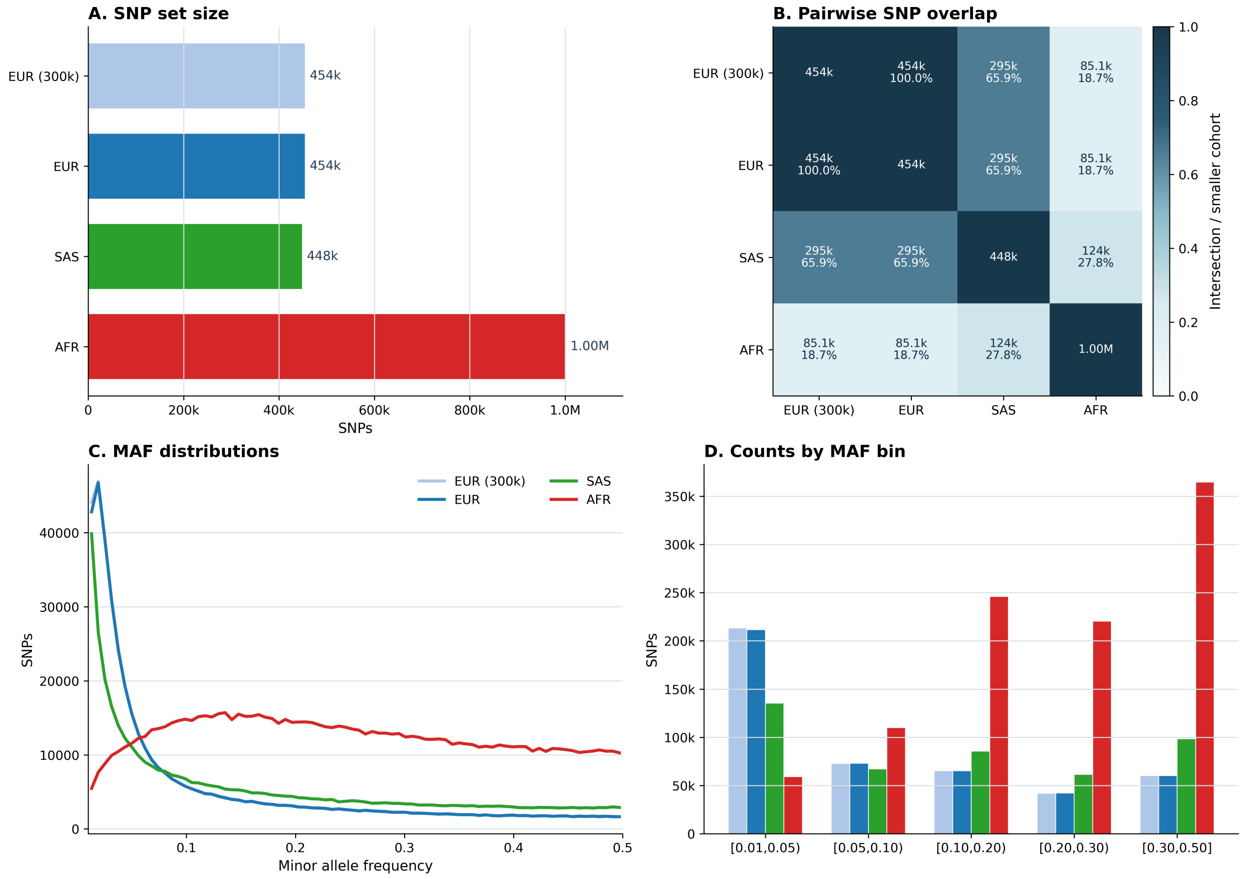

Figure S1: **Population-specific SNP-set overlap and allele-frequency distributions.** The QC-passing common-SNP sets are summarized for each cohort. **Panel A:** Total number of common variants that pass QC for each cohort. EUR (300k) and EUR SNP sets are identical. **Panel B:** Number of overlapping SNPs for each pair of cohorts. For AFR, SNP counts are reported after conversion to GRCh37 coordinates. **Panel C:** MAF distribution for each cohort. **Panel D:** SNP count for each MAF bin. Despite having the largest variant set, AFR cohort has the smallest number of low-MAF variants followed by SAS, given the UKB array and imputed SNPs are focused on EUR samples.

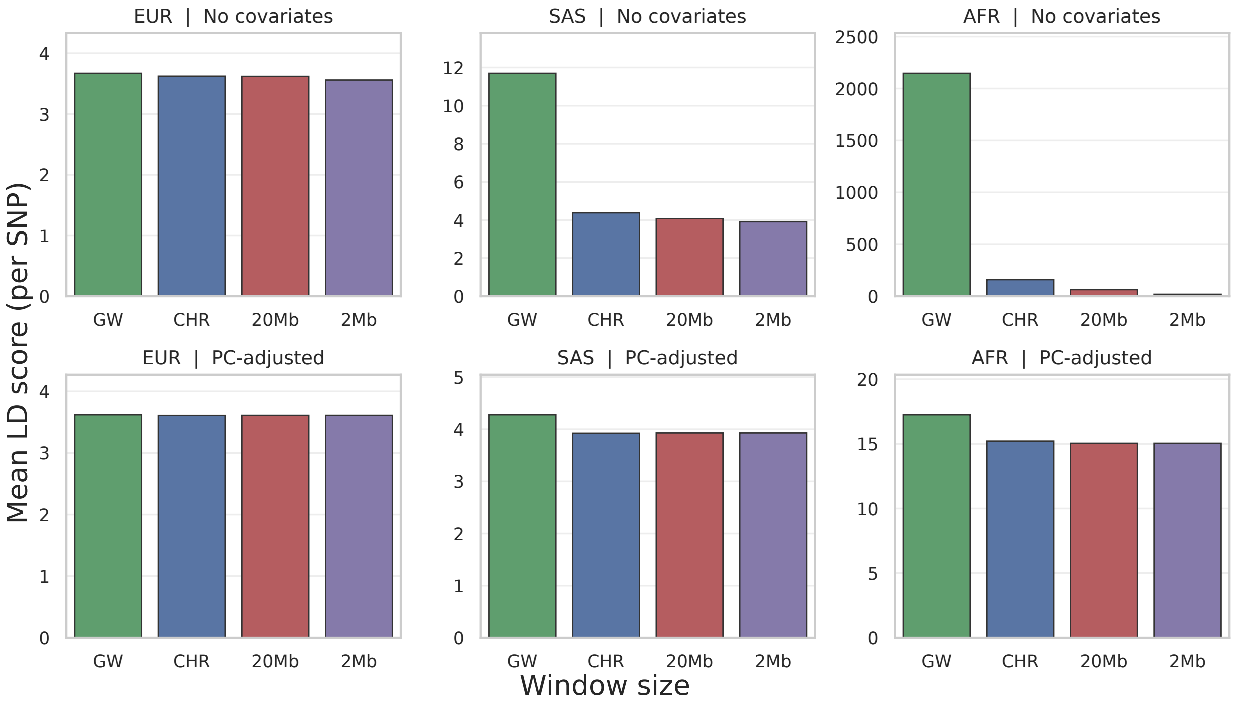

Figure S2: **Average LD scores of each cohort** Mean LD scores for each population with varying window sizes without genotype-PC adjustment (top) and after genotype-PC adjustment (bottom).

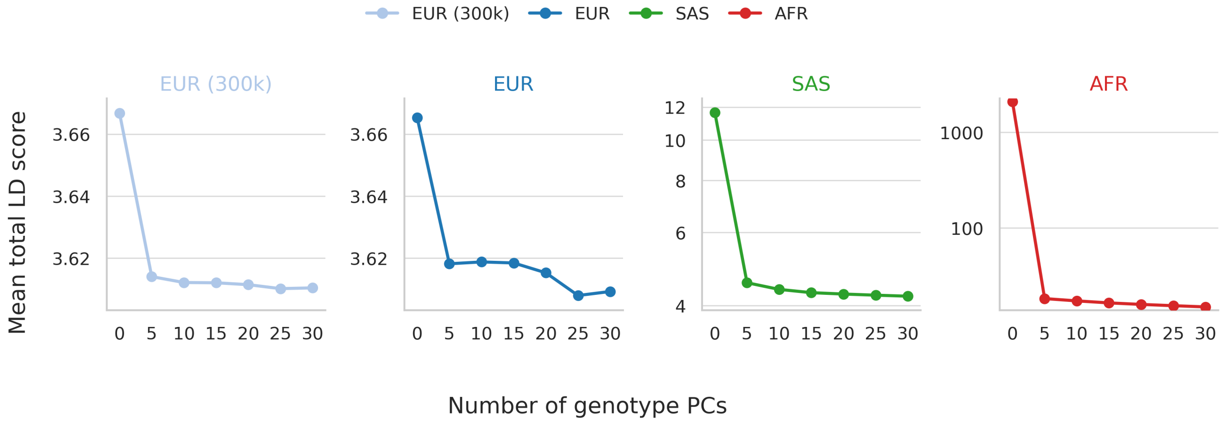

Figure S3: **Sensitivity of SUMMIT genome-wide LD score estimates to genotype PC adjustment.** Mean LD scores for each population as more genotype PCs are included as covariates. EUR results are shown in linear scale, whereas SAS and AFR results are shown in log scale.

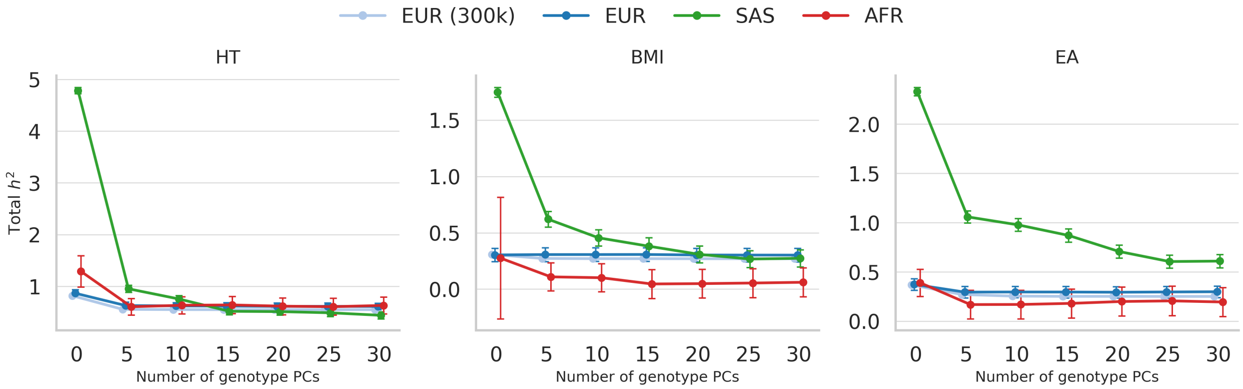

Figure S4: **Sensitivity of SUMMIT genome-wide $h^2$ estimates to genotype PC adjustment.** Genome-wide $h_g^2$ estimates for height (HT), BMI, and educational attainment (EA) with varying numbers of genotype PCs as covariates. All analyses used the same eight-component MAF–LD model (two MAF bins × four LD bins). Within each setting, the genome-wide LD scores and GWAS summary statistics were adjusted for the same covariates. Points and vertical bars show estimates and standard errors.

#### S3.2 Simulation benchmarks: genome-wide and partitioned heritability

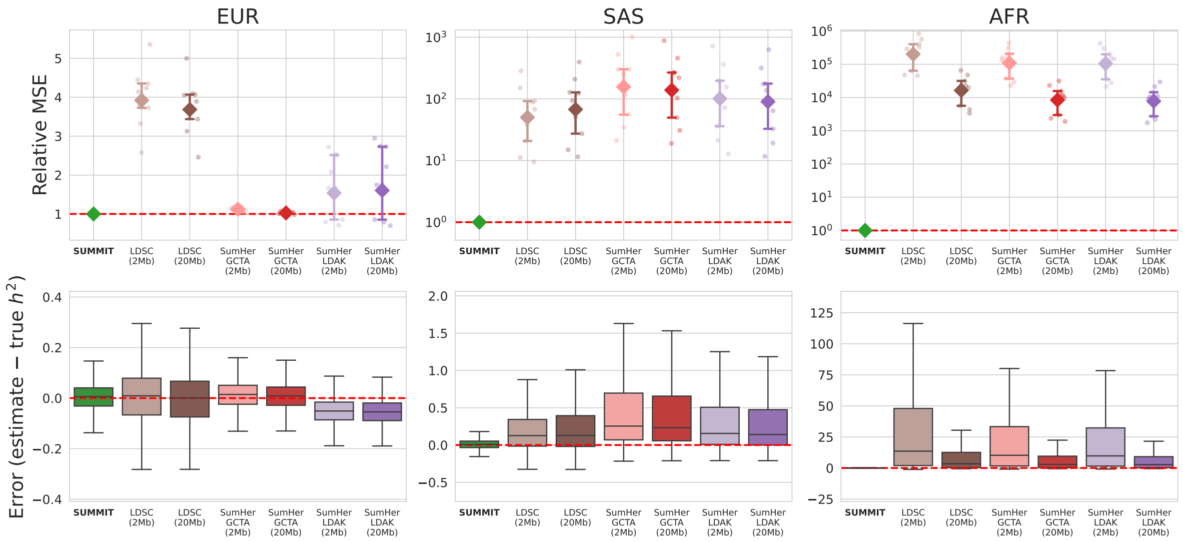

Figure S5: **Window-size sensitivity in single-component genome-wide heritability simulations.** We show how window sizes affect MSE and error for fixed-windowed methods in EUR, SAS and AFR across nine settings per cohort (single-component, defined by $h_g^2 \in \{0.10, 0.25, 0.40\}$ and $p_{\text{causal}} \in \{1.0, 0.10, 0.01\}$). The top row shows MSE relative to SUMMIT for LDSC, SumHer-GCTA, and SumHer-LDAK using 2- and 20-Mb windows. Faint points denote individual settings, diamonds denote geometric means across the nine settings, and bars denote interquartile ranges. The bottom row shows boxplots of finite replicate-level signed errors, $\widehat{h}_g^2 - h_g^2$, pooled over the nine settings, with up to 900 estimates per method and window. In EUR, where the LD is much more local, we see that increasing window sizes do not meaningfully change results for those methods. However in SAS and AFR, window sizes improve performance as it better captures long-range LD. We use the largest window (20Mb) for the main text results.

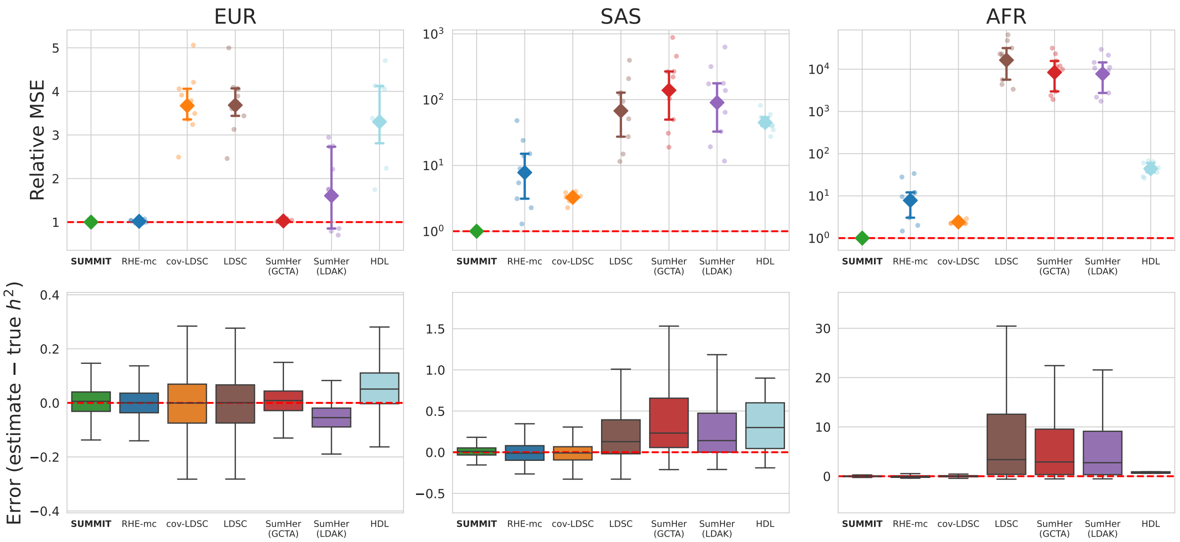

Figure S6: **Full single-component genome-wide heritability benchmark.** MSE relative to SUMMIT and error distributions are shown across the nine single-component, genome-wide simulation settings in EUR, SAS and AFR. In the top row, each faint point is the MSE from one setting relative to SUMMIT; diamonds show geometric means and bars show interquartile ranges across the nine settings. The bottom row shows boxplots of finite replicate-level signed errors, $\widehat{h}_g^2 - h_g^2$, pooled over the nine settings, with up to 900 estimates per method and cohort. cov-LDSC, LDSC, SumHer-GCTA, and SumHer-LDAK used 20-Mb LD summaries.

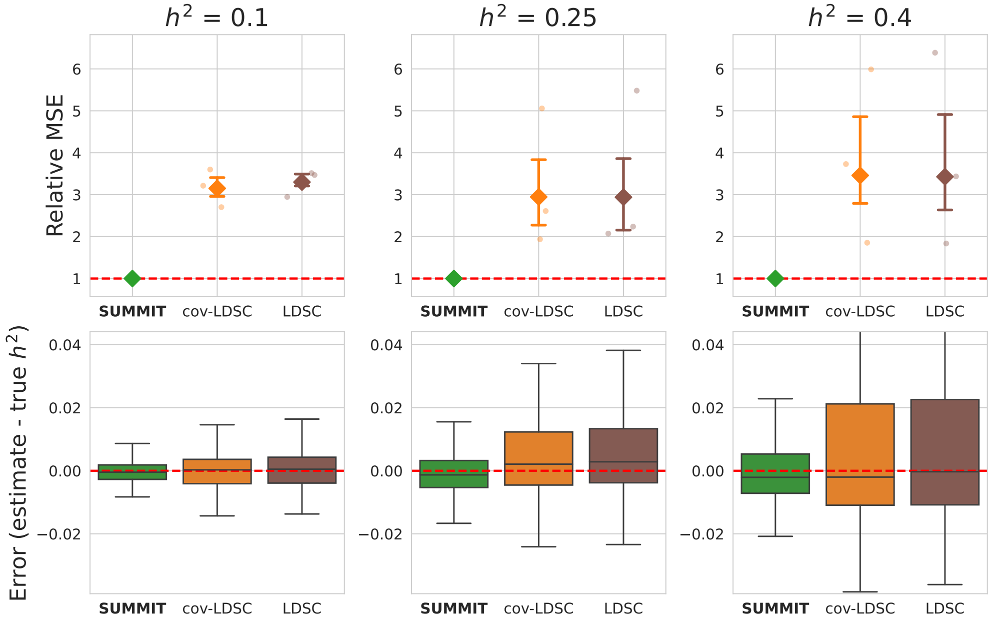

Figure S7: **Genome-wide heritability estimation benchmark with EUR (300k).** MSE relative to SUMMIT and signed estimation-error distributions are shown for the large EUR cohort. As in the small-cohort simulations, the benchmark includes nine settings defined by $h^2 \in \{0.1, 0.25, 0.4\}$ and causal proportion $p \in \{1.0, 0.1, 0.01\}$.

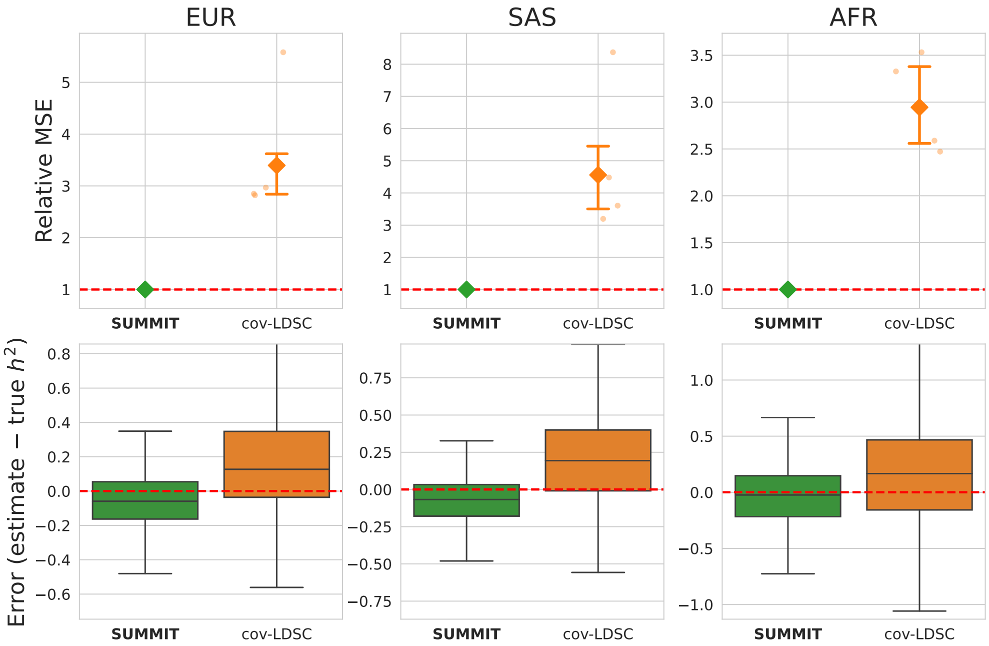

Figure S8: **Genome-wide heritability estimation benchmark for binary traits.** MSE relative to SUMMIT and signed estimation-error distributions are shown for binary trait simulations with liability-scale heritability $h^2_{\text{liability}} = 0.25$ and causal proportions $p \in \{1.0, 0.1, 0.01\}$.

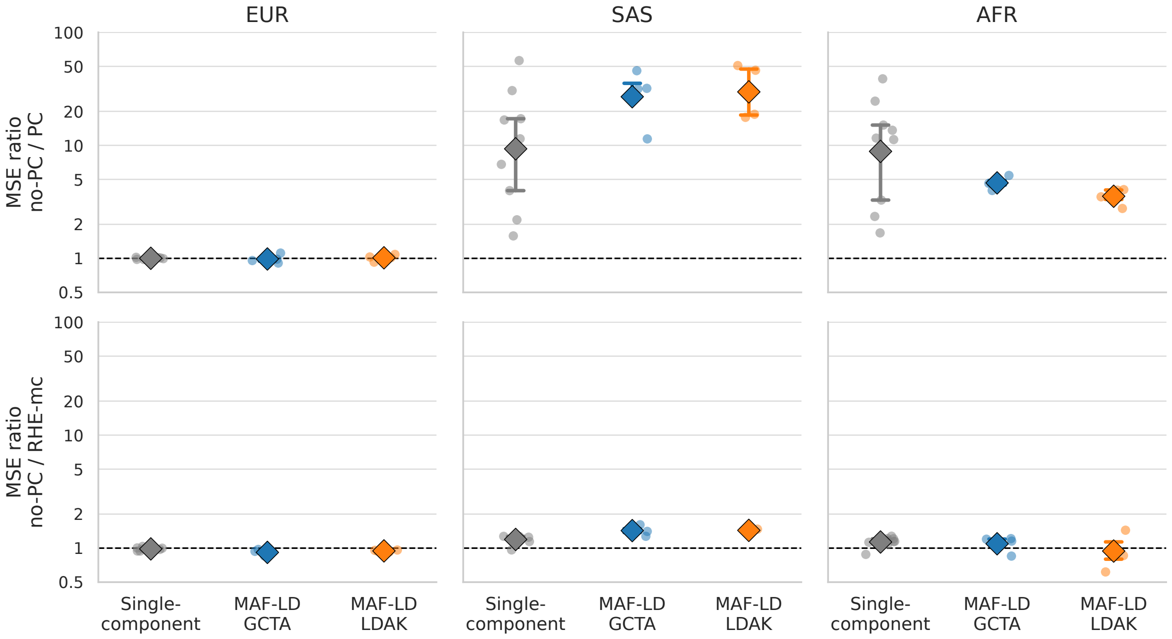

Figure S9: **Effect of covariate residualization on SUMMIT estimates.** Genome-wide heritability simulation benchmarks are compared with and without genotype-PC residualization. Points show the ratio of mean squared error, diamonds denote geometric means, and the dashed line marks equal error. Ratios are computed from matched finite replicate pairs. Simulations do not have any PC-modulated effects. **Top:** Ratio between SUMMIT (no PC-residualization) vs SUMMIT (with PC-residualization). The two are identical in EUR, but in structured or admixed cohorts (SAS and AFR), there is a meaningful difference in performance, as global ancestry dominates both genotypic and phenotypic variances. **Bottom:** Ratio between SUMMIT (no PC-residualization) vs RHE-mc (no PC-residualization). We see that the two perform very similarly, which is expected from Jeong et al. (2024) (showed in EUR cohorts), though there is a small difference in SAS cohorts.

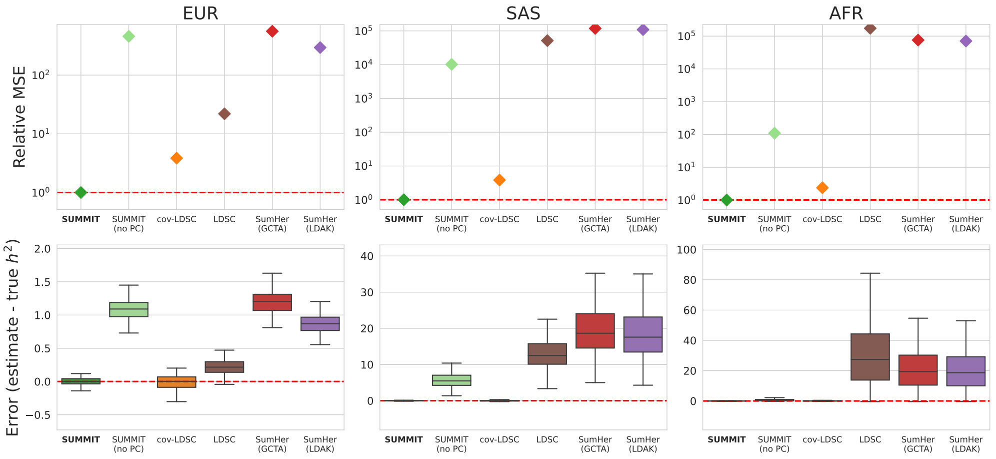

Figure S10: **Genome-wide heritability estimation with PC-modulated effects.** We repeat the benchmark ($h_g^2 = 0.25$ and $p = 1.0$) by adding environmental component, which is aligned with PC1 with variance 0.04, similar to the simulations in Luo et al. (2021). SUMMIT and cov-LDSC used summary statistics and LD scores that use top 10 (EUR and AFR) or 20 (SAS) genotype PCs as covariates (as done in the initial benchmarks), whereas remaining methods used inputs without any PC residualization.

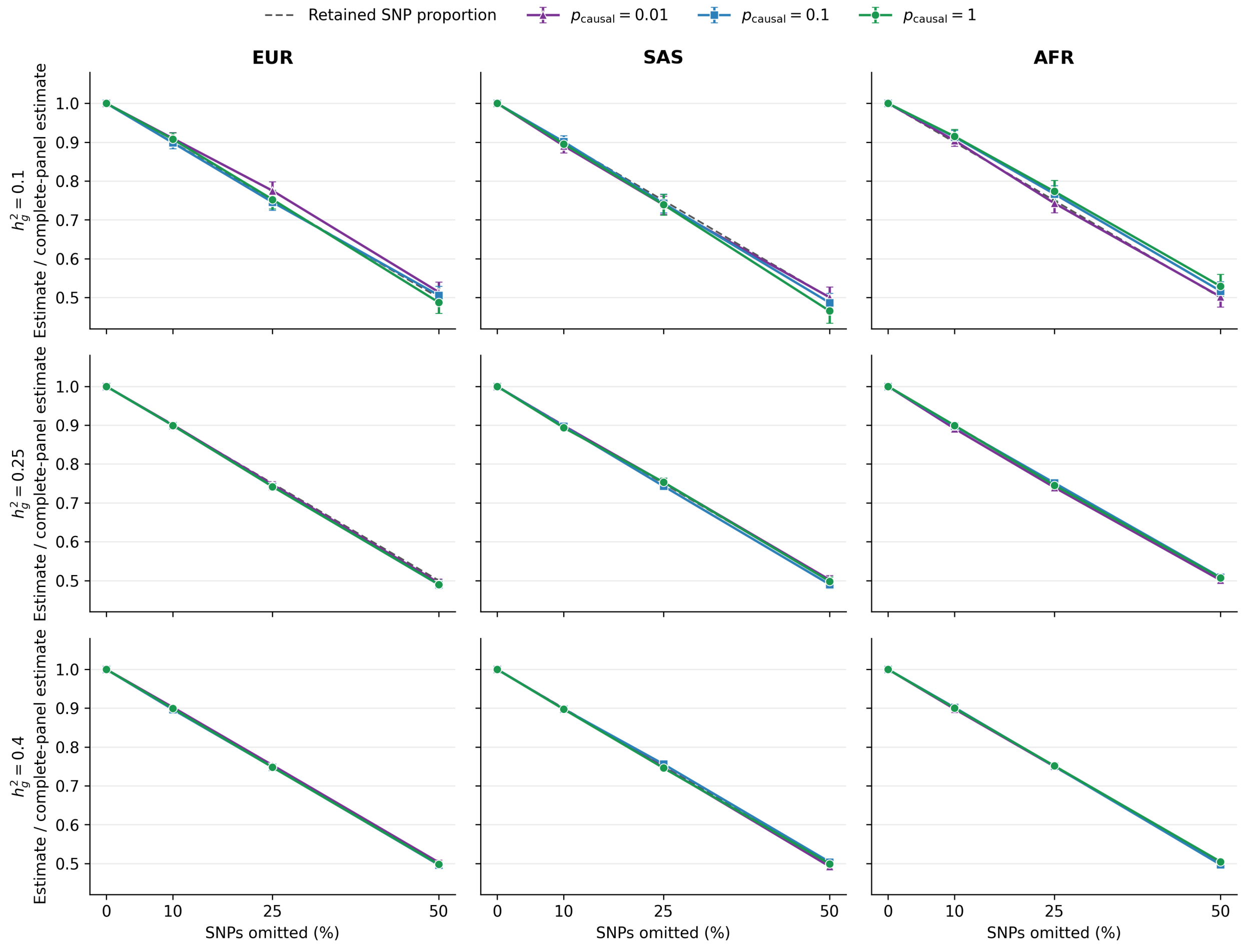

Figure S11: **Sensitivity of SUMMIT genome-wide heritability estimates to random SNP omission.** The same nested random SNP mask was applied within each cohort across 100 simulation replicates and all nine combinations of genome-wide heritability and causal proportion. Rows show genome-wide $h_g^2$, columns show cohorts, and colors and symbols show $p_{\text{causal}}$. The vertical axis is the ratio of the mean SUMMIT estimate after omission to the mean estimate from the complete SNP set in the same setting; error bars are 95% paired-bootstrap confidence intervals from 10,000 resamples. The dashed line is the proportion of SNPs retained, $1 - f$. LD-score values for retained rows were not recomputed. This analysis reduces the modeled SNP set and does not use surrogate-variant assignment.

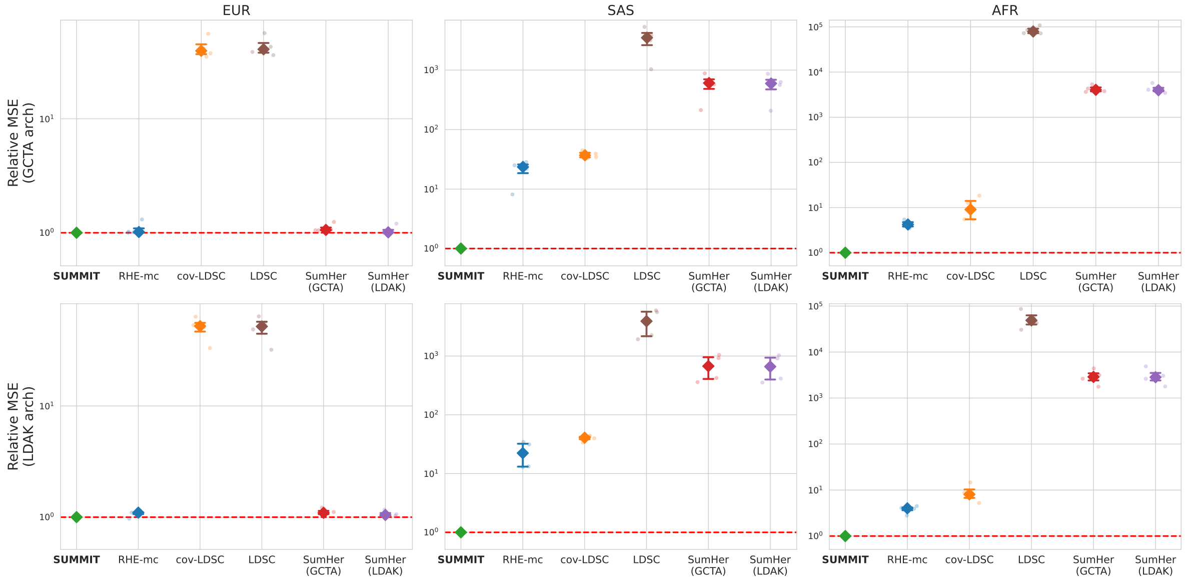

Figure S12: **MAF–LD heritability simulation: relative MSE.** Relative MSE of genome-wide $h_g^2$ estimates is shown under GCTA and LDAK generative architectures across EUR, SAS, and AFR. Within each architecture, four settings combine two causal-MAF supports (MAF $< 0.1$ or MAF $\geq 0.1$) and two causal proportions ($p_{\text{causal}} = 1.0$ or $0.01$). Each faint point denotes one setting; diamonds show medians and bars show interquartile ranges across the four settings. MSE is expressed relative to SUMMIT within each setting.

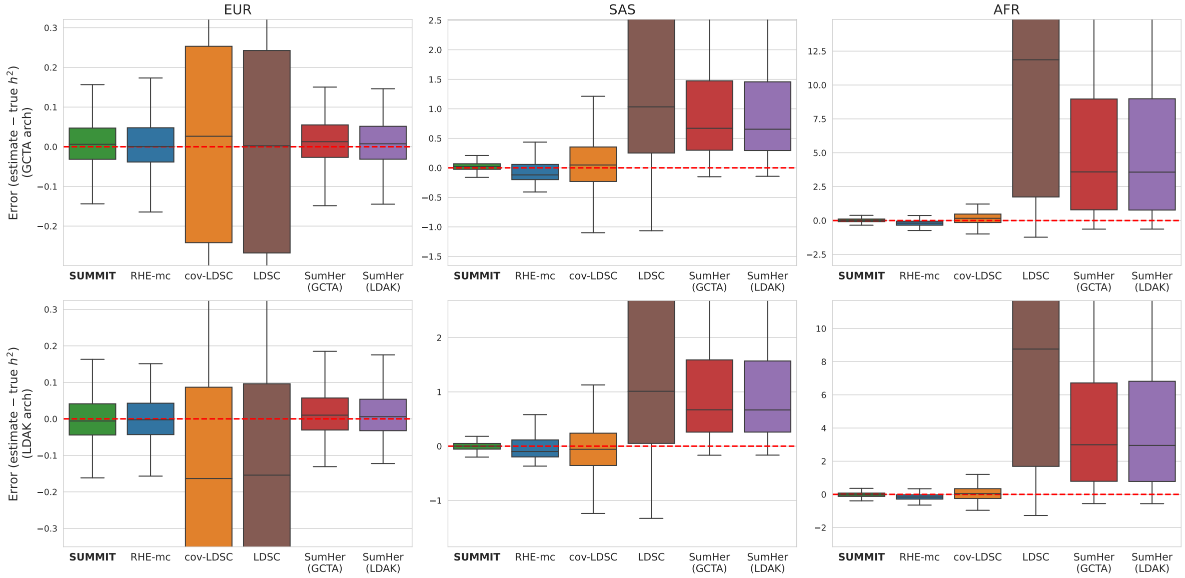

Figure S13: **MAF–LD heritability simulation: error.** Replicate-level errors in genome-wide $h_g^2$, $\widehat{h}_g^2 - h_g^2$, are shown under GCTA and LDAK generative architectures across EUR, SAS, and AFR. Within each architecture, boxes pool 400 estimates per method across four simulation settings and 100 replicates per setting. Boxes show medians and interquartile ranges, whiskers extend to 1.5 interquartile ranges, and observations beyond the whiskers are shown as points.

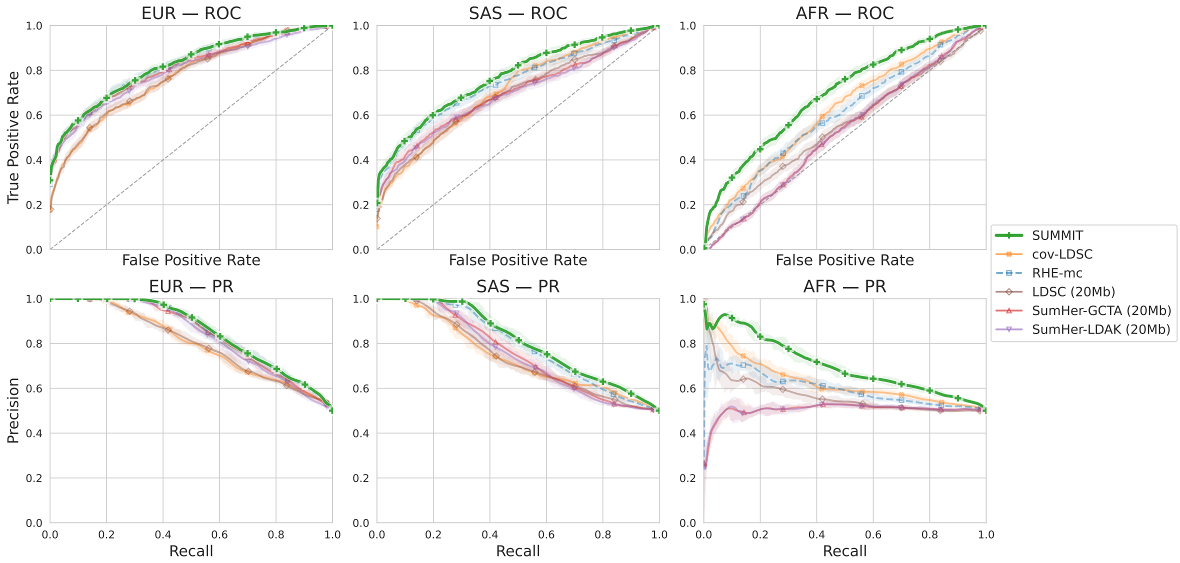

Figure S14: **Functional annotation simulation: enrichment classification.** ROC and precision–recall curves are shown for distinguishing coding-enriched simulations from matched null simulations. The pooled enriched/null set is balanced within simulation strata so that discrimination reflects coding-enrichment recovery rather than class imbalance.

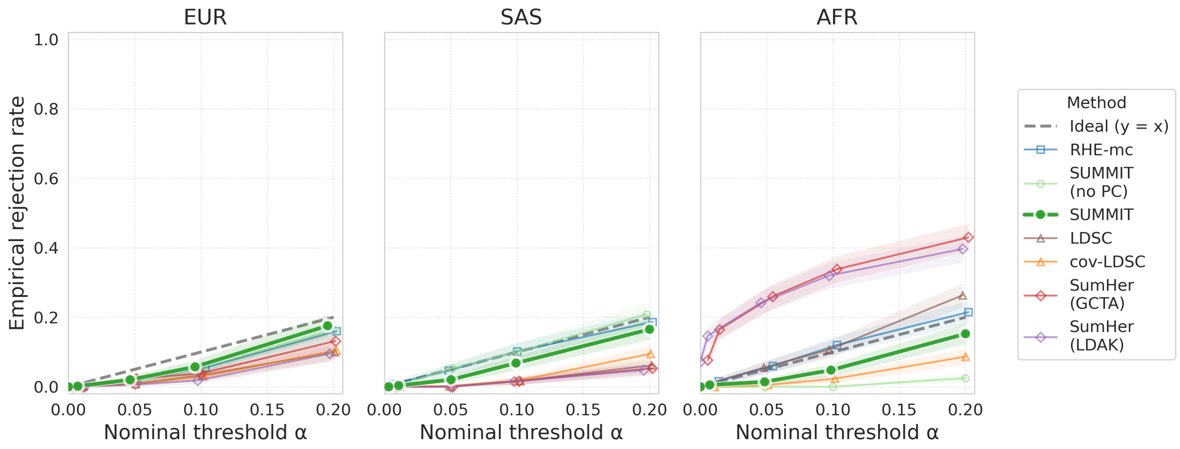

Figure S15: **Functional annotation simulation: null calibration.** Empirical rejection rates are shown under simulations with no coding enrichment for the one-sided test of coding enrichment.

#### S3.3 Simulation benchmarks: genetic correlation

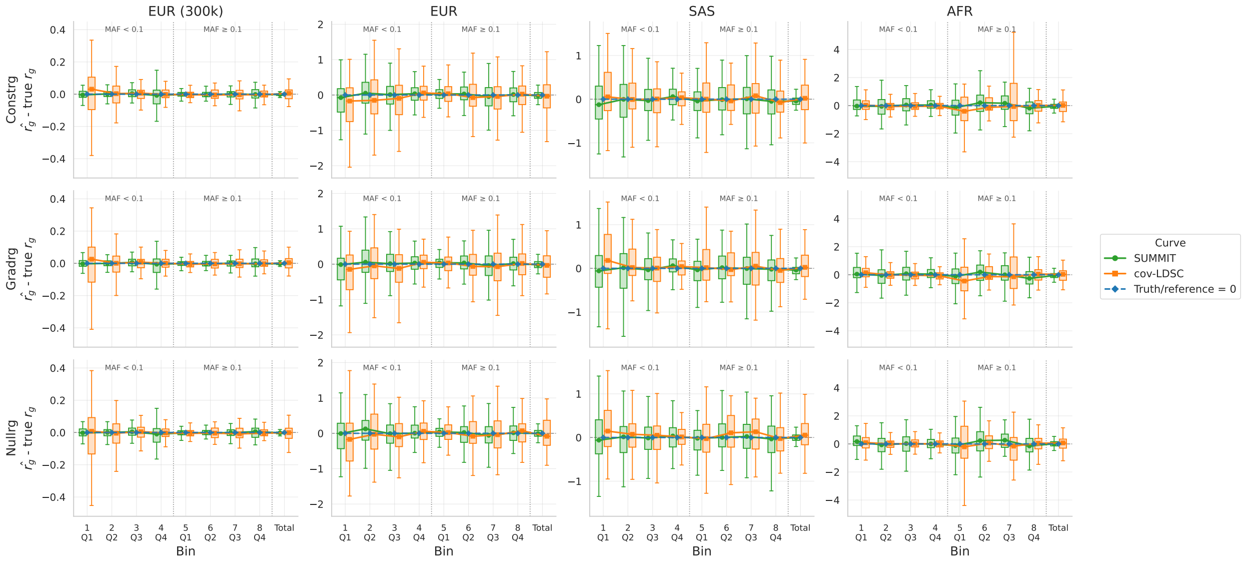

Figure S16: **MAF–LD partitioned genetic-correlation simulation (per-bin error).** Bin-level and genome-wide $r_g$ estimation errors are shown under constant (**top**), gradient (**middle**) and null (**bottom**) $r_g$ architectures.

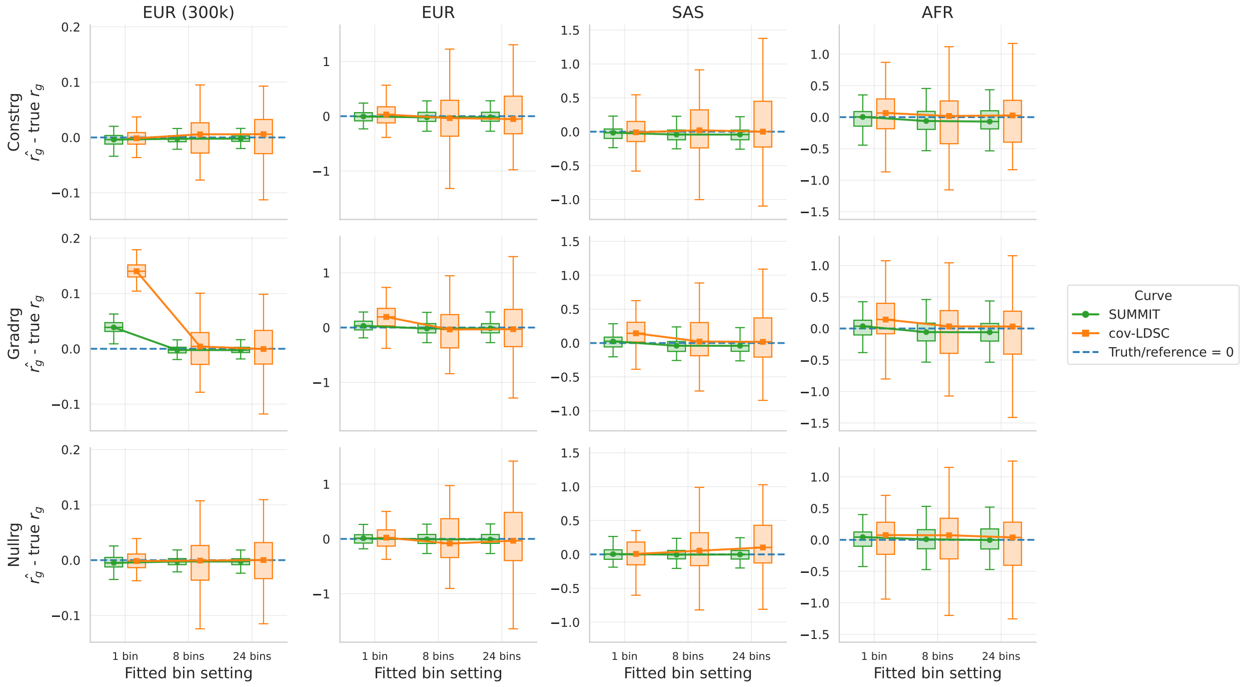

Figure S17: **MAF–LD partitioned genetic-correlation simulation: genome-wide $r_g$ error under different inference models.** Total $r_g$ estimation error is shown when inference uses one bin, 8 MAF–LD bins or 24 MAF–LD bins, under constant (**top**), gradient (**middle**) and null (**bottom**) $r_g$ architectures. When $r_g$ is no longer constant across bins, we find that both SUMMIT and cov-LDSC can yield biased genome-wide $r_g$ estimates when unpartitioned (1 bin) model is used for inference, whereas MAF–LD partitioned models yield unbiased estimates. We therefore use MAF–LD partitioning to estimate the genome-wide $r_g$ in real-data analysis.

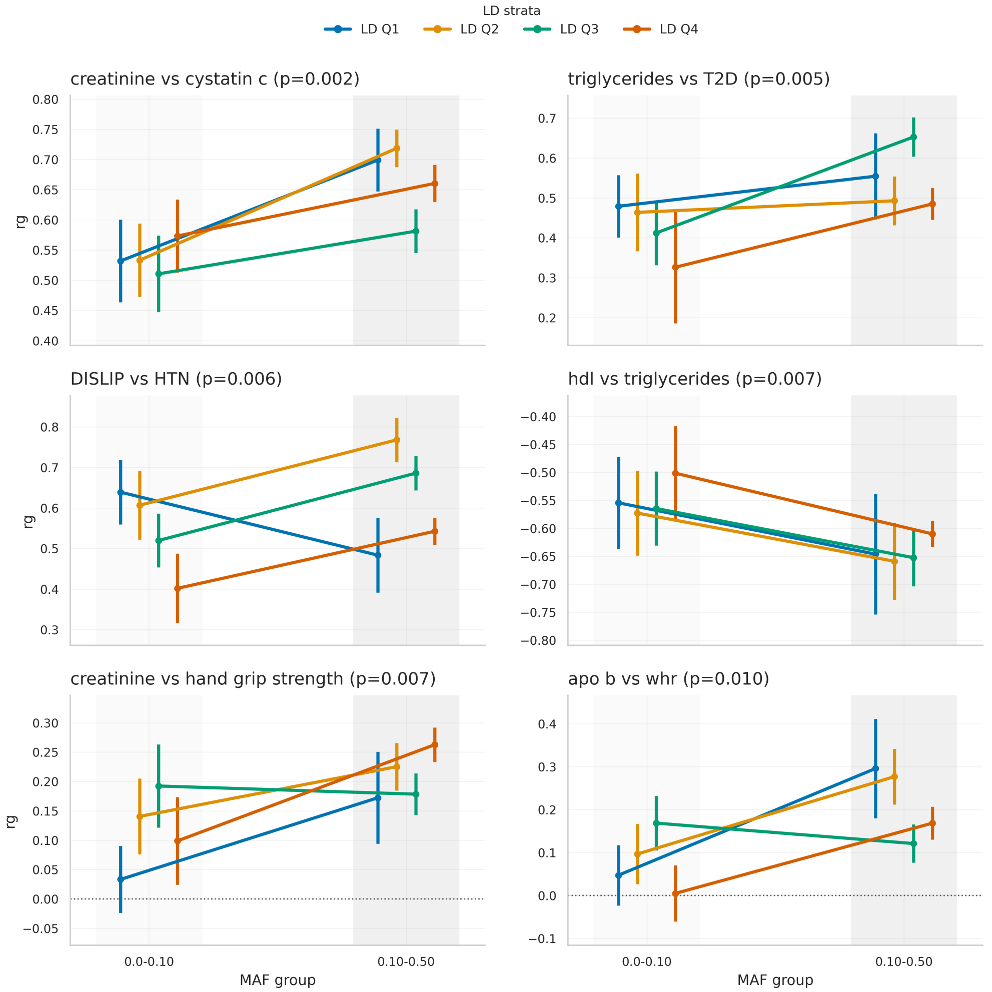

Figure S18: **Example trait pairs with evidence of MAF-dependent genetic correlation in EUR (300k).** Each panel shows estimates of $r_g$ for one trait pair stratified by MAF and LD. Points and lines correspond to LD quartiles, and vertical bars show SEs. We use the same 8 MAF–LD-bin model for estimation. The $p$ value in each panel title tests whether $r_g$ becomes larger in magnitude from low to high MAF, after multiplying the difference by the sign of the genome-wide genetic correlation. Specifically, for LD quartile $\ell$, define $\delta_\ell = \text{sign}(r_g^{\text{total}}) \left( r_{g,\ell}^{\text{high MAF}} - r_{g,\ell}^{\text{low MAF}} \right)$, where $r_g^{\text{total}}$ is the genome-wide genetic correlation for the trait pair, and $r_{g,\ell}^{\text{low MAF}}$ and $r_{g,\ell}^{\text{high MAF}}$ are the LD-quartile-specific estimates at low- and high-MAF bins. Let $\boldsymbol{\delta}$ be the vector of valid $\delta_\ell$ values across LD quartiles and let $\widehat{\Sigma}$ be their covariance matrix estimated from SNP-block jackknife replicates. The plotted $p$ value is from the one-sided test $H_0 : \mu \le 0$ vs. $H_A : \mu > 0$, where $\mu$ is the common mean of the sign-adjusted differences $\delta_\ell$ across LD quartiles. We estimate this mean using the jackknife covariance matrix: $\widehat{\mu} = \frac{\mathbf{1}^T \widehat{\Sigma}^{-1} \boldsymbol{\delta}}{\mathbf{1}^T \widehat{\Sigma}^{-1} \mathbf{1}}$, $\quad \text{SE}(\widehat{\mu}) = \left( \mathbf{1}^T \widehat{\Sigma}^{-1} \mathbf{1} \right)^{-1/2}$. The reported nominal $p$ value is the upper-tail normal probability for $Z = \widehat{\mu}/\text{SE}(\widehat{\mu})$.

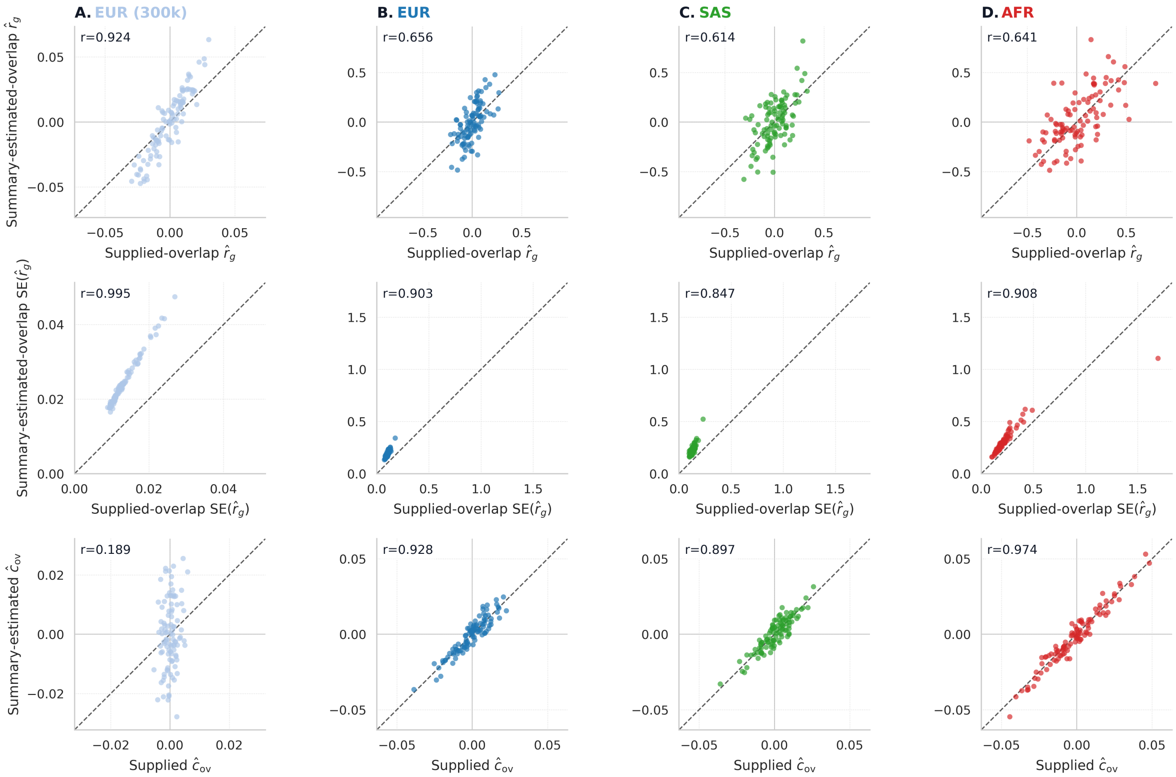

Figure S19: **SUM-CORE supplied-overlap and summary-estimated-overlap modes in single-component null simulations.** SUM-CORE was run either with the sample-overlap covariance supplied on the HE scale or with this quantity estimated from summary statistics in the single-component null benchmark ($h^2_{g,1} = 0.25$, $h^2_{g,2} = 0.60$, and $r_g = 0$). Each point is one simulation replicate and the dashed line marks equality. **Top row:** Point estimates of $r_g$. **Middle row:** Standard errors of $r_g$. **Bottom row:** Supplied and summary-estimated values of $\widehat{c}_{\text{ov}}$. The two modes produced broadly consistent point estimates, although estimates were more variable when $\widehat{c}_{\text{ov}}$ was estimated from summary statistics.

#### S3.4 Genome-wide LD score sensitivity and runtime scaling

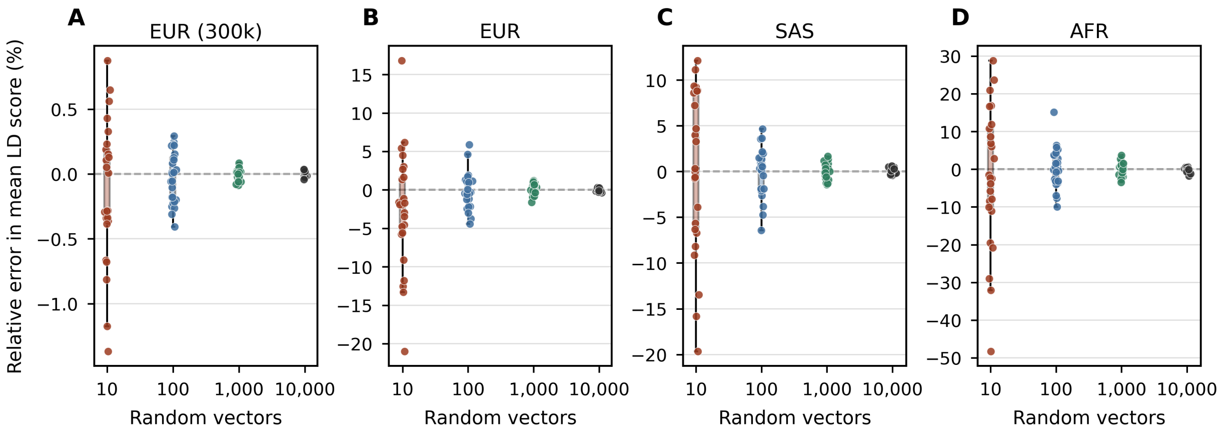

Figure S20: **Sensitivity of SUMMIT mean LD score estimates to number of random vectors.** Mean LD scores for each population as more random vectors $B$ were used to estimate the genome-wide LD scores (with genotype PC as covariates), compared with the SNP-wise average of the 25 $B = 10{,}000$ runs. We see that larger cohorts require less number of random vectors to get stable LD score estimates (e.g., EUR (300k) variance is less than 1% even with 10 random vectors), whereas small cohorts with long-range LD require many more to get accurate estimates (e.g., AFR requiring more than 1,000 random vectors to get within 5%).

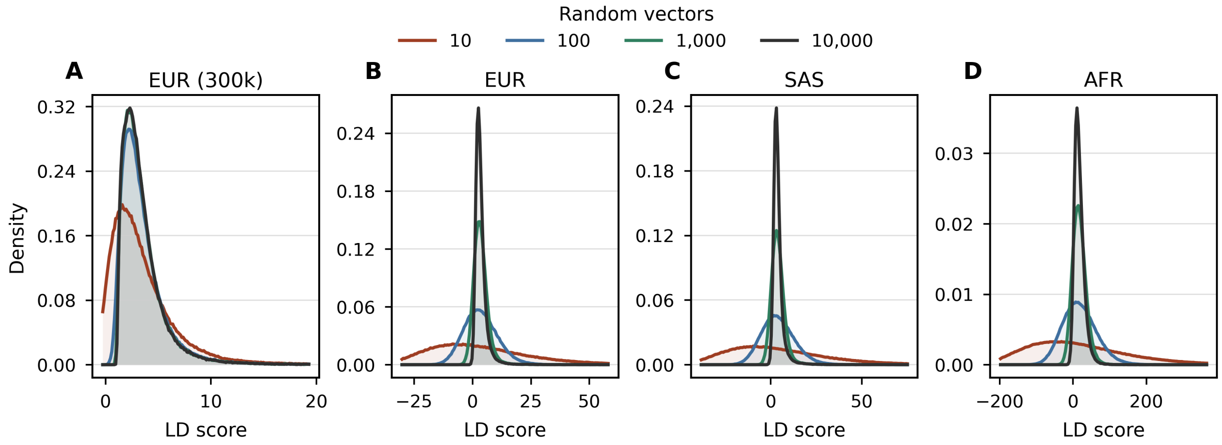

Figure S21: **Sensitivity of SNP-level genome-wide LD scores to the number of random vectors.** SNP-level genome-wide LD-score distributions are shown for each cohort as the number of random vectors $B$ increases. As with mean LD scores, in large cohort, SNP-level LD scores are stable after 1,000 random vectors. However in smaller cohorts, due to the stochasticity in LD score estimation, we see wide range of values (even negative) unless a significantly larger number of random vectors are used. We do note that SUMMIT normal equations only requires sum of LD scores unless variants are filtered out.

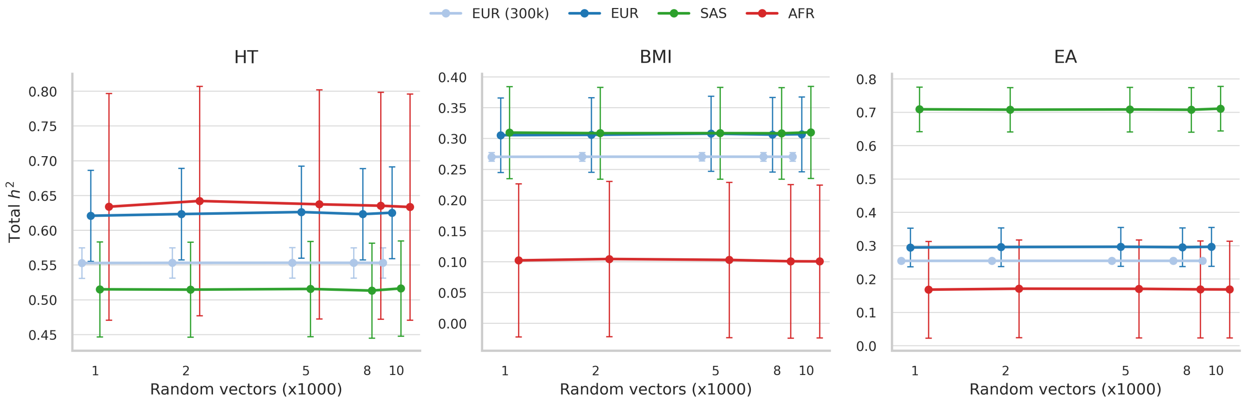

Figure S22: **Sensitivity of SUMMIT total $h^2$ estimates to number of random vectors.** Total $h^2$ estimates of height (HT), BMI, and educational qualifications (EA) as the number of random vectors for the LD scores were varied (2 MAF–4 LD annotation was used). We see that generally the estimates are extremely stable when a sufficiently large ($B = 1000$) random vectors were used, even in the small cohorts.

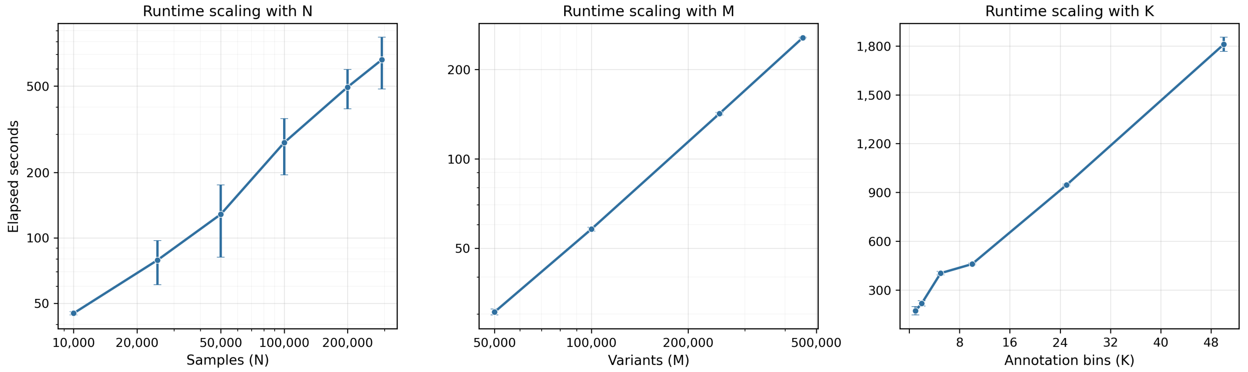

Figure S23: **Runtime scaling of Genome-wide LD score estimator.** We measure the runtime scaling of the estimator, as a function of sample size ($N$), number of variants ($M$), and number of bins ($K$), to validate the theoretical scaling of $\mathcal{O}\big(N_{\text{LD}}\, B \sum_k M_k\big)$. Scaling for $N$ and $M$ are shown in log-log scales, whereas $K$ is shown in linear-linear scales. We see that within reasonable variation, the scaling is linear for each variable.

##### S3.5 Real-data MAF–LD heritability and genetic architecture

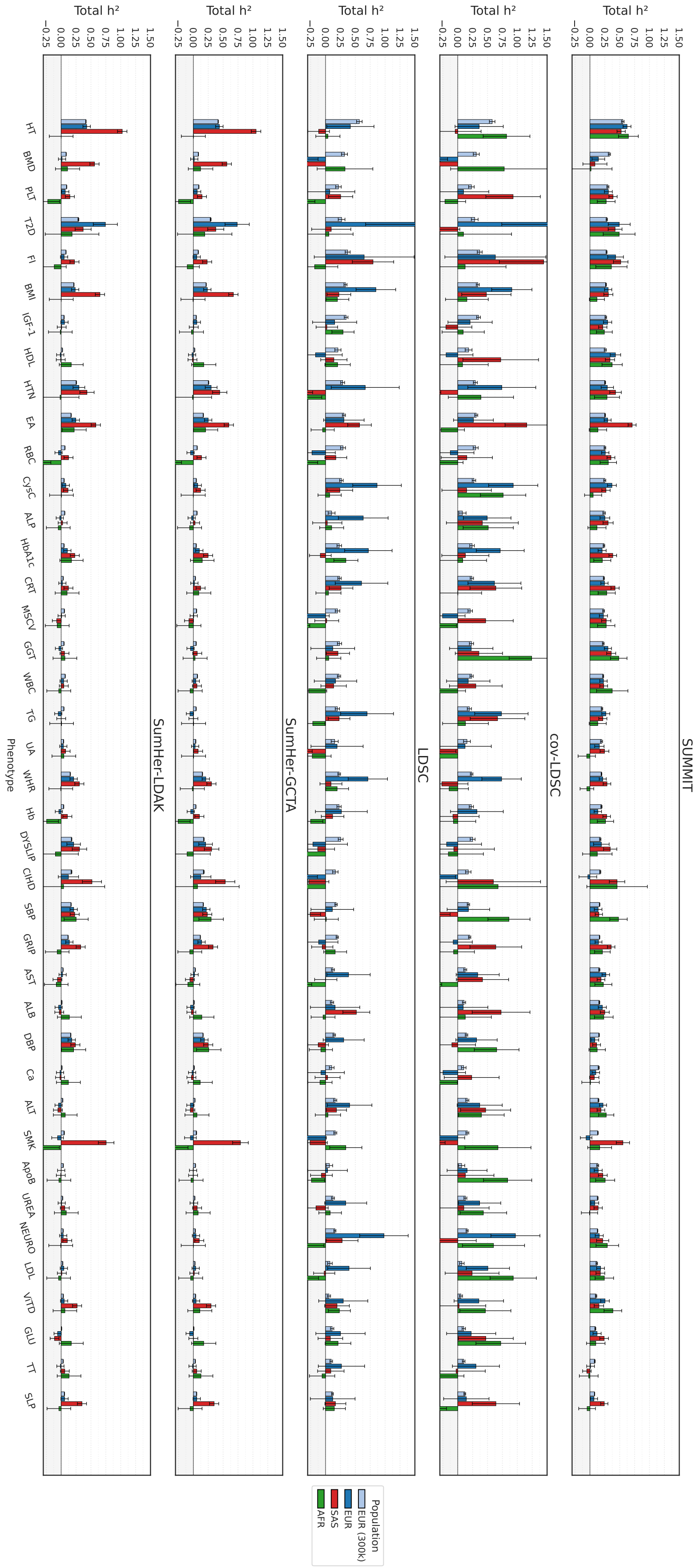

Figure S24: **Total heritability estimates with 24-bin MAF–LD partitioning across all 40 traits.**

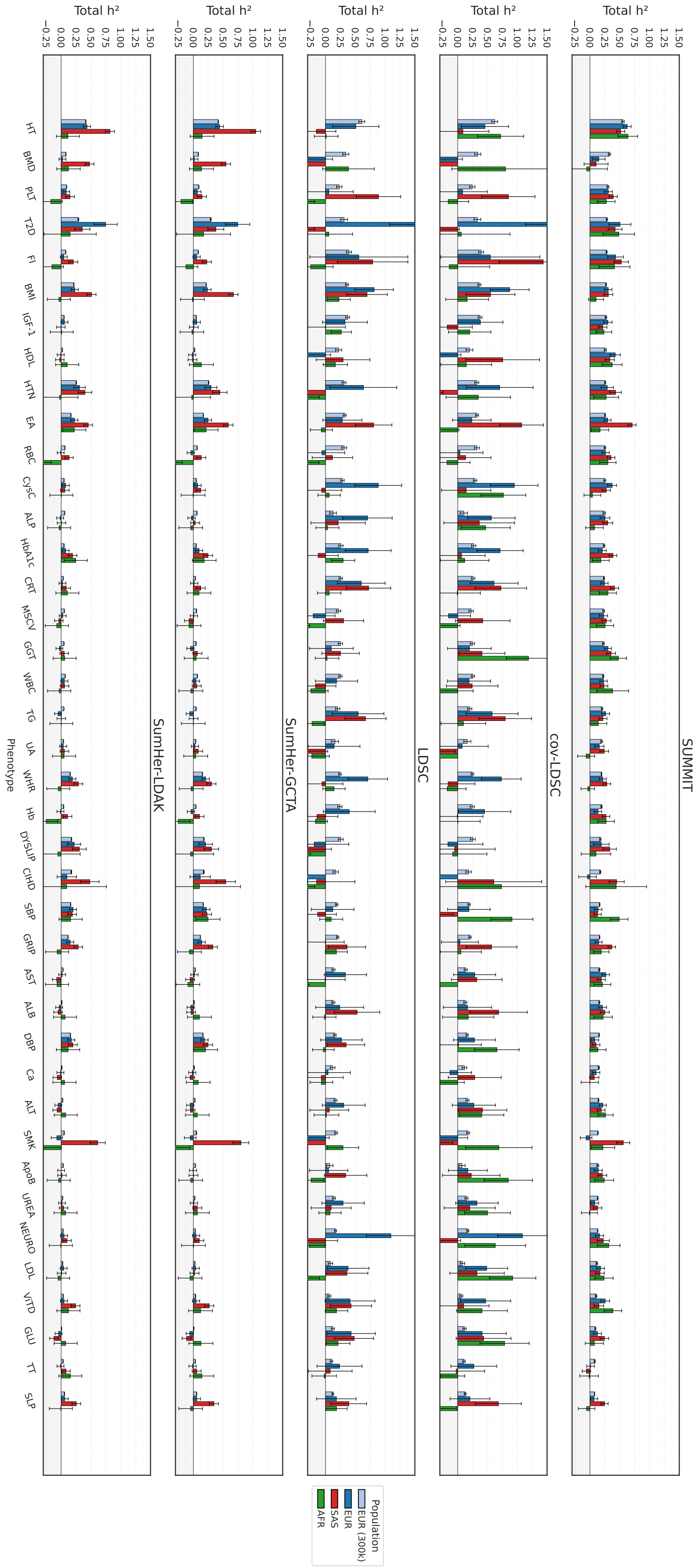

Figure S25: **Total heritability estimates with 8-bin MAF–LD partitioning across all 40 traits.**

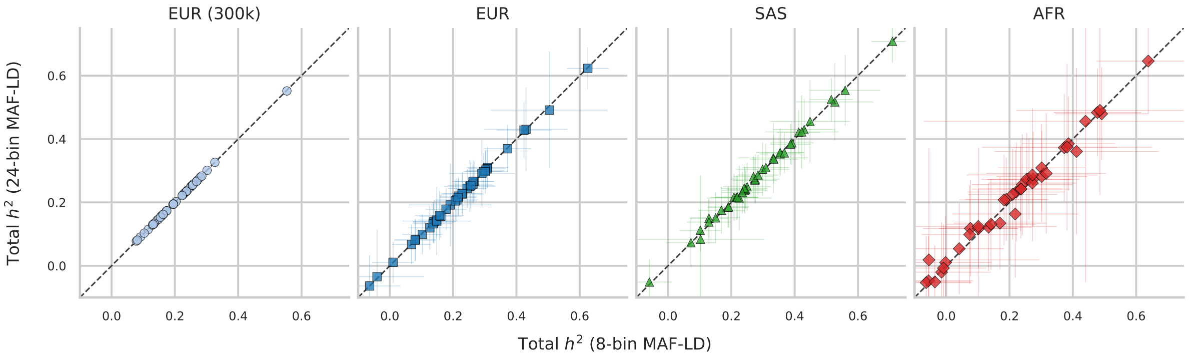

Figure S26: **Robustness of genome-wide heritability to MAF–LD annotation choice.** SUMMIT genome-wide $h^2$ estimates are compared between the 8-bin and 24-bin MAF–LD annotations across traits and cohorts. Correlation coefficients are 1.000 for EUR (300k), 1.000 for EUR, 0.999 for SAS and 0.992 for AFR.

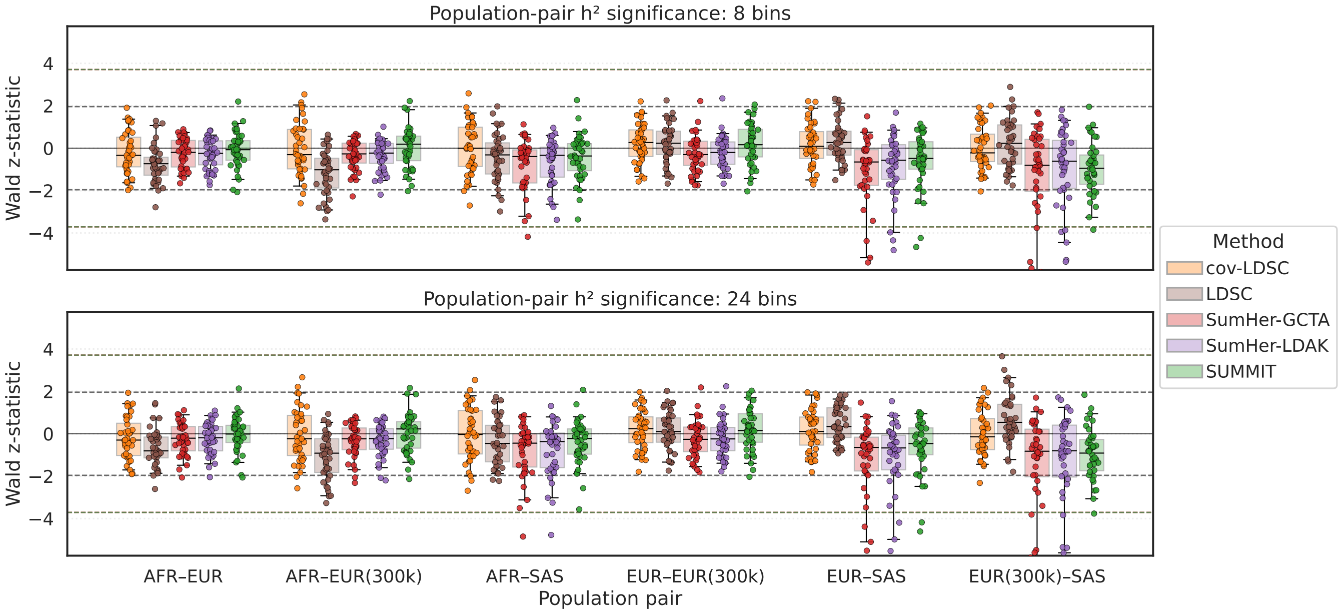

Figure S27: **Statistical test of population differences in genome-wide $h^2$ across all methods.**

#### S3.6 Baseline annotation partitioned heritability

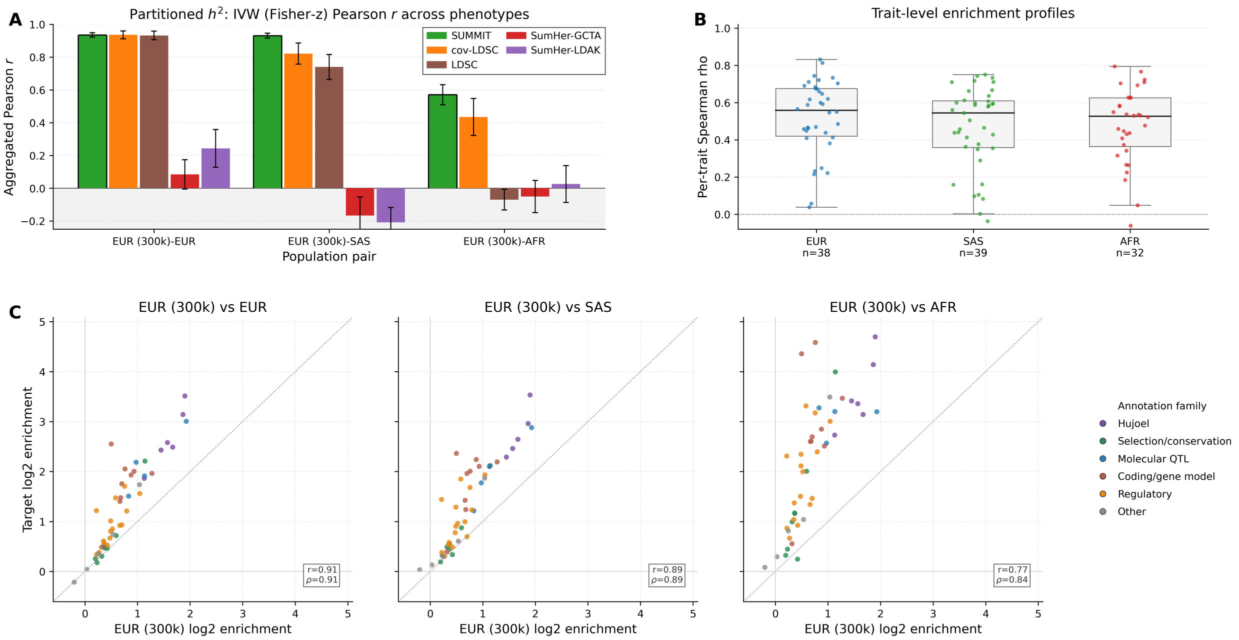

Figure S28: **Full comparison of Baseline-LD heritability and enrichment estimates across cohorts. Panel A:** Full method comparison for cross-cohort concordance of Baseline-LD annotation-specific $h_g^2$ profiles, using the same trait-level correlation and meta-analysis as in Figure 5. **Panel B:** Cross-cohort agreement in enrichment estimates for each trait, summarized as the Spearman correlation between each trait's annotation-enrichment estimates in EUR (300k) and the corresponding estimates in the target cohort. **Panel C:** Comparison of enrichment estimates across cohorts. Each point is one non-base, non-MAF Baseline-LD annotation, comparing $\log_2$ enrichment in EUR (300k) to the corresponding estimate in EUR, SAS or AFR. Enrichment was meta-analyzed across traits on the log scale, which requires restricting the meta-analysis to positive per-trait enrichment estimates. This positive-only restriction can shift log-scale enrichment estimates upward in lower-powered cohorts, especially AFR, due to large variance in enrichment (which can be negative). Annotation-enrichment rankings remain concordant across cohorts, although AFR estimates show greater dispersion.

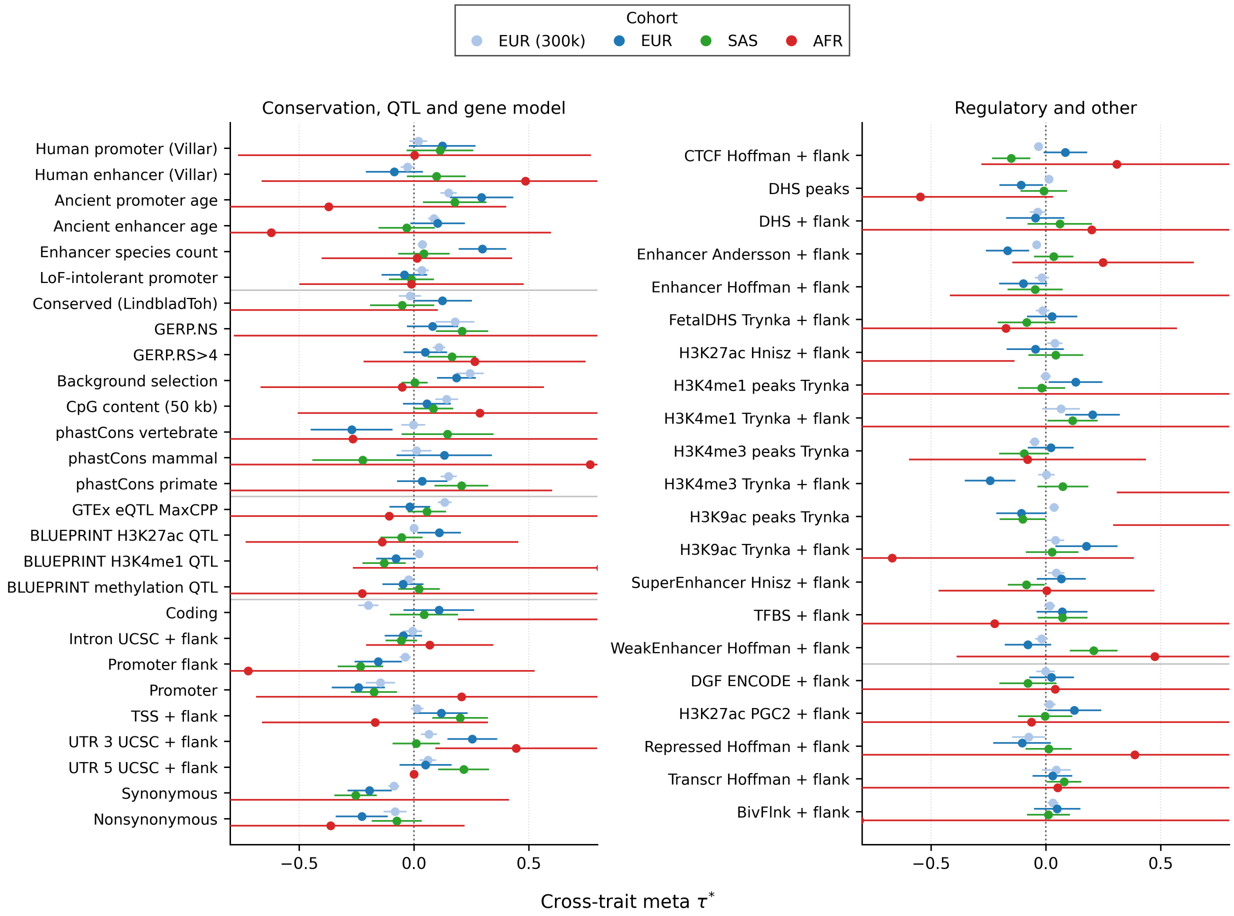

Figure S29: **Baseline annotation $\tau^*$ estimates across cohorts.** Cross-trait inverse-variance weighted meta-analyzed $\tau^*$ estimates are shown for all 48 non-base, non-MAF Baseline-LD annotations across cohorts. Annotations are split by annotation class for readability: conservation, QTL, and gene-model annotations are shown on the left, and regulatory and other annotations on the right. Points denote cohort-specific meta-analyzed estimates, and horizontal bars denote $\pm 1$ aggregate SE. Aggregate SEs were estimated by reconstructing trait-level $\tau^*$ values in each of the 22 SUMMIT leave-one-chromosome SNP jackknife replicates and recomputing the cross-trait IVW aggregate within each replicate. Bars are truncated at $\tau^* = \pm 0.8$ for display; all point estimates are within the plotted range. Pairwise cohort tests using these aggregate jackknife SEs did not identify any annotation-level difference passing BH FDR or Bonferroni correction at 0.05. The largest nominal shift was lower background-selection $\tau^*$ in AFR relative to EUR (300k), but this did not survive multiple-testing correction, supporting broad cross-cohort agreement in conditional annotation effects with greater AFR uncertainty.

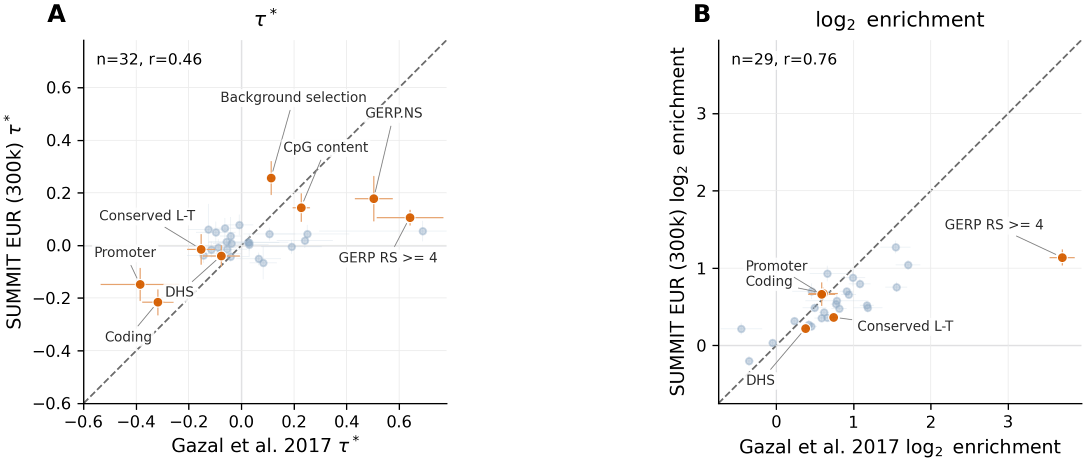

Figure S30: **SUMMIT estimates of functional enrichment compared with prior literature.** We compare our meta-analyzed estimates of $\tau^*$ (**Panel A**) and enrichment (**Panel B**) from the EUR (300k) array data against Gazal et al. (2017). Despite the difference in the annotation version (ours use Baseline v2.2., while the latter uses v1.1.), variant set, sample QC, and set of traits, we see that the estimates are generally concordant in magnitude and ordering. Error bars for our estimates are based on SNP-level block jackknife. We highlight several annotations that are biologically relevant (e.g., DHS) or focal in Gazal et al. (2017).

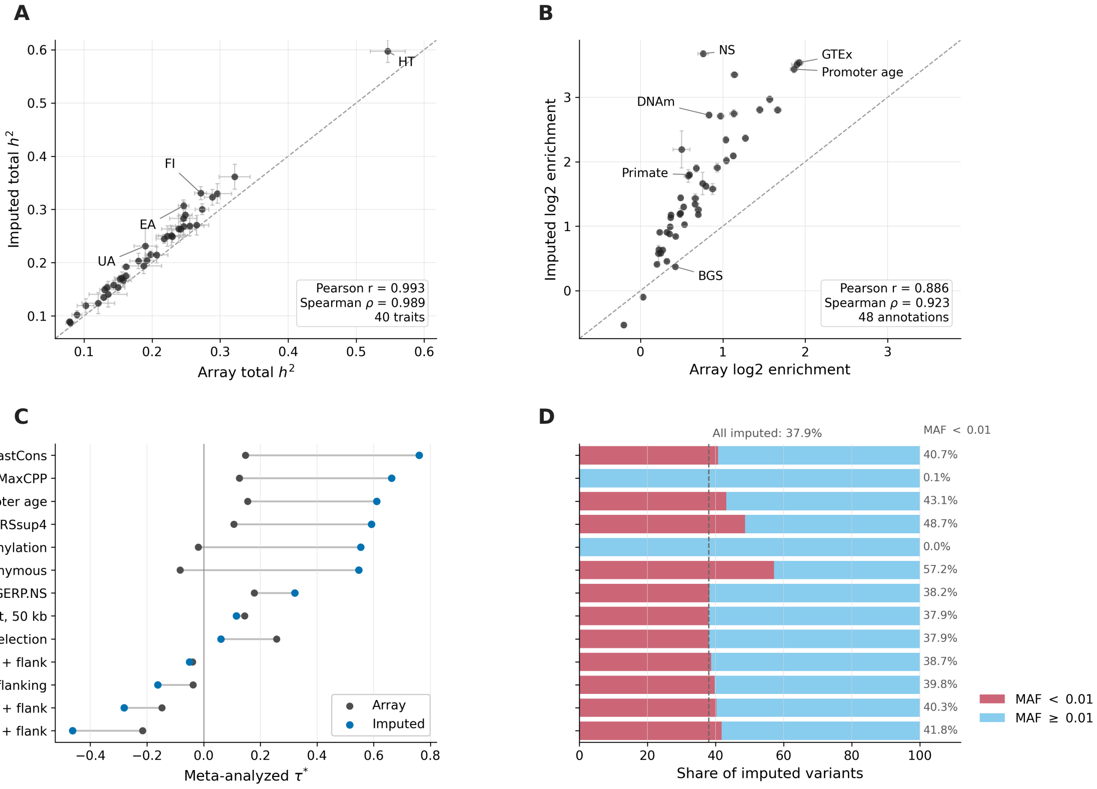

Figure S31: **Comparison of array and imputed panel for EUR (300k) in baseline annotation heritability results. Panel A:** Total $h^2$ estimates for 40 traits in the array and imputed analyses. Points are traits and error bars denote standard errors. Imputed genome-wide $h^2$ was larger across traits, while trait ordering was nearly unchanged. Traits with the biggest increase in heritability are annotated. **Panel B:** Meta-analyzed log2 enrichment for non-base, non-MAF annotations. Error bars denote standard errors on the log2-enrichment scale. Enrichment estimates were higher in the imputed analysis for many sparse functional annotations, but annotation rankings remained highly concordant. We annotate a few example bins in the figure. **Panel C:** Meta-analyzed $\tau^*$ for selected annotations, comparing array and imputed estimates. Conditional effects were more sensitive to genotype coverage than enrichment ranks. **Panel D:** Fraction of imputed variants with MAF $< 0.01$ for all imputed variants and selected annotations.

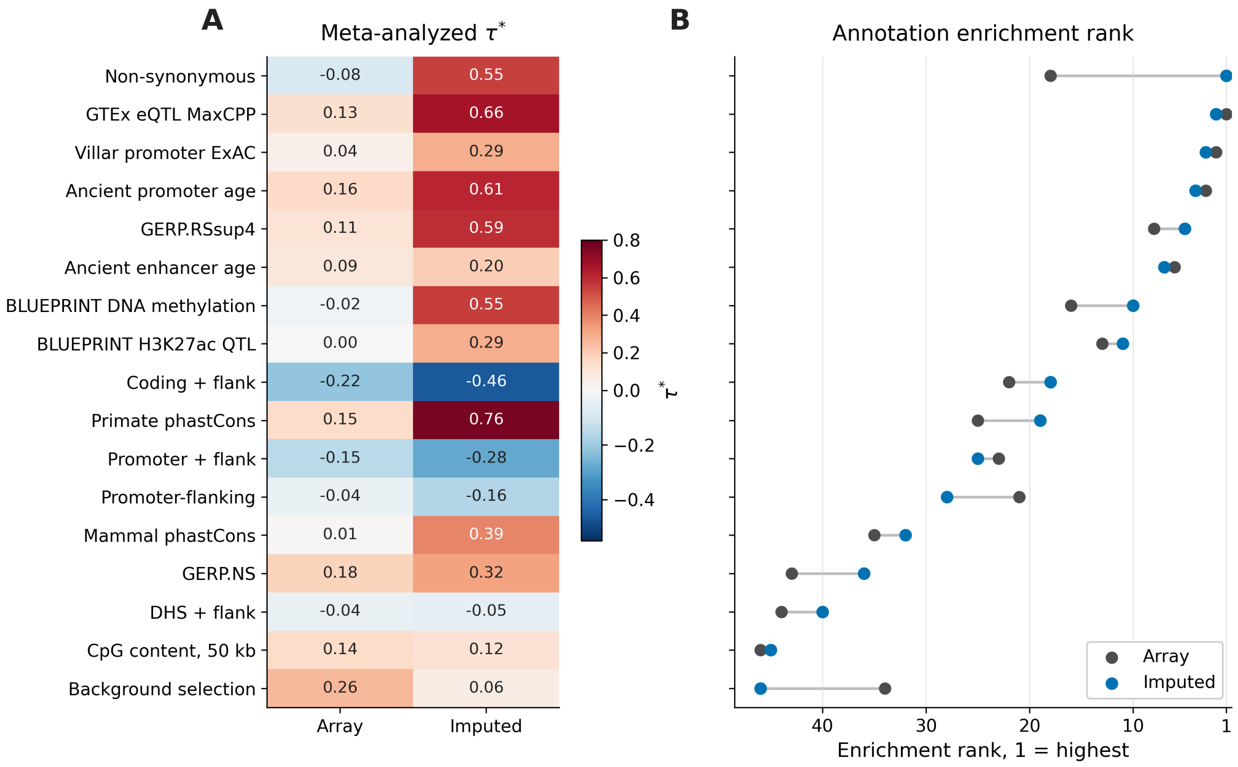

Figure S32: **Conditional effects and enrichment rankings for selected baseline annotations. Panel A:** Meta-analyzed $\tau^*$ estimates in the EUR (300k) array and imputed analyses. Rows are selected non-base, non-MAF annotations and are ordered by imputed enrichment rank in panel B. **Panel B:** Array and imputed enrichment ranks for the same annotations, where rank 1 denotes the highest enrichment among non-base, non-MAF annotations. Enrichment rankings were generally preserved across genotype panels, whereas the corresponding $\tau^*$ estimates showed larger annotation-specific shifts, including increased imputed estimates for constrained, molecular-QTL-linked and protein-altering annotations.

#### S3.7 Real-data genetic-correlation sensitivity analyses

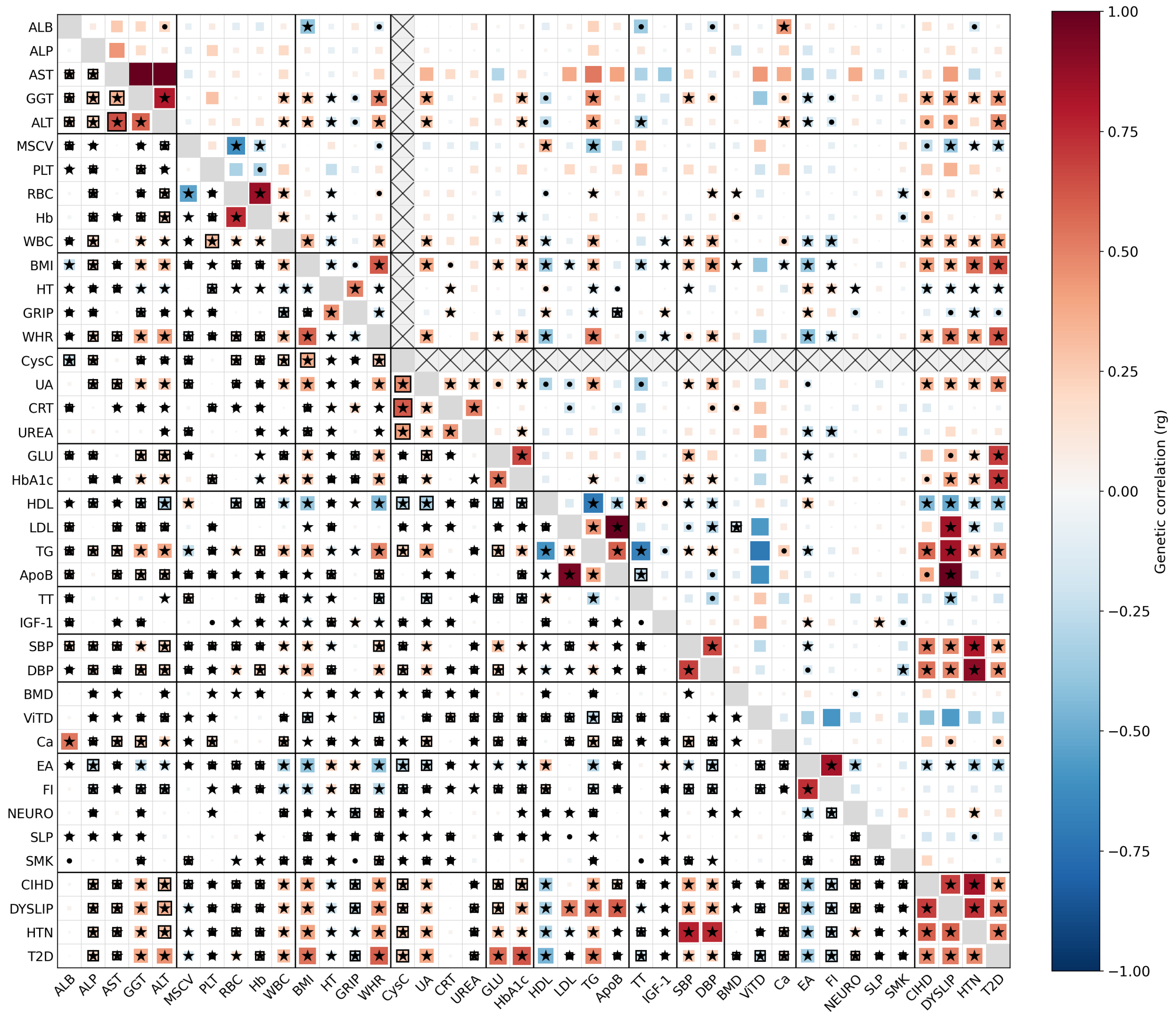

Figure S33: **Genetic correlation heatmap across all 40 traits, EUR (300k). Bottom left is SUMMIT estimates and top right is cov-LDSC estimates. Stars denote BH FDR-significant estimates (FDR <0.05) and boxes denote such hits that are not replicated in the other method.**

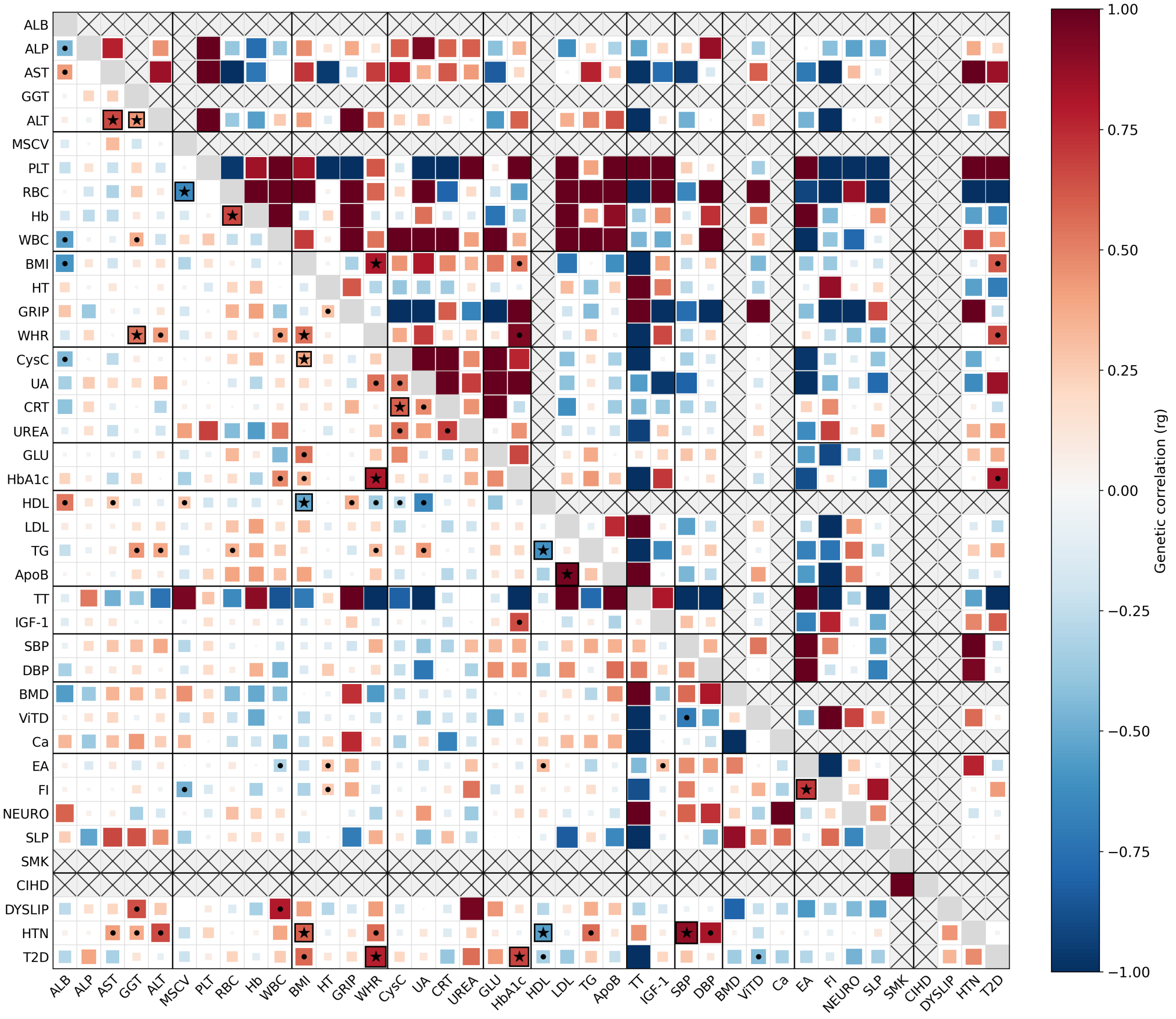

Figure S34: **Genetic correlation heatmap across all 40 traits, EUR. Bottom left is SUMMIT estimates and top right is cov-LDSC estimates. Stars denote BH FDR-significant estimates (FDR <0.05) and boxes denote such hits that are not replicated in the other method.**

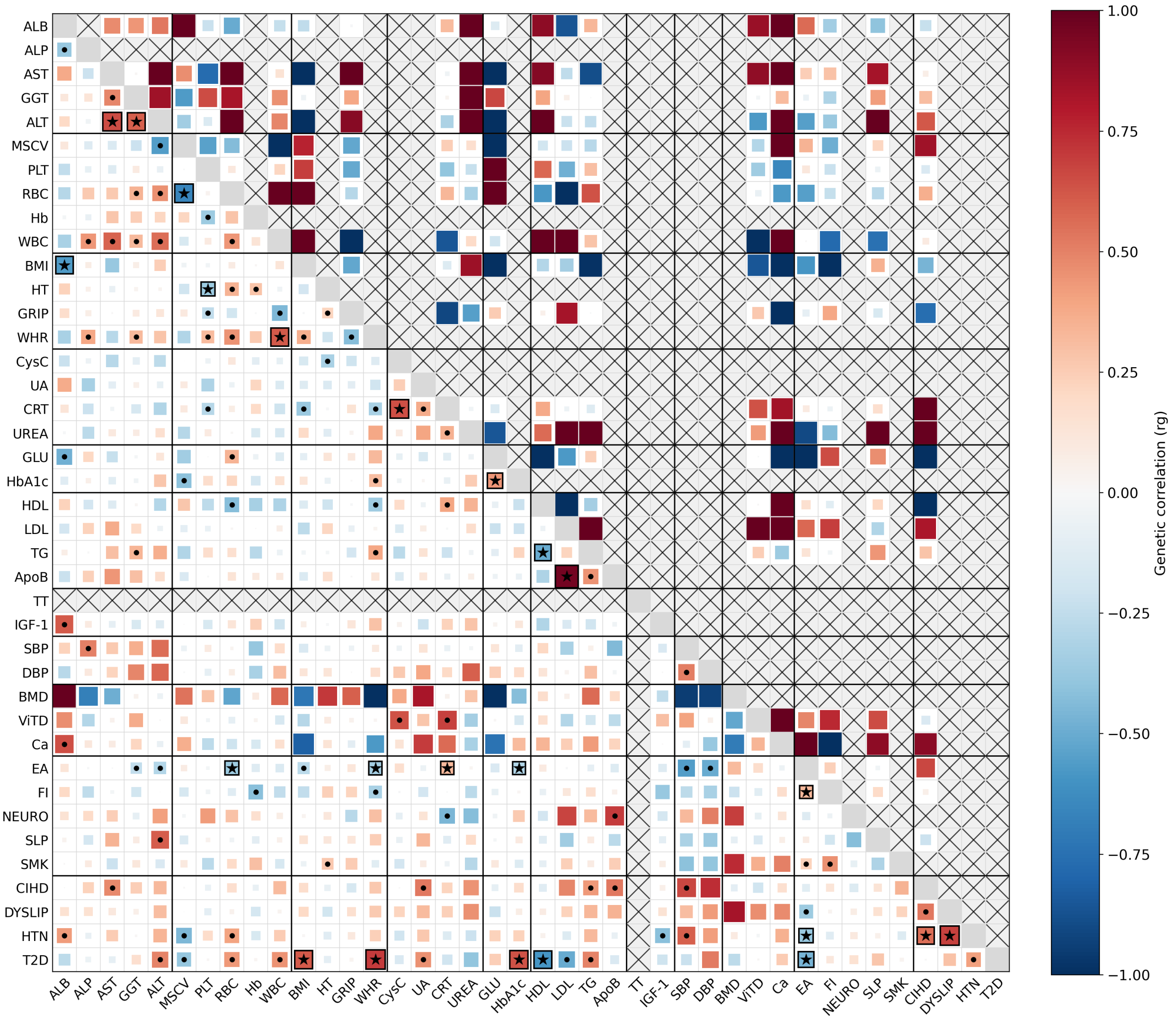

Figure S35: **Genetic correlation heatmap across all 40 traits, SAS. Bottom left is SUMMIT estimates and top right is cov-LDSC estimates. Stars denote BH FDR-significant estimates (FDR <0.05) and boxes denote such hits that are not replicated in the other method.**

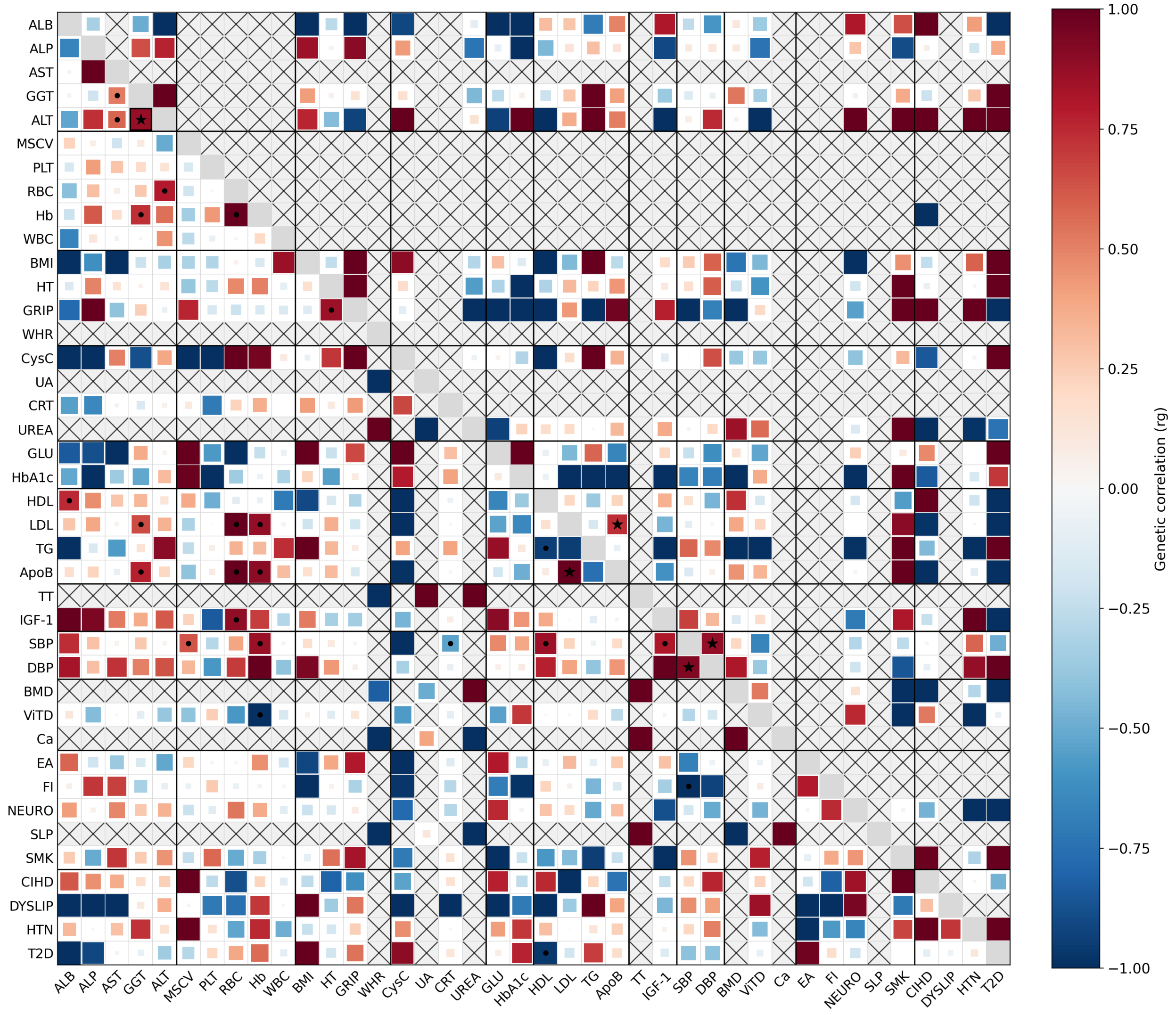

Figure S36: **Genetic correlation heatmap across all 40 traits, AFR. Bottom left is SUMMIT estimates and top right is cov-LDSC estimates. Stars denote BH FDR-significant estimates (FDR <0.05) and boxes denote such hits that are not replicated in the other method.**

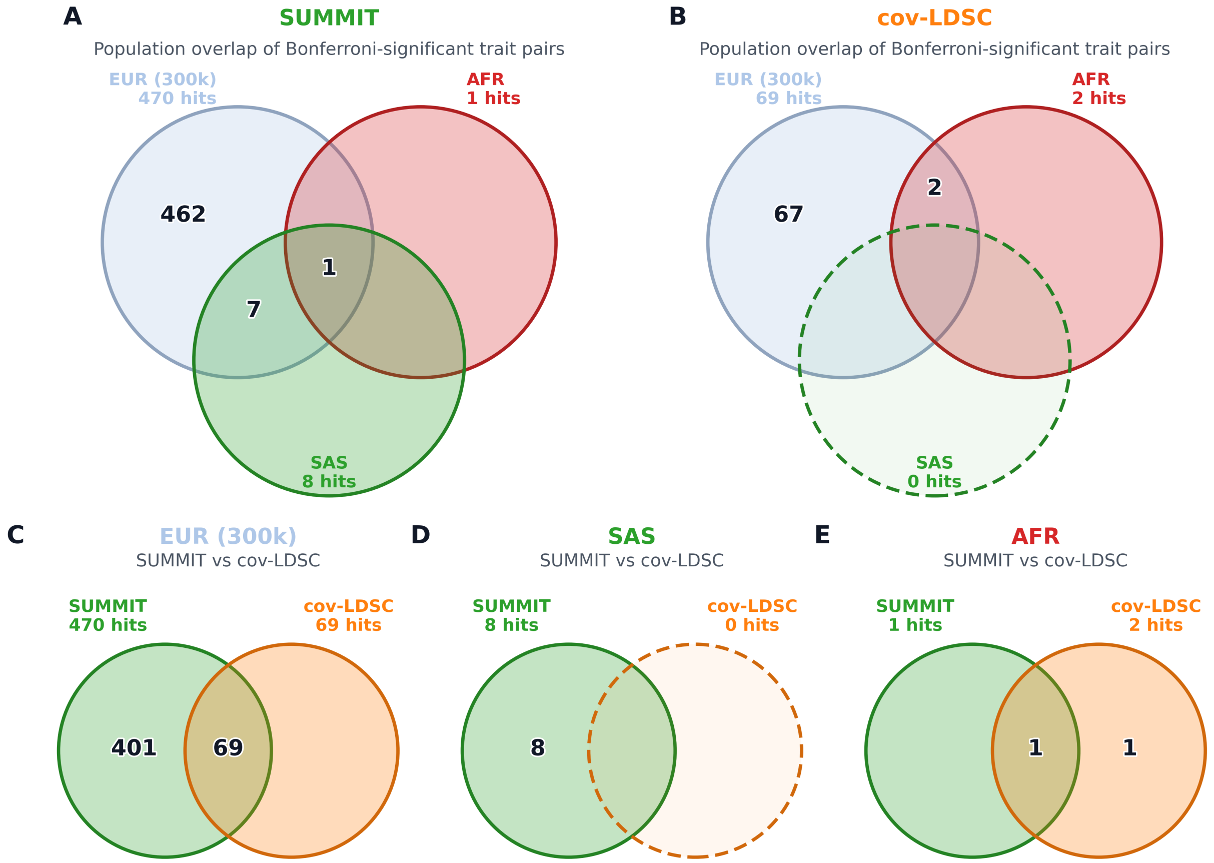

Figure S37: **Venn diagram of genome-wide Bonferroni-significant $r_g$ hits. Panels A, B:** We see that SUMMIT recovers many Bonferroni-significant hits for SAS and AFR that are also replicated in EUR (300k) whereas cov-LDSC only has 2 for AFR and none for SAS ($p = 0.05/m$, $m = 780$). **Panels C, D, E:** Across the populations, majority of the cov-LDSC hits are also replicated in SUMMIT.

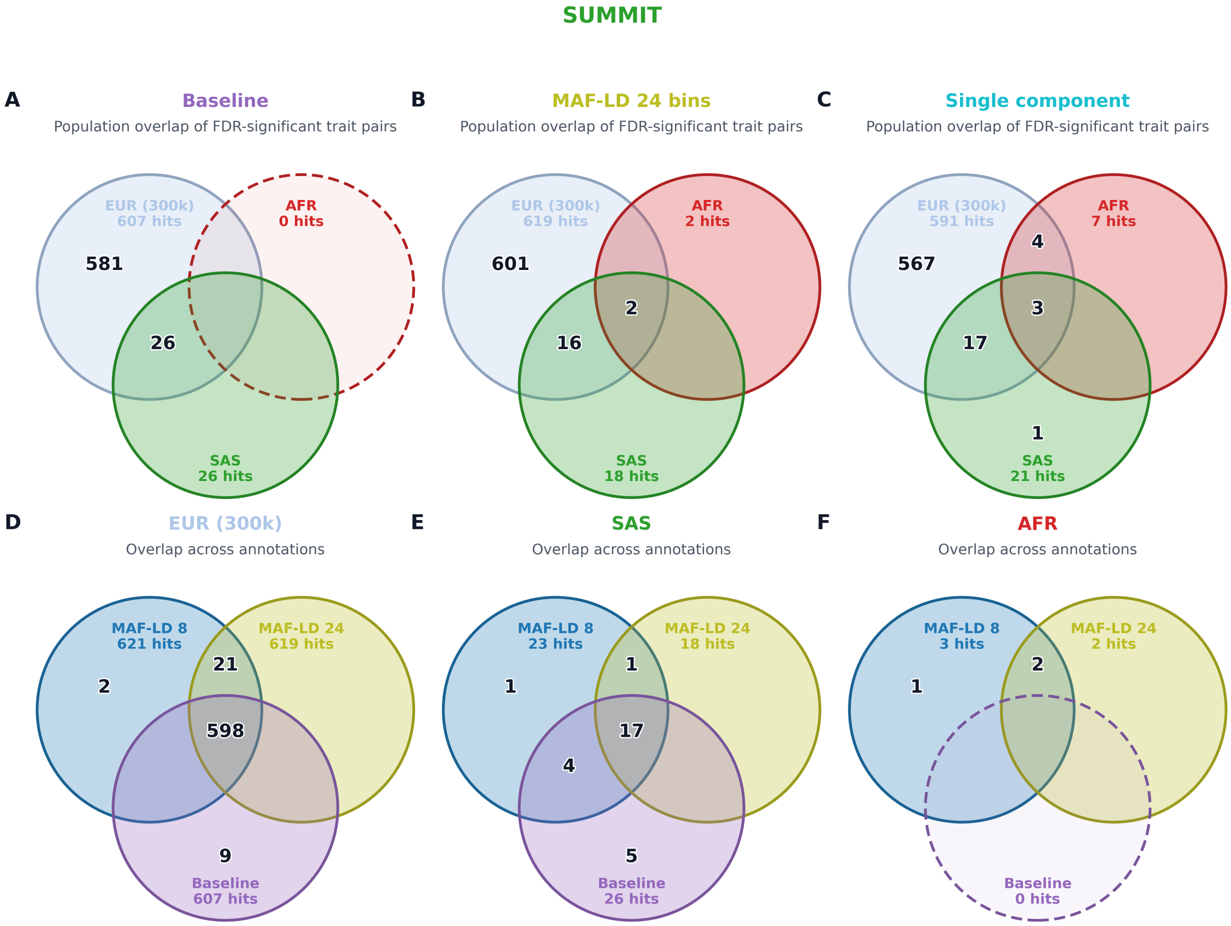

Figure S38: **Overlap of FDR-controlled total genetic-correlation hits across annotation models (SUMMIT)** FDR-significant hits are compared for each annotation across population (**top**) and across three partitioned annotations (**bottom**) for SUMMIT.

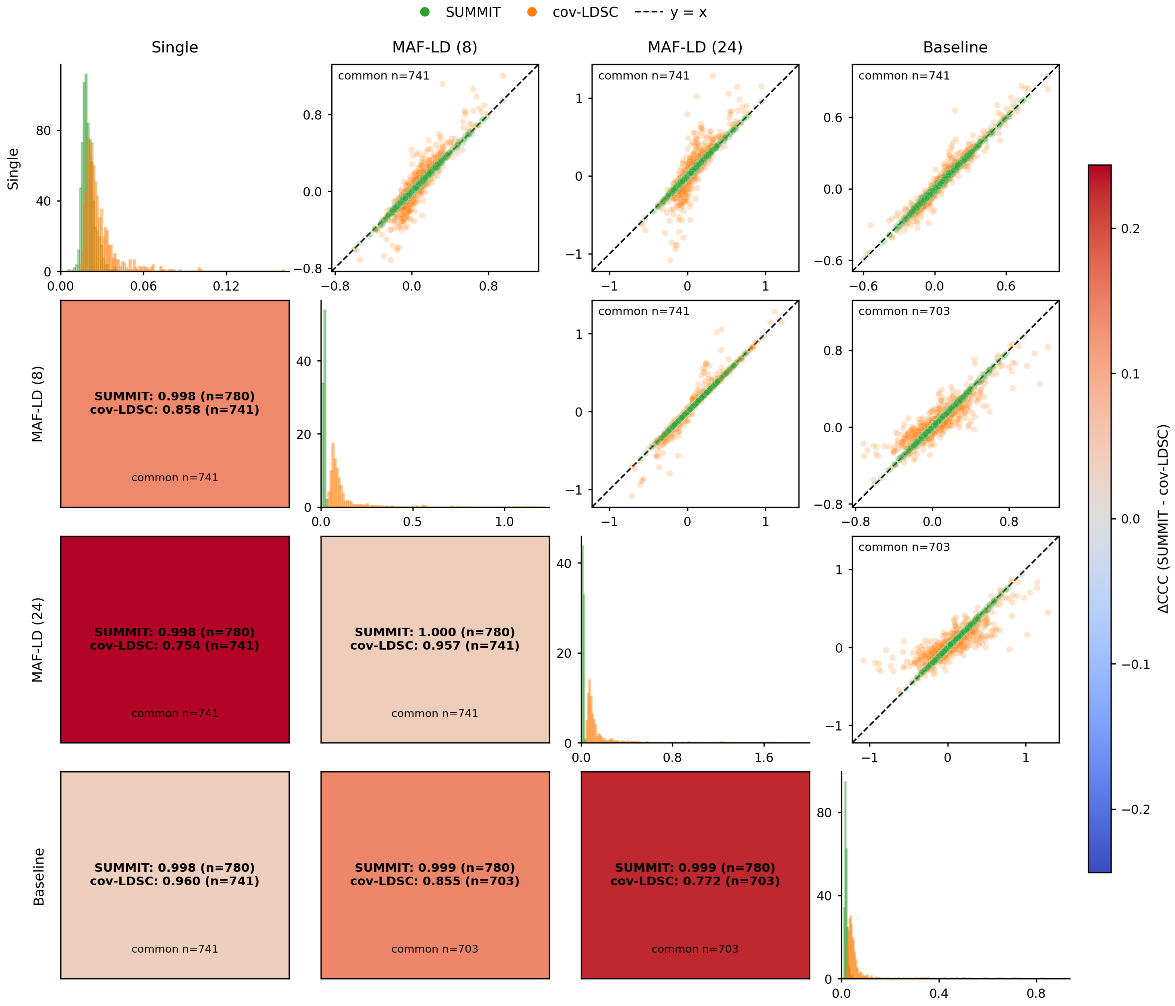

Figure S39: **Stability of genome-wide $r_g$ estimates across annotation choices, EUR (300k).** Rows and columns denote the annotation models used to estimate genome-wide genetic correlation. Upper-triangle panels compare pairwise $r_g$ estimates between annotation choices for SUMMIT and cov-LDSC; the dashed line marks equality, and $n$ gives the number of shared trait pairs plotted. Lower-triangle panels report the concordance correlation coefficient (CCC) between annotation-specific estimates within each method, with color indicating $\Delta\text{CCC} = \text{CCC}_{\text{SUMMIT}} - \text{CCC}_{\text{cov-LDSC}}$. Diagonal panels show the corresponding standard-error distributions. In this cohort, SUMMIT estimates are nearly invariant to the annotation model and have much tighter standard errors, while cov-LDSC shows visibly greater dispersion and lower cross-annotation concordance.

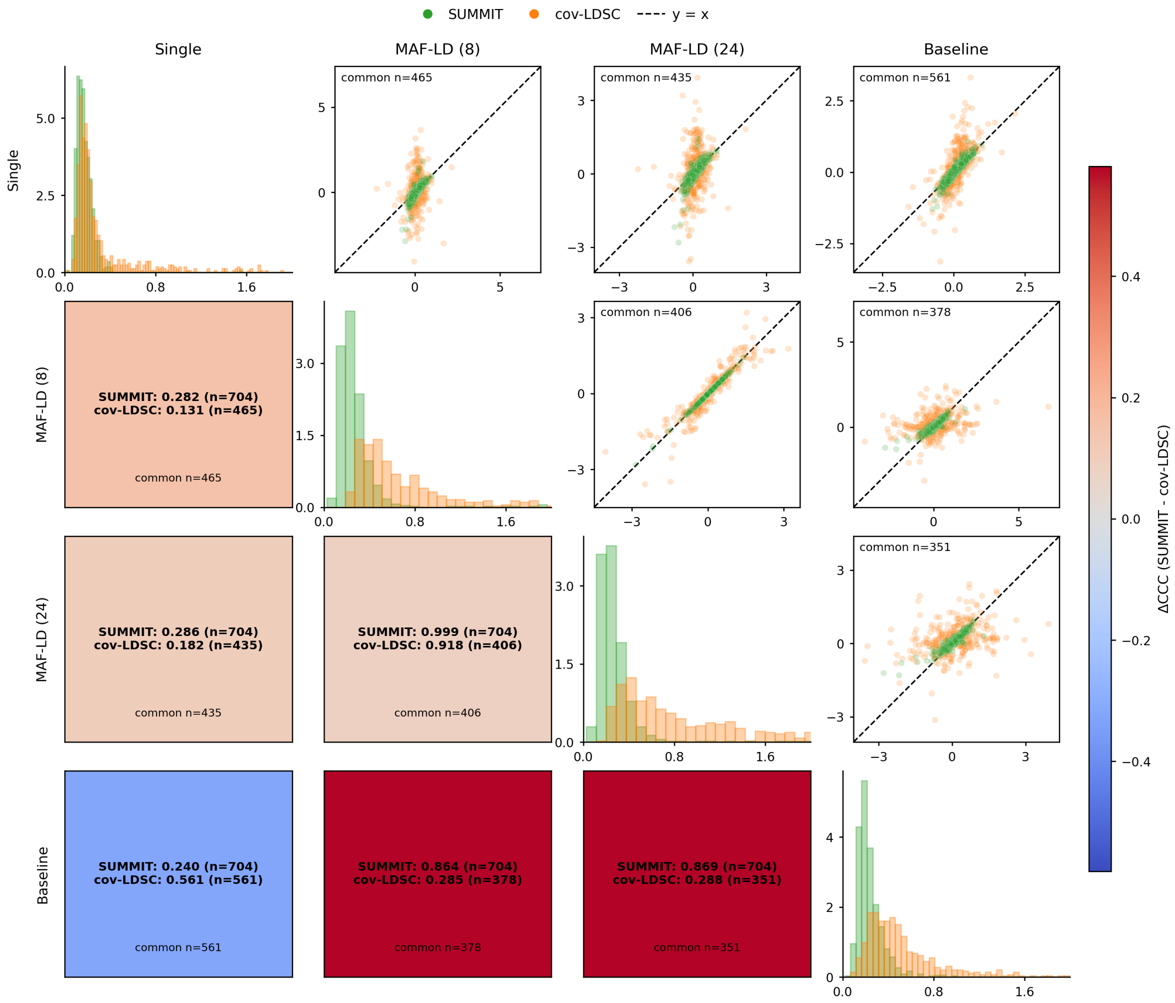

Figure S40: **Scatter plot of genome-wide $r_g$ stability across annotation choices, EUR.** In EUR, SUMMIT is highly concordant among the MAF-LD and baseline models but differs more from the single-annotation model; cov-LDSC shows broader uncertainty and lower concordance for most annotation comparisons.

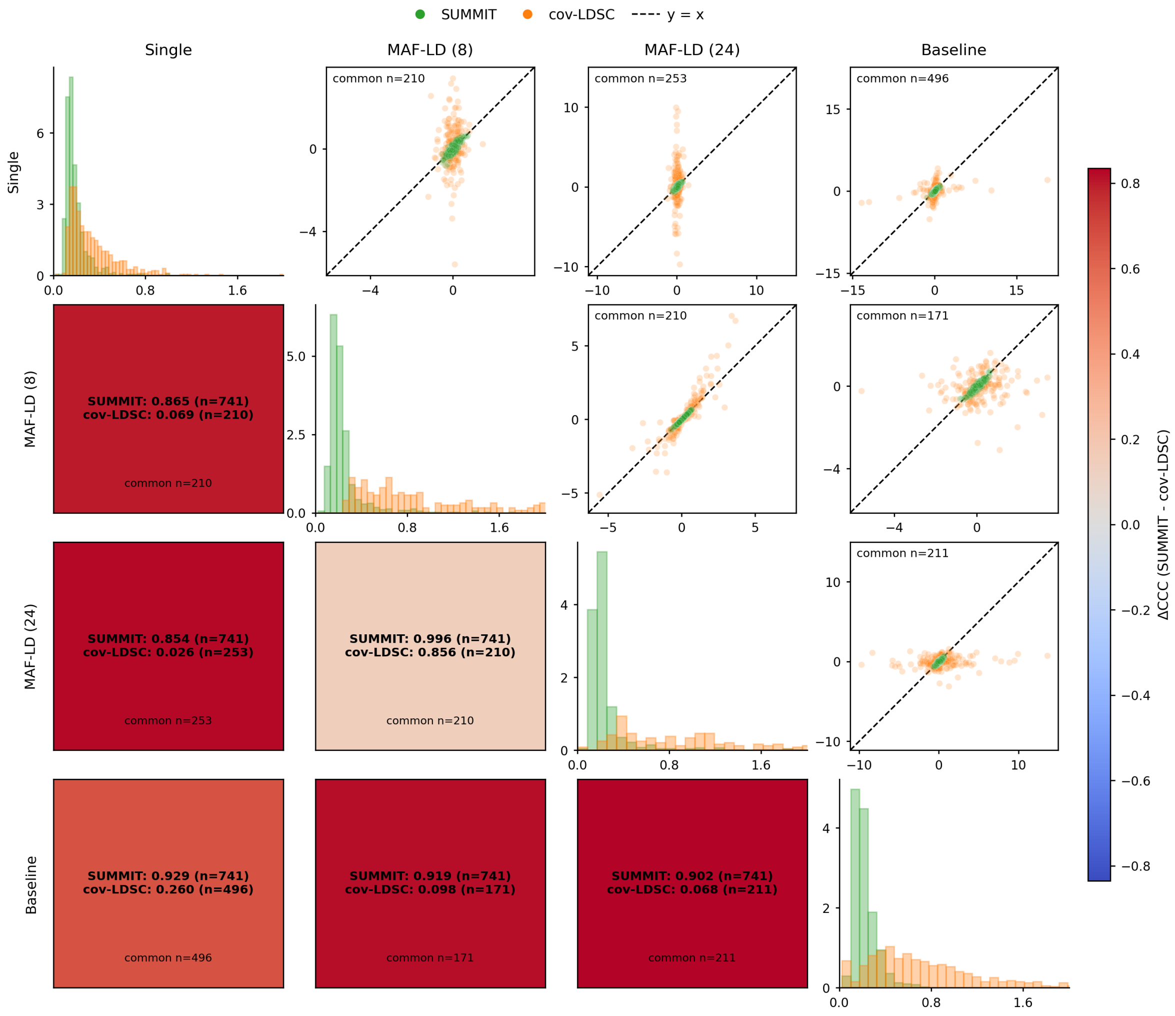

Figure S41: **Scatter plot of genome-wide $r_g$ stability across annotation choices, SAS.** In SAS, SUMMIT estimates are highly stable across annotation choices, whereas cov-LDSC shows marked annotation-dependent dispersion and substantially lower concordance.

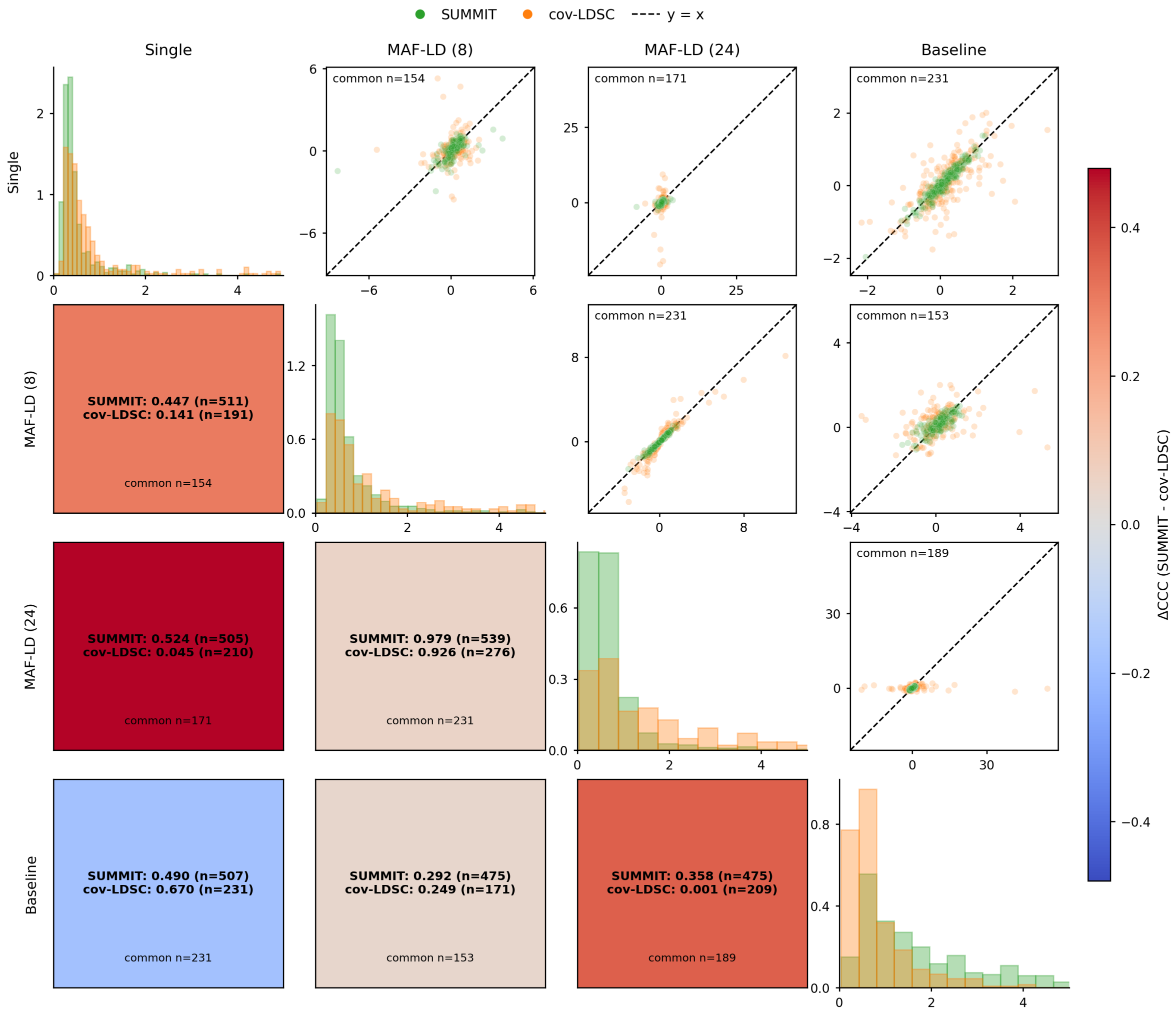

Figure S42: **Scatter plot of genome-wide $r_g$ stability across annotation choices, AFR.** In AFR, SUMMIT shows higher concordance for most annotation comparisons and smaller standard errors overall, although agreement is weaker than in the larger EUR (300k) cohort.

#### S3.8 External GWAS analyses with UKB LD scores

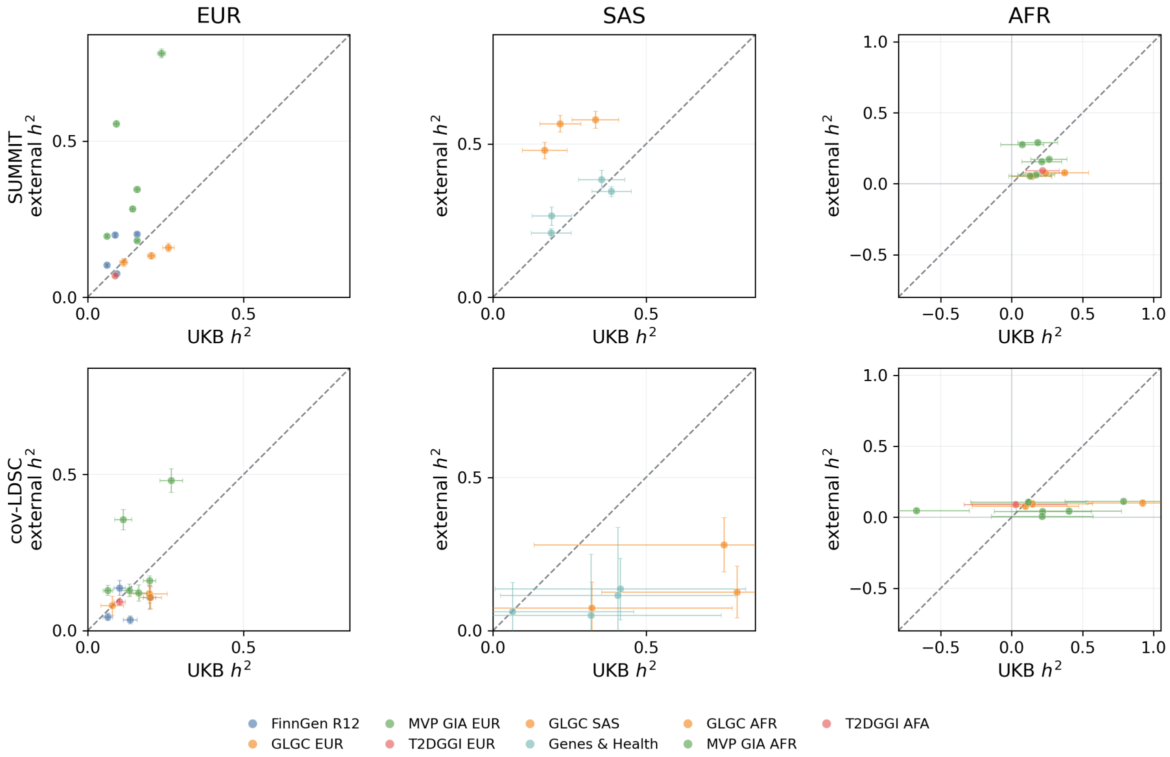

Figure S43: **External GWAS heritability estimates compared with UKB estimates.** We estimated SNP heritability from external GWAS summary statistics, then compared those estimates with corresponding UKB estimates for matched traits. Analyses are performed separately for each ancestry (EUR, SAS, and AFR) using ancestry-matched UKB LD scores and the MAF–LD 8-bin partition. Top panels show SUMMIT and bottom panels show cov-LDSC. Points denote matched external-source traits, colored by external source; horizontal and vertical bars denote standard errors for UKB and external estimates, respectively. Dashed lines denote $y = x$. Finite paired counts were 14, 7 and 10 in EUR, SAS and AFR, respectively, for both methods. Median external-to-UKB $h_g^2$ ratios were 1.47, 1.39 and 0.44 for SUMMIT, and 0.86, 0.28 and 0.16 for cov-LDSC in EUR, SAS and AFR, respectively. Median external-GWAS SEs for SUMMIT versus cov-LDSC were 0.008 versus 0.025 in EUR, 0.028 versus 0.096 in SAS, and 0.008 versus 0.017 in AFR. The largest EUR discrepancies were driven primarily by MVP GIA EUR traits, including HbA1c, glucose, AST, ALT and ischemic heart disease; the four quantitative laboratory traits also had inflated cov-LDSC intercepts (1.60, 1.50, 1.49 and 1.38 for HbA1c, glucose, AST and ALT, respectively; one-sided tests of intercept $> 1$, all $p < 10^{-12}$). We see that absolute $h_g^2$ estimates are not consistently calibrated across external GWAS and UKB under reference mismatch and differences in GWAS scale.

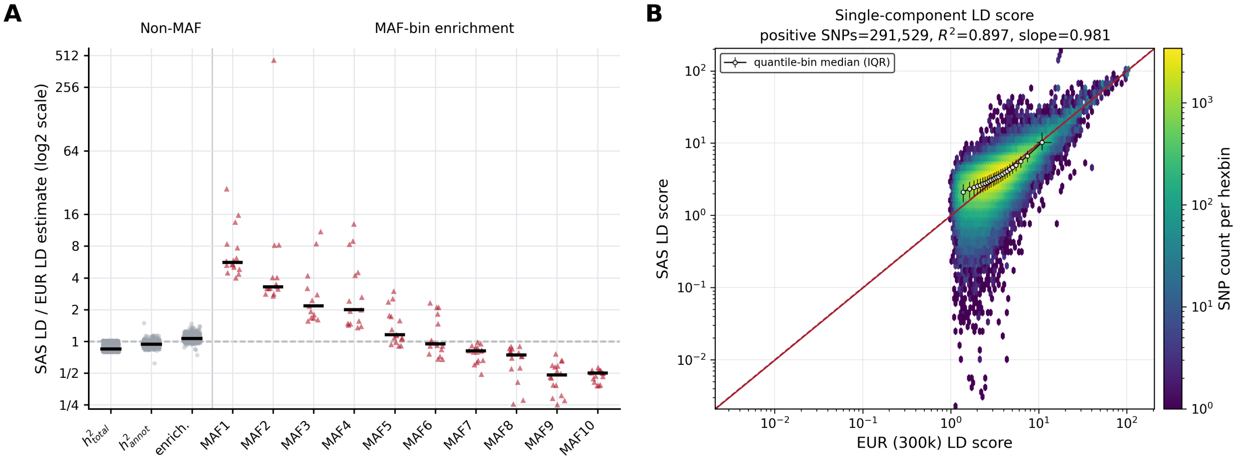

Figure S44: **Additional diagnostics of enrichment robustness under LD-reference mismatch.** SUMMIT was run on the same 16 external EUR GWAS used in Figure 8, analyzing each trait twice with either the EUR (300k) or deliberately mismatched SAS LD and annotation reference. Both analyses used the Baseline-LD v2.2 annotation model and the same EUR–SAS common SNP set (291,793 variants). **Panel A:** Ratios of estimates using the SAS reference to estimates using the EUR reference, shown on a $\log_2$ scale. For non-MAF annotations, total heritability estimates were lower with the SAS reference (median ratio = 0.849), annotation-specific heritability was less attenuated (median ratio = 0.942), and enrichment remained comparatively stable (median ratio = 1.065; 5th–95th percentile = 0.982–1.204). In contrast, MAF-bin enrichment was strongly distorted and changed direction across MAF strata, with median ratios ranging from 5.59 and 3.28 in the two lowest-MAF bins to 0.48 and 0.50 in the two highest-MAF bins. Black horizontal bars denote medians. **Panel B:** SNP-level single-component LD scores computed using the EUR (300k) and SAS references. The log-scale hexbin density includes 291,529 SNPs with positive LD scores in both references. Black points and error bars denote quantile-bin medians and interquartile ranges. Individual SNP LD scores differed between references, although the binned trend and fit through the origin remained close to a shared scale ($R^2 = 0.897$, slope = 0.981). The dashed line marks equality.
